# Sterile spikelets assimilate carbon in sorghum and related grasses

**DOI:** 10.1101/396473

**Authors:** Taylor AuBuchon-Elder, Viktoriya Coneva, David M. Goad, Doug K. Allen, Elizabeth A. Kellogg

## Abstract

Sorghum and its relatives in the grass tribe Andropogoneae bear their flowers in pairs of spikelets, in which one spikelet (seed-bearing, or SS) of the pair produces a seed and the other is sterile or male (staminate). This division of function does not occur in other major cereals such as wheat or rice. Additionally, one bract of the seed-bearing spikelet often produces a long extension, the awn, which is in the same position as but independently derived from that of wheat and rice. The function of the sterile spikelet is unknown and that of the awn has not been tested in Andropogoneae. We used radioactive and stable isotopes of carbon, as well as RNA-seq of metabolically important enzymes to show that the sterile spikelet assimilates carbon, which is translocated to the largely heterotrophic SS, thereby functioning as a nurse tissue. The awn shows no evidence of photosynthesis. These results apply to distantly related species of Andropogoneae. Thus, the sterile spikelet, but not the awn, could affect yield in the cultivated species and fitness in the wild ones.

---

The grain and bioenergy crop sorghum (*Sorghum bicolor*), like most of its 1200 relatives in the grass tribe Andropogoneae, produces sterile flowers in its inflorescence, with more sterile flowers than fertile ones. This pattern is counter-intuitive, in that the plant appears to be sacrificing potential reproductive output by taking many of its floral structures “off line.” The sterile structures must therefore have another function to compensate for the loss of reproductive potential.

The seed of sorghum, like that of all grasses, is enclosed in floral bracts (glumes and lemmas), which together form a terminal unit known as a spikelet (a little spike)^1^. The seed-bearing, or sessile, spikelet (SS) sits directly on the inflorescence axis and is paired with a pedicellate (stalked) spikelet (PS)(Fig. 1). Unlike the seed-bearing SS, which is bisexual, the PS is generally sterile in sorghum, although in some lines the PS produces stamens. Presence of a PS, its shape, size and sex expression appear to be genetically fixed among plants of any given accession.

**Figure 1.**
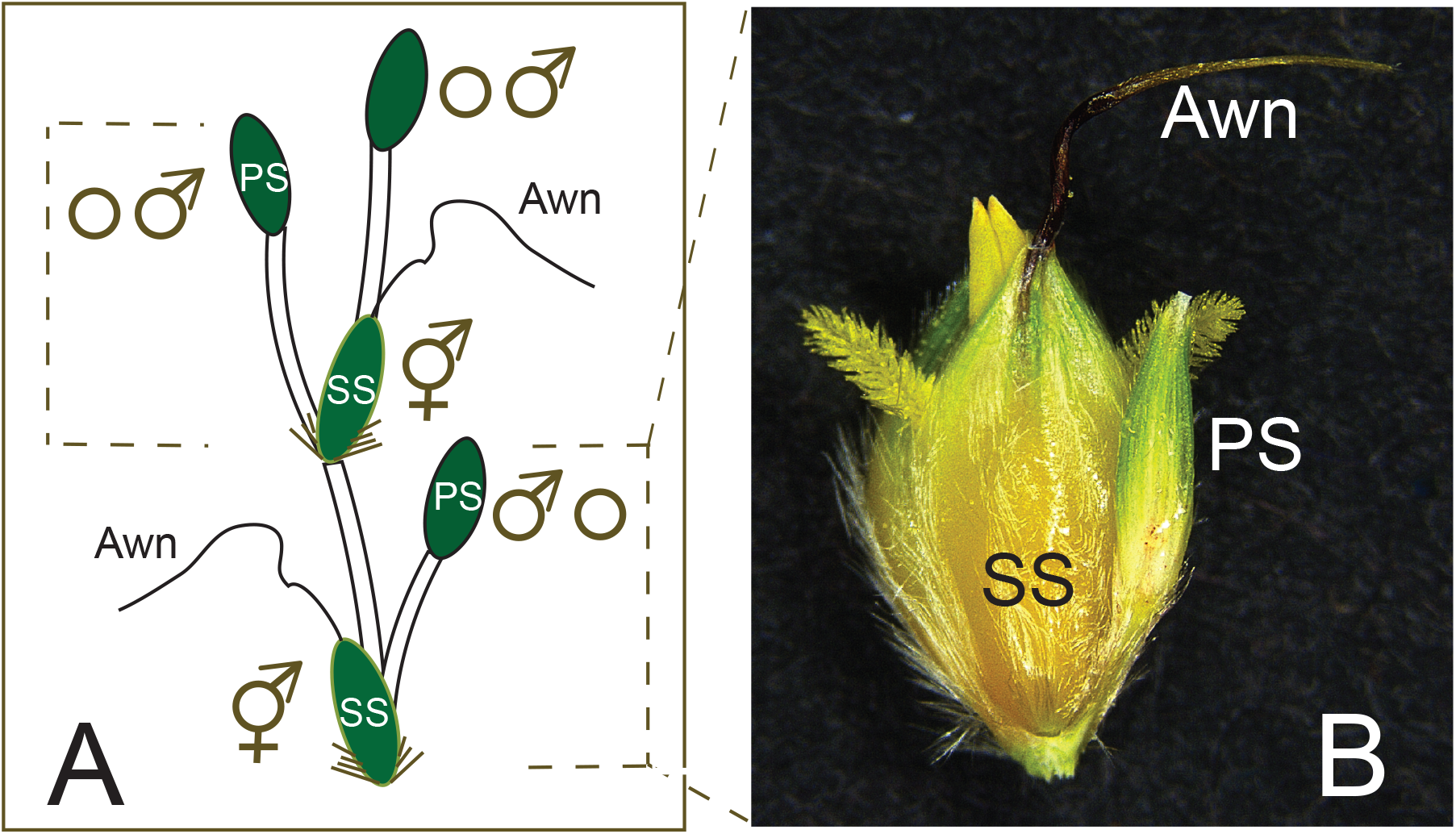
Spikelet pair structure in sorghum. A. Illustration of two spikelet pairs, marked by dotted lines, plus a terminal spikelet, which is morphologically identical to a pedicellate spikelet. Each pair is composed of a sessile spikelet, which includes a bisexual flower and bears the seed, and a pedicellate spikelet, which may be either sterile (most commonly) or staminate. The sessile spikelet bears a twisted awn from the lemma (floral bract). B. Spikelet pair of sorghum accession SAP-15 (PI 656014). SS, sessile spikelet; PS, pedicellate spikelet. Scale bar = 1 mm.

The function of the PS is unknown in sorghum or in any other member of Andropogoneae, in most of which the PS is also sterile or staminate^1^. Because grasses are wind-pollinated, the PS cannot function in pollinator attraction, as sterile flowers do in some other angiosperms. In some wild species, the PS may function in seed dispersal and/or in controlling pollen to ovule ratio. However, the dispersal function is not universal, nor is it relevant in cultivated sorghum, in which the grains are not shed, and the pollen production is only relevant in a few accessions.

The spikelet pair in sorghum may also include an awn on the SS (Fig. 1). The awn, when present, is a slender extension of the lemma (floral bract) of the upper flower, and may be twisted, nearly straight, and/or geniculate. Like the PS, the presence, shape and size of the awn appear to be genetically fixed within an accession and its function has not been demonstrated. In some wild Andropogoneae, the awn is hygroscopic and can help orient the spikelet in the soil to enhance germination^2–5^. Awns in wheat and rice are vascularized and assimilate carbon, which is then transferred to the grain^6–8^, often contributing appreciably to yield; however, wheat and rice have solitary (unpaired) spikelets. Carbon assimilation has not been assessed in awns, or more generally in any floral structures of Andropogoneae.

The PS, the awn, or both could produce photosynthate that contributes to grain filling. The spikelet is green, which suggests that it could be photosynthetically active, but its tiny size (3-6 mm) makes it unclear whether it could assimilate and export enough carbon to contribute to the carbon economy of other floral organs. In contrast, the awn turns brown soon after heading, suggesting limited capacity for photosynthesis. If so, its function must differ from that of the wheat or rice awn. The awn is also small, and in most sorghum lines, missing entirely.

Here we test the hypothesis that the PS and awn contribute photosynthate to the developing seed in sorghum and other members of the tribe Andropogoneae. We report a combination of radioactive and stable carbon isotopic labeling and metabolite analyses, RNA-seq experiments, morphological observations with scanning electron microscopy (SEM), and spikelet removal experiments. Our goal was to assess photosynthetic capacity in these organs. Given that Andropogoneae, including sorghum, are C_4_ NADP-ME subtype^9,10^, we also considered whether the PS had a gene expression pattern consistent with a carbon concentrating mechanism, as suggested for inflorescence tissues in a few other grasses^11–14^.

## RESULTS

### The SS and awn are carbon sinks, the PS is a source

For a qualitative assessment of carbon assimilation, we used a pulse-chase experiment to assess whether sorghum spikelets or the awn could take up ^14^C. Inflorescence branches were exposed to ^14^C for one hour. Measurements were standardized by weight (disintegrations per minute per mg), and presented as a percentage of the total counts in spikelets and awns (Fig. 2A; Table S1). The PS accounted for a significantly greater percentage of ^14^CO_2_ taken up than the SS or awn (p<<0.0001), with the latter being scarcely detectable. This was true whether the inflorescences were intact (attached), or whether the spikelets and awns were detached and lying on the bottom of the flask, suggesting that relative carbon uptake by the individual structures was consistent regardless of the effects of transpiration (difference non-significant, p>0.997).

**Figure 2.**
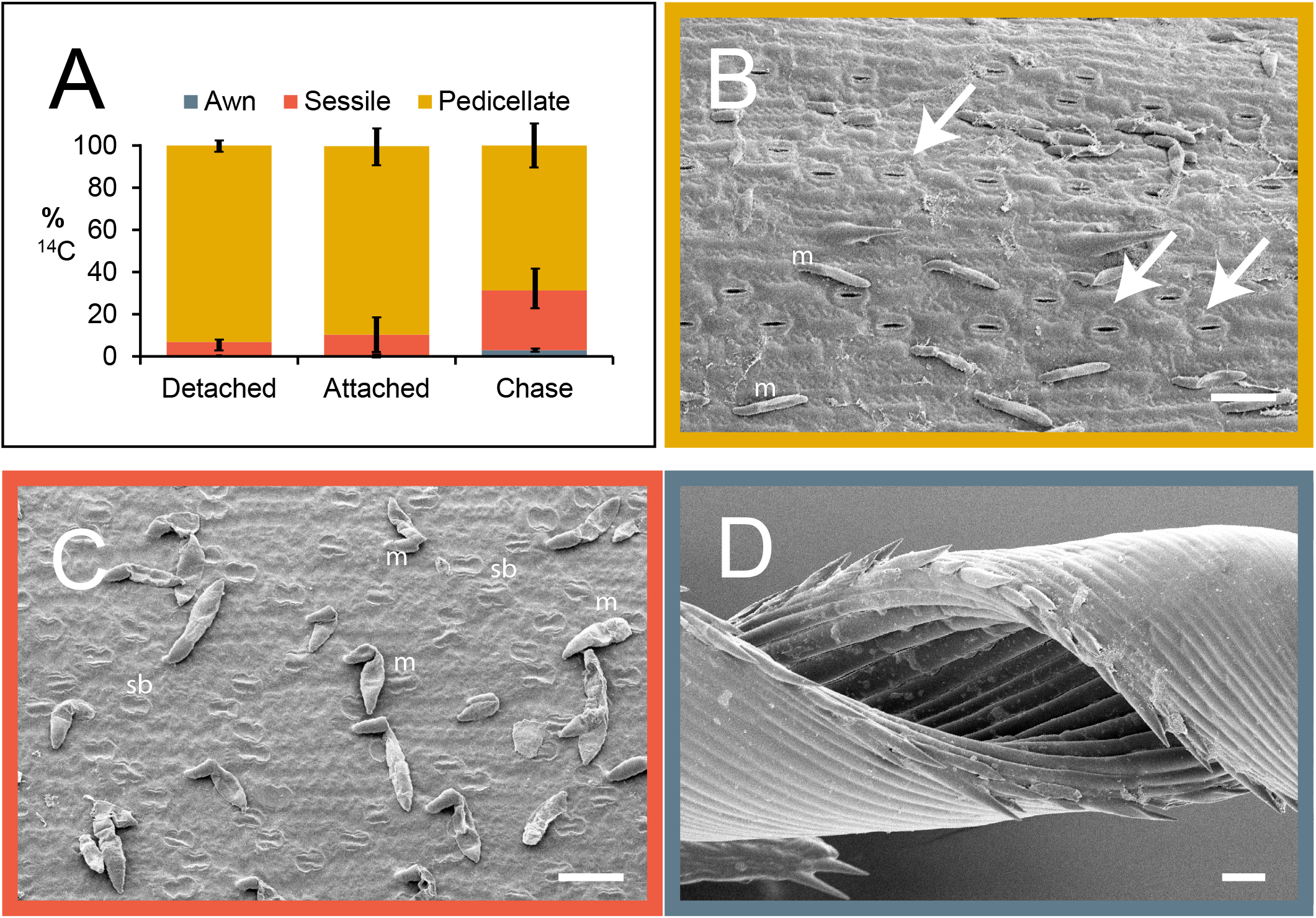
Sorghum bicolor. A. 14C results. Percent dpm for each organ after 1 hour exposure to ^14^C with organs removed from the axis (detached), inflorescence intact (attached), or after 24-hour chase. Plot includes mean percentages and standard deviations, (n=3). Values are similar after 1 hour, whether organs are attached or lying on filter paper. Change after 24-hour chase suggests movement of ^14^C from pedicellate to sessile spikelet. Awn is largely unlabeled. B. Abaxial epidermis of pedicellate spikelet showing rows of stomata (arrows) and bicellular microhairs (m). C. Abaxial epidermis of sessile spikelet showing no stomata, but bicellular microhairs (m) and silica bodies (sb). D. Awn showing no stomata or other epidermal structures, except for prickles on the margin. Scale = 50 μm; note that B and C are more highly magnified than D.

Intact inflorescences were also subjected to a pulse-chase with one-hour pulse labeling followed by 24-hour chase period in air (i.e., no additional labeled carbon provided). Over 24 hours, the fraction of counts in the PS decreased and that of the SS increased significantly (0.01<p<0.5), indicating translocation of carbon from one structure to the other (Fig. 2A, Table S1). The fraction in the awn also increased but was still small relative to the spikelets, and not significantly different from the 1-hour labeling experiment.

To determine whether our result was specific to sorghum or might apply generally to Andropogoneae, we repeated the experiment with the distantly related species *Themeda triandra* and *Andropogon schirensis*, which, like sorghum, have an SS and PS, with an awn on the SS (Fig. S1). These two species represent distinct clades of Andropogoneae and diverged ca. 15 million years ago^15^. The results were similar to those for sorghum: most ^14^C appeared in the PS after 1 hour of labeling, whether attached to the inflorescence or not (difference non-significant, 0.5<p<1.0), but after a 24–hour chase the proportion of label in the PS had decreased and that in the SS had increased significantly (0.0003<p<0.0022 (*A. schirensis*) and 1×10^−5^<p<4×10 ^−5^ (*T. triandra*); Figs. S2, S3; Tables S2, S3). Awns were scarcely labeled.

The surface morphology of each organ is consistent with what would be expected from the ^14^C results. The outer bracts (glumes) of the PS have obvious stomata in all three species (Figures 2B, S2B, S3B), and are similar in this respect to leaves. In contrast, stomata are entirely absent for much of the surface of the glumes of the SS (Figs. 2C, S2C, S3C), although a few can be found near the apex in sorghum and on the sides of the glumes in *A. schirensis* (not shown). No stomata were found on awns (Figs. 2D, S2D, S3D), unlike wheat^7^. SEM data combined with the ^14^C data suggest that the PS may contribute photosynthate to the SS, whereas the awn may not.

### The PS produces Calvin-Benson cycle intermediates, SS and awns are heterotrophic

If the PS is indeed a source of carbon for the grain, it should produce metabolites characteristic of photosynthesis. To test this hypothesis, intact inflorescences were exposed to ^13^CO_2_ for 30s, 60s or 300s, and then key metabolites assessed using LC-MS/MS. Knowing that inflorescence tissues often lack a fully developed C_4_ cycle even in C_4_ plants such as sorghum or maize^13^, we looked for evidence of photosynthetic metabolites, whether from C_4_ (e.g. malate), C_3_ (Calvin-Benson cycle), sucrose (UDPG), or starch (ADPG) biosynthetic pathways. Conversely, we expected that SS and awn should be metabolically distinct, likely heterotrophic.

As in the ^14^C experiments, more carbon was labeled in the PS than in either the SS or awn. In a principal component analysis (PCA) of the metabolite results, the first principal component (PC1) accounted for over 87% of the variance, distinguishing between values for unlabeled metabolites (loading just below 0) and those for labeled metabolites (loading positively)(Fig. 3). By five minutes, metabolite labeling was greatest in the PS, whether considering the weighted average of all isotopologues for each metabolite (i.e., one data point per metabolite per organ and time point; Fig. 3) or individual isotopologues (i.e., three or four data points per metabolite per organ and time point; Fig. S4A). Overall effects of organ, time, and organ with time were all significant except for aspartate (generally p ≤ 0.0001)(Table S4A). Values for the PS were significantly different from those of either of the other two organs (generally p≤0.0001)(Tables S4B, S4C), but the SS and awn were not significantly different from each other. In addition, the five-minute time point was significantly different from the 30 or 60 second pulses (p<0.0001), except for aspartate, but no measurable difference was observed between the early two time points (Tables S4B, S4C).

**Figure 3.**
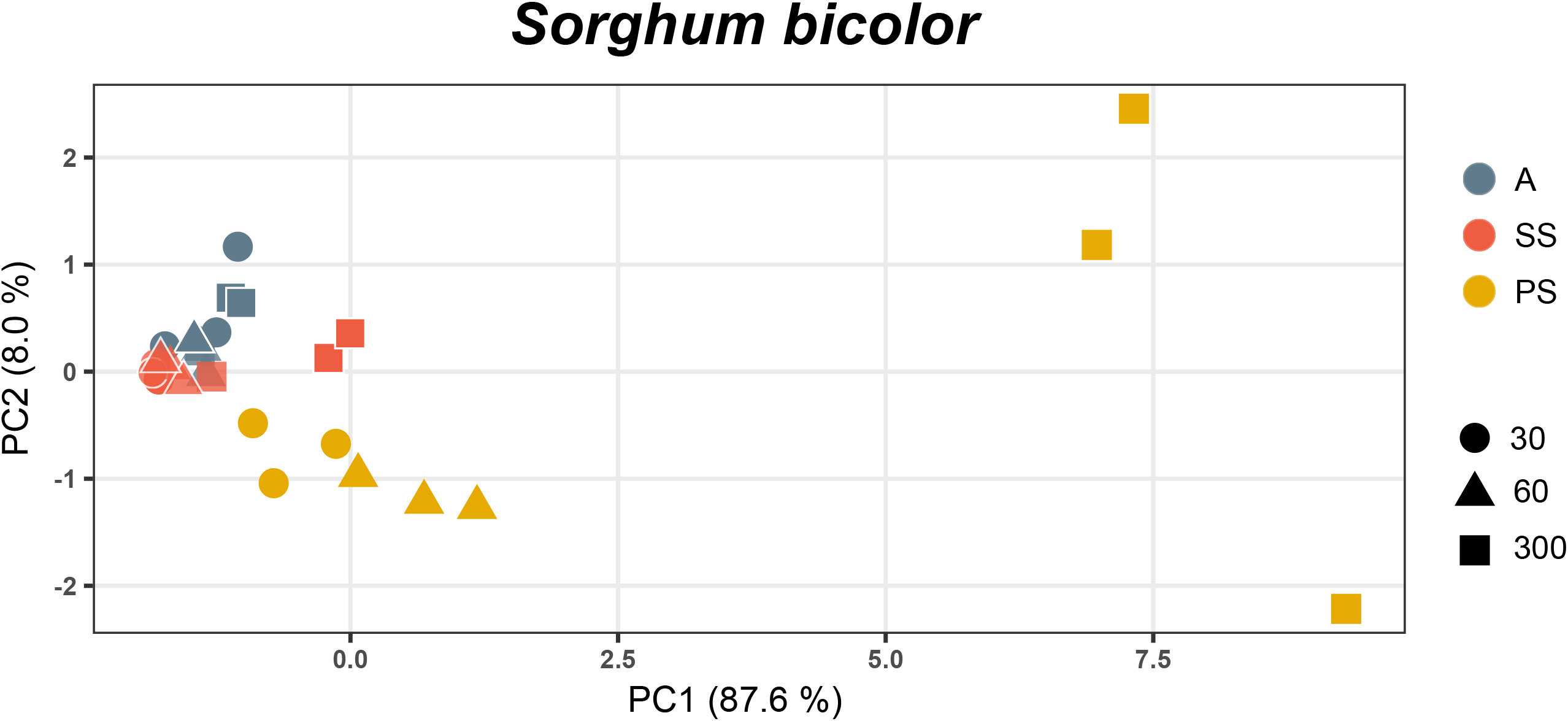
Principal components analysis of ^13^C labeled metabolites in sorghum, average labeling. Values for awn and sessile spikelet are not significantly different at any time point, whereas, values for the pedicellate spikelet are significantly different from the other organs, with the greatest variation in labeling at 5 minutes (300 sec). Organs are distinguished by color, time points by shape. A, awn; SS, sessile spikelet; PS, pedicellate spikelet; 30, values at 30 sec of labeling; 60, values at 60 sec of labeling; 300, values at 300 sec of labeling.

The fraction of unlabeled metabolites (M0) decreased significantly over time in the PS (p generally ≤0.0001), but not in the SS or awn (Fig. 4, Table S5), confirming the ability of the PS to fix carbon. Three-carbon sugar phosphates, the immediate products of RuBisCO-based carbon assimilation, were rapidly labeled (Fig. 4), and the unlabeled fraction constituted < 40% of the total pool in the PS after five minutes. Pyruvate, a product of glycolysis and several steps removed from the Calvin-Benson cycle, was significantly labeled (p ≤ 0.0001), as were hexose phosphates (glucose-6-phosphate, UDP-glucose and ADP-glucose; all p ≤ 0.0001) used in sucrose and starch biosynthesis respectively.

**Figure 4.**
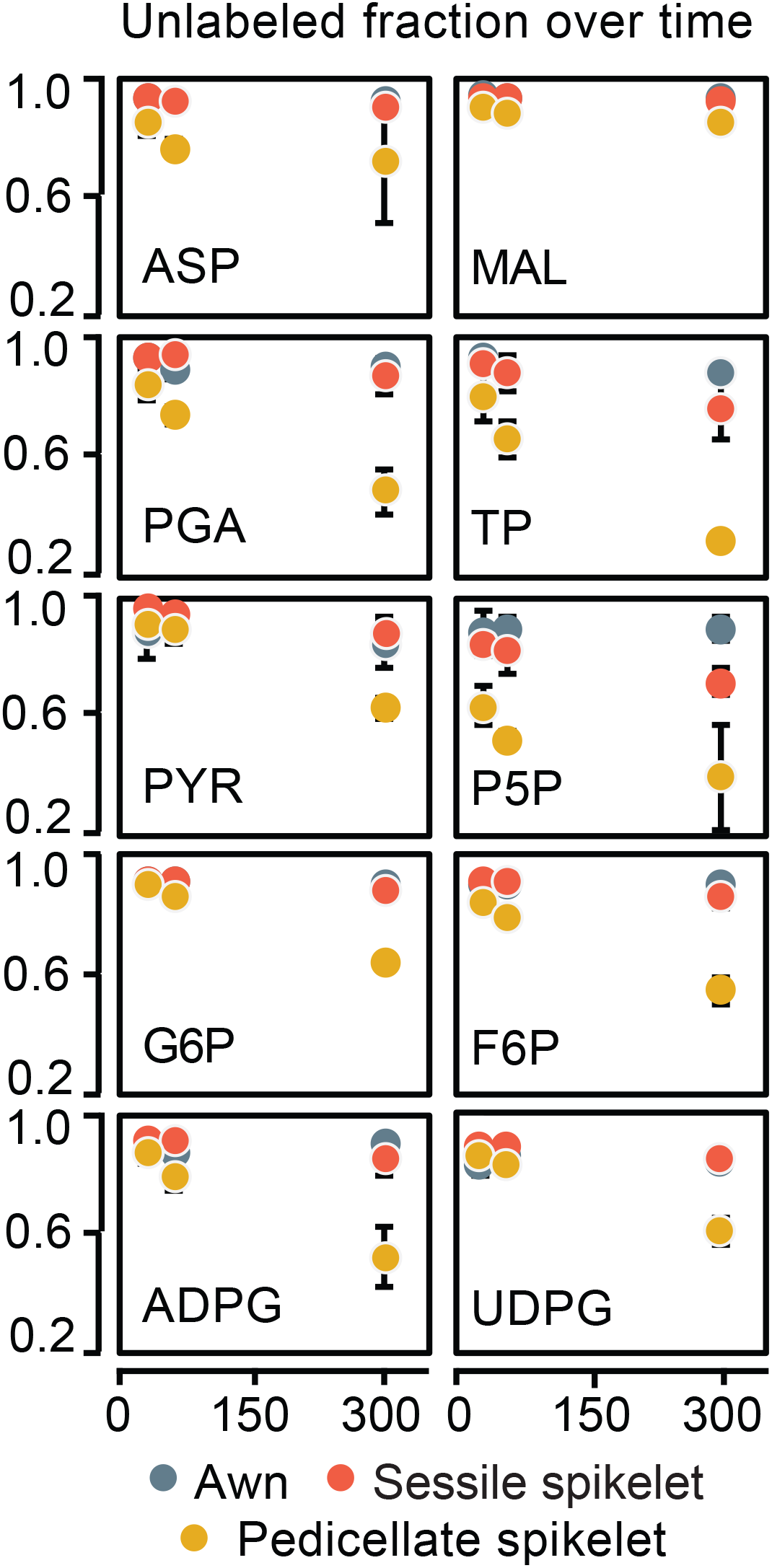
^13^C labeling for individual metabolites at three time points. Fraction of metabolite unlabeled. ASP, aspartate; ADPG, ADP-glucose; F6P, fructose-6-phosphate; G6P, glucose-6-phosphate; MAL, malate; PGA, phosphoglycerate; P5P, pentose 5 phosphates; PYR, pyruvate; TP, triose phosphates; UDPG, UDP-glucose. Points are mean percentages, bars are standard deviations (n=3). Colors distinguish the three organs. Most label accumulation occurs in the pedicellate spikelet and can be seen at 300 seconds.

While determining the extent and type of C_4_ metabolism was not a goal of these experiments, we found that labeling of malate and aspartate was generally low and aspartate was not or only weakly significant (Tables S4, S5). Limited aspartate labeling is consistent with early reports of low PEP carboxykinase (PEPCK) activity in sorghum leaves^16^. We also looked for 2-phosphoglycolate (2-PG), glyoxylate, and glycolate, intermediates associated with photorespiration that could indicate C_3_ metabolism, although low levels of photorespiration may occur in C_4_ plants^17^. Typically only a subset of these particular metabolites are reliably detected or reported^18,19^. 2-PG did become labeled in the PS by 300s, but not in the other organs (Fig. S5).

We investigated vein spacing in glumes of the PS (Fig. S6). Veins are separated by more mesophyll cells than in C_4_ leaves but fewer than those in the SS or C_3_ leaves, hinting that photosynthesis in inflorescence structures is neither wholly C3 nor C4; however more elaborate analyses of flux based on methods specific to these tiny tissues would be necessary to conclusively assess photosynthetic mechanisms, as done by e.g. Hibberd & Furbank^20^ or Henry et al.^14^ in more tractable systems.

To test the generality of our results, we repeated the ^13^CO_2_ experiments with *Themeda triandra*. As in sorghum, the PCA showed a clear distinction between the PS vs. the SS and awn. PC1 explained 66% of the variance and the overall pattern of the PCA was similar to that in sorghum (average labeling, Fig. S7A; Table S6), suggesting that the results may apply to many other members of Andropogoneae. Labeling of individual metabolites was also similar between the two species, except that aspartate labeling was significant (p<0.001) and ADPG, pyruvate, and pentose-5-phosphate labeling were non-significant (Table S6).

### Photosynthetic genes are expressed in leaf and PS, but not SS or awn

To determine if the metabolite data corresponded to underlying patterns of gene expression in the three organ types, we compared transcriptomes for spikelets and awns to those for leaves, focusing on a hand-curated set of 1441 genes encoding enzymes of central carbon metabolism (hereafter called metabolic genes). A PCA of the transcript data separated the leaf and SS along PC1 (Fig. S8), whereas values for awn and PS separated most clearly along PC2, a pattern virtually identical to that of a PCA for all transcripts (not shown). The PCA indicates distinct differences in carbon metabolism among the four organ types. Of the 1441 metabolic genes, 922 were differentially expressed (DE) in at least one pairwise comparison of organs. A heat map of these 922 transcripts (Fig. S9) found that genes upregulated in leaves were down-regulated in the SS and vice versa. Expression in awns and PSs was clearly different from the other two organs. A distinct set of genes was upregulated in awns but these appeared unrelated to either photosynthesis or sugar metabolism.

We examined expression of genes encoding the enzymes that directly produced the ten metabolites measured with ^13^C, a 52-gene subset of the 922 DE genes. Genes were assigned a provisional subcellular location by Target P^21^, although this should be regarded as a preliminary hypothesis of localization. A heat map of this subset (Fig. 5) produced a pattern similar to that for the full set of metabolic genes. Genes specific to C_4_ metabolism, including the copy of phosphoenolpyruvate carboxylase (PEPC) located on chromosome 10 (# 27 in the figure), NADP-dependent malate dehydrogenase (NADP-MDH), and NADP-dependent malic enzyme (NADP-ME) were highly expressed in the leaf, followed by the PS. The gene for the non-C_4_ PEPCK (chromosome 4) was expressed highly in the leaf and PS (Fig. 5), whereas the C_4_-based phosphoenolpyruvate carboxykinase (PEPCK) located on chromosome 1 was not detected, indicating that the sorghum PS, like sorghum leaves, lacks an active PEPCK pathway^22,23^. The small subunit of RuBisCO (transcript #5) was strongly expressed in the leaf, moderately in the spikelets, and scarcely at all in the awn.

**Figure 5.**
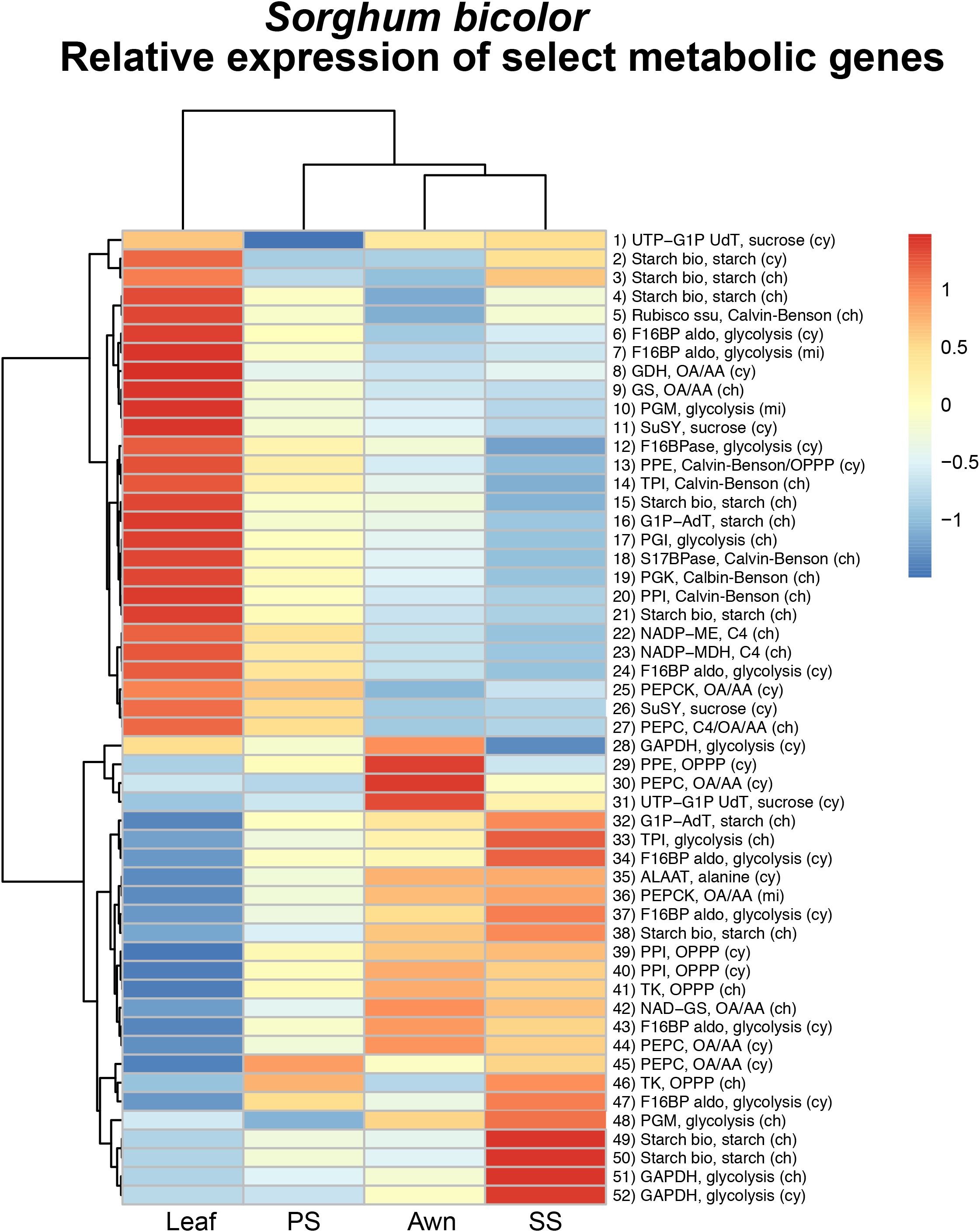
*Sorghum bicolor*. Relative expression of genes encoding biosynthetic enzymes immediately responsible for producing the metabolites labeled with ^13^C, a subset extracted from full set of 922 DE metabolic genes in Fig. S9. Colors reflect scaled z-scores of log2-normalized expression values. Labels of genes indicate enzyme name, biochemical process, and subcellular localization. SS, sessile spikelet; PS, pedicellate spikelet; cy, cytosolic localized; ch, chloroplast localized; mi, mitochondrial localized; OA/AA, organic acid/amino acid metabolism. 1) UTP-G1P UdT, UTP-glucose-1-phosphate-uridyltransferase, Sobic.002G291200.1; 2) Starch synthase, Sobic.001G239500.2; 3) Starch synthase, Sobic.010G047700.1; 4) Starch synthase, Sobic.002G116000.1; 5) Ribulose-1,5-bisphosphate carboxylase/oxygenase, small subunit, Sobic.005G042000.1; 6) F16BP aldo, Fructose-1,6-bis-phosphate aldolase, Sobic.005G056400.1; 7) F16BP aldo, Fructose-1,6-bis-phosphate aldolase, Sobic.008G053200.1; 8) GDH, glutamate dehydrogenase, Sobic.003G188400.1; 9) GS, Glutamine synthetase, Sobic.006G249400.1; 10) PGM, Phosphoglucomutase, Sobic.003G222500.1; 11) SuSY, sucrose phosphate synthase, Sobic.004G068400.1; 12) F16BPase, fructose-1,6-bisphosphatase, Sobic.003G367500.1; 13) PPE, Phosphopentose epimerase, Sobic.001G491000.1; 14) TPI, triose phosphate isomerase, Sobic.002G277100.1; 15) Starch synthase, Sobic.006G221000.1; 16) G1P-AdT, glucose-1-phosphate adenyltransferase, Sobic.007G101500.1; 17) PGI, Phosphoglucoisomerase, Sobic.002G230600.1; 18) S17BPase, sedoheptulose 1,7-bisphosphatase, Sobic.003G359100.1; 19) PGK, Phosphoglycerate kinase, Sobic.009G183700.1; 20) PPI, phosphopentose isomerase, Sobic.001G069000.1; 21) Starch synthase, Sobic.004G238600.1; 22) NADP-ME, NADP-malic enzyme, Sobic.003G036200.1; 23) NADP-MDH, NADP-malate dehydrogenase, Sobic.007G166300.1; 24) F16BPase, fructose-1,6-bisphosphatase, Sobic.010G188300.1; 25) PEPCK, phosphoenol pyruvate carboxykinase, Sobic.004G338000.1; 26) SuSY, sucrose phosphate synthase, Sobic.003G403300.1; 27) PEPC, phosophoenol pyruvate carboxylase, Sobic.010G160700.1; 28) GAPDH, glyceraldehyde-3-phosphate dehydrogenase, Sobic.005G159000.1; 29) PPE, Phosphopentose epimerase, Sobic.002G257300.1; 30) PEPC, phosphoenol pyruvate carboxylase, Sobic.004G106900.1; 31) UTP-G1P UdT, Sobic.006G213100.1; 32) G1P-AdT, glucose-1-phosphate adenyltransferase, Sobic.002G160400.1; 33) TPI, triose phosphate isomerase, Sobic.003G072300.2; 34) F16BP aldo, Fructose-1,6-bisphosphate aldolase, Sobic.004G146000.1; 35) ALAAT, alanine amino transferase, Sobic.001G260701.1; 36) PEPCK, phosphoenol pyruvate carboxykinase, Sobic.006G198400.2; 37) F16BP aldo, Fructose-1,6-bisphosphate aldolase, Sobic.003G393900.1; 38) Starch synthase, Sobic.007G068200.1; 39) PPI, Phosphopentose isomerase, Sobic.003G182400.1; 40) PPI, phosphopentose isomerase, Sobic.008G135701.1; 41) TK, transketolase, Sobic.010G024000.2; 42) NAD-GS, Sobic.003G258800.1; 43) F16BP aldo, Fructose-1,6-bisphosphate aldolase, Sobic.003G096000.2; 44) PEPC, phosphoenol pyruvate carboxylase, Sobic.003G301800.1; 45) PEPC, phosphoenol pyruvate carboxylase, Sobic.002G167000.1; 46) TK, Transketolase_Sobic.009G062800.1; 47) F16BP aldo, Fructose-1,6-bisphosphate aldolase, Sobic.009G242700.1; 48) PGM, phosphoglucomutase, Sobic.001G116500.1; 49) Starch synthase, Sobic.010G022600.1; 50) Starch synthase, Sobic.010G093400.1; 51) GAPDH, glyceraldehyde-3-phosphate dehydrogenase, Sobic.004G056400.1; 52) GAPDH, glyceraldehyde-3-phosphate dehydrogenase, Sobic.004G205100.1.

Analogous data for *Themeda triandra* produced similar but not identical findings. Because *Themeda triandra* spikelets are subtended by a large leaf-like bract (see below), expression of the 769 metabolic genes in the spikelets was compared to that of the bract rather than to a foliage leaf. PCA of these transcripts placed the awns and PS at opposite sides of PC1 (32.9% of variance); the three replicate bract samples were distinguished from the other organs on PC2 (Fig. S10).

Of the 769 genes, 322 were DE between at least one pair of organs. Relative expression of these genes showed a different pattern for each of the four organs (Fig. S11). Because the variability among samples of *T. triandra* was higher than that in sorghum, fewer genes were significantly DE, but overall trends were similar. Awns in *T. triandra* had a distinct set of upregulated genes that were not reflected in photosynthetic metabolism. Of the genes encoding enzymes that could produce the ^13^C metabolites, only 24 were DE (Fig. S12). (Note that *T. triandra* lacks a reference genome, so the genes are labeled with the names of their sorghum orthologues.) Genes related to photosynthesis are upregulated in the leaf-like bract and down-regulated in the SS, whereas the situation is opposite for genes involved in starch and sucrose metabolism. The PS is similar to both bracts and SSs, whereas awns have a unique expression profile. Unlike sorghum, genes for aspartate metabolism are DE, and the C_4_ isoform of PEPCK (#19 in Fig. S12) is moderately expressed in the SS, PS, and bract, suggesting that this species may have an active PCK-type of C_4_ photosynthesis in addition to NADP-ME, and use aspartate to transport carbon between the bundle sheath and mesophyll.

### PS may contribute to seed weight

To test the influence of PS function on plant grain yield, individual PSs were removed from alternating panicle branches at anthesis. Eight different accessions differing in inflorescence architecture and relative size of the PS and SS (Fig. S13) were tested. A preliminary experiment on a small number of plants showed that removal of the PS results in significantly reduced seed weight (p=0.0018; Table S10A), but experiments that additionally included awn removal or controlled for the effects of wounding were inconclusive (Tables S10B and S10C), with an overall effect of PS removal that was not significant (p=0.108 and 0.092, respectively). Analyzing the latter two experiments together found that removal of the PS caused a significant 4.5% reduction in seed weight (p=0.0417)(Fig. S14). The direction of the seed weight response to spikelet removal was generally consistent within individual genotypes, but effect size, statistical significance, and reproducibility were variable, suggesting a need for increased replication to resolve genotype-specific effects of PS removal on seed weight.

### Bracts enhance carbon assimilation in wild species

The contribution and position of photosynthetic organs to seed production was also considered by analyzing the bracts in *Themeda triandra* and *Andropogon schirensis*, which are much closer to the spikelets than the flag leaves of sorghum. Our ^14^C pulse-chase experiments tested whether these played a role in carbon assimilation.

In *A. schirensis*, the modified leaf is reduced largely to a sheath, with only a short blade. As expected, this modified leaf assimilated ^14^C, although the difference in percent counts between the pulse and chase for the attached bract was non-significant (p=0.9987; Figure 6A, Table S8). The leaf sheath bears stomata throughout (Fig. 6B), as expected. The internode between the leaf node and the lowermost inflorescence node contained little ^14^C, with the counts associated predominantly with segments next to the nodes. The leaf is thus not a major source of photosynthate for the inflorescence within the timeframe and conditions of this experiment (anthesis), and the increased ^14^C in SSs during the chase was largely accounted for by the drop in PS values.

**Figure 6.**
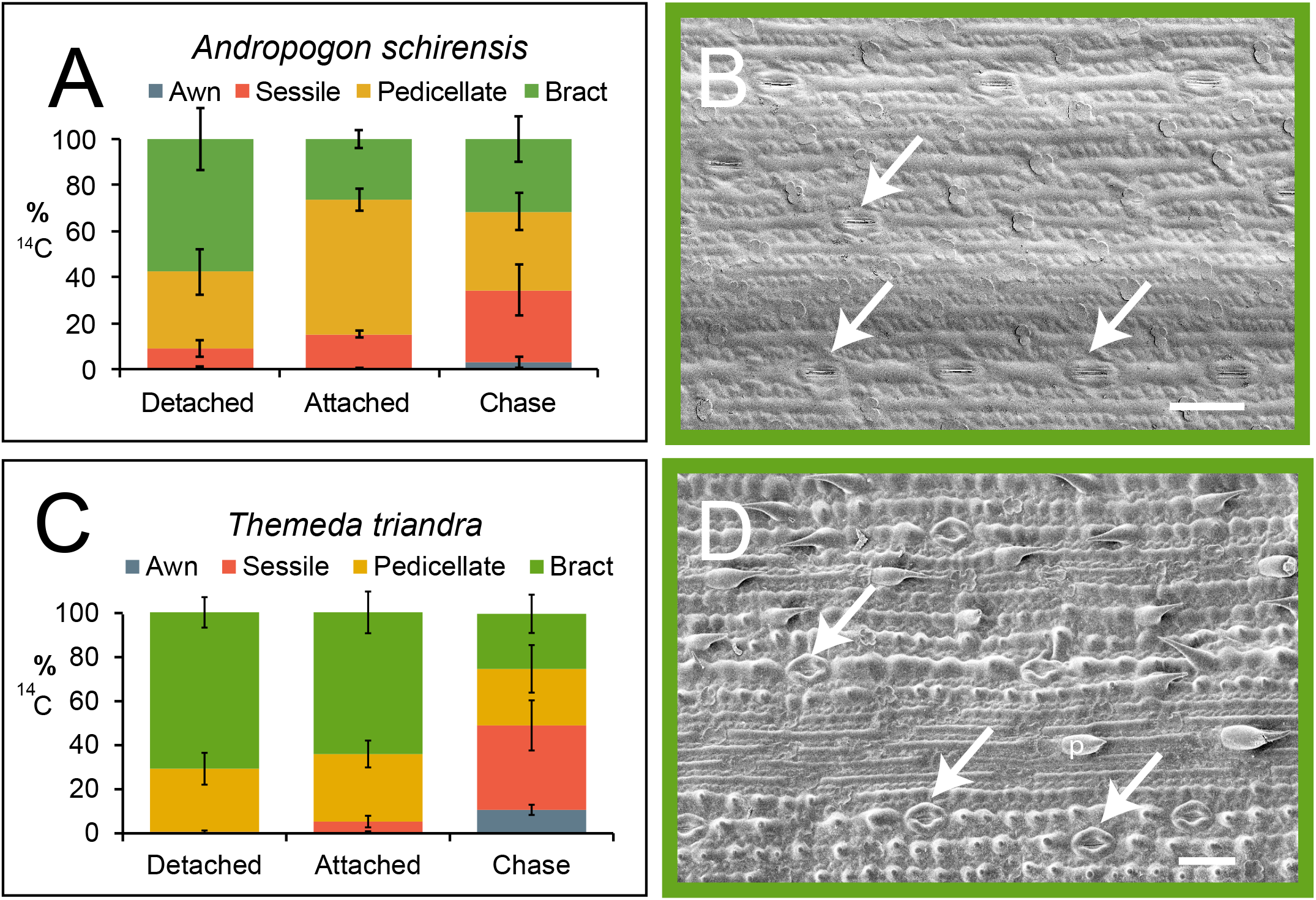
A, C. ^14^C results including bract. Percent dpm for each organ after 1 hour exposure to ^14^C with organs removed from the axis (detached), inflorescence intact (attached), or after 24-hour chase. Plot includes mean percentages and standard deviations, (n=4, *Andropogon;* n=3, *Themeda)*. After 1 hour, most counts are in the bract when organs are detached from the stem but in the pedicellate spikelet when they are attached. Change after 24-hour chase suggests movement of ^14^C from pedicellate to sessile spikelet. Awns remain largely unlabeled. A. *Andropogon schirensis*. C. *Themeda triandra*. B, D. Abaxial epidermis of bract showing rows of stomata (arrows). B. *Andropogon schirensis*. D. *Themeda triandra*, also with prickles (p). Scale = 50 μm.

In contrast, the bract of *Themeda triandra* is closely connected to the spikelets, separated by an internode of only 2-4.5 mm. The bract accounted for 64-71% of the fixed ^14^C in the 1-hour pulse (Table S9), and that percentage dropped to 25% in the chase (p<0.0001), with a corresponding increase in the amount in the SS (p<0.01, Table S9, Fig. 6C). The epidermal pattern of the bract is leaf-like, with extensive stomata (Figure 6D) Results of ^13^C assimilation were qualitatively similar to those for ^14^C (Fig. S4B, all isotopologues; S7B, average of isotopologues for each metabolite), with the bract being significantly different from the other three organs (Table S10). Thus, adding a leaf-like structure close to the spikelets can enhance carbon assimilation in the form of PS or bract and relative contribution is dependent on architecture and location.

## DISCUSSION

Our data suggest that carbon capture in sterile spikelets may help compensate for the approximately 50% reduction in meristems caused by having non-seed-bearing spikelets paired with seed-bearing ones, as summarized in Fig. 7. It is unexpected that one set of floral structures would contribute carbon to the metabolism of other flowers; previously, flowers and inflorescences were reported only to contribute fixed carbon to their own metabolism (e.g.,^24–28^). Reproductive tissues of oilseeds are also often green and may assimilate or re-assimilate carbon^29–34^ and use photosynthetic light energy for ATP or reductant production^31,35–37^. A number of these studies focused on the contribution of sunlight to metabolically demanding processes such as fatty acid biosynthesis that generate large amounts of CO_2_.

**Figure 7.**
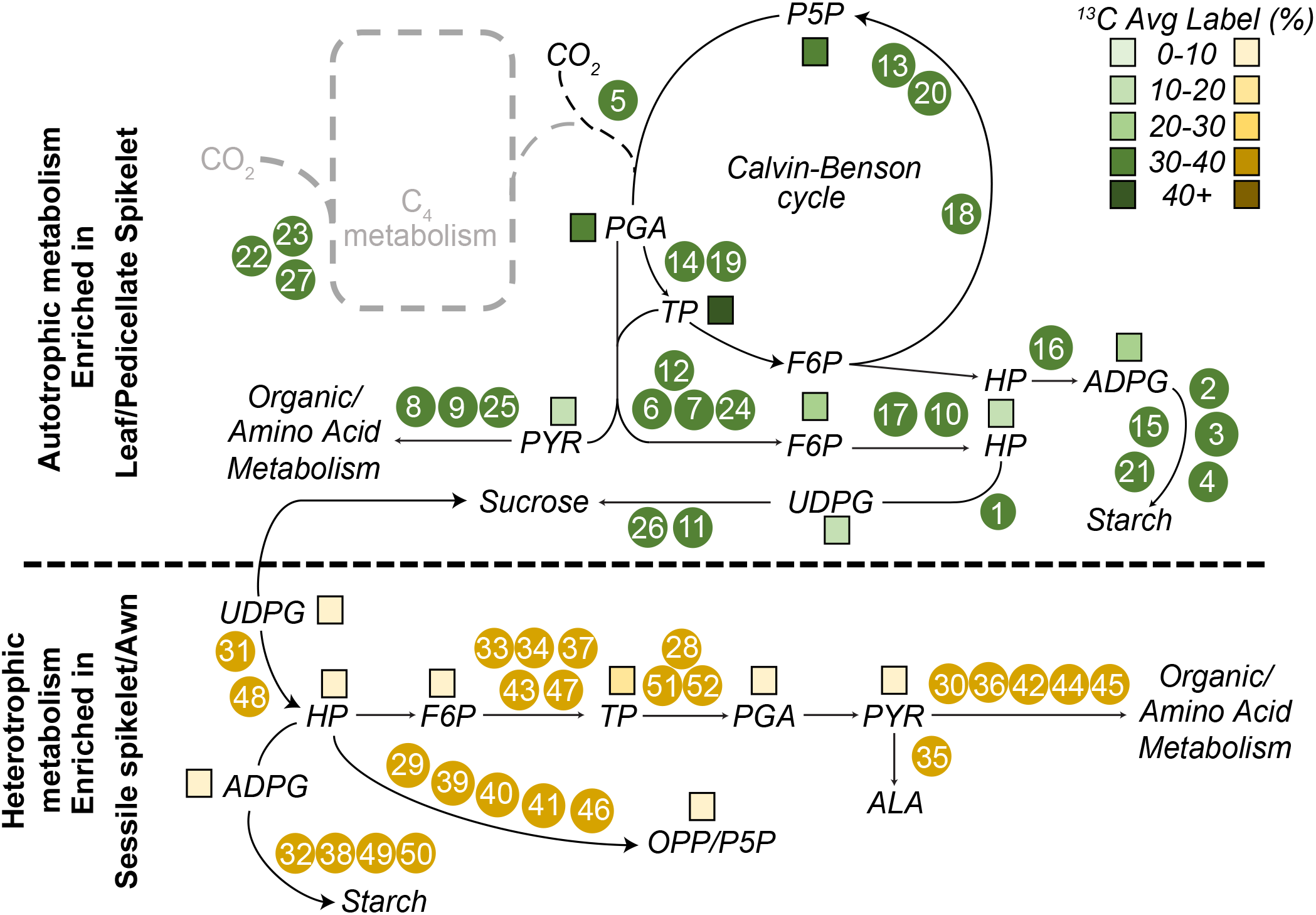
Summary of biochemical pathways assessed by ^13^C and RNA-seq data. Numbered circles correspond to enzymes whose expression is shown in Figure 5. Colored squares reflect percent average label in ^13^C assays of metabolites. Upper part of figure shows autotrophic metabolism and is enriched in the leaf and PS; lower part of figure shows heterotrophic metabolism, enriched in the SS.

The PS exhibits autotrophic metabolism with metabolite and transcript accumulation similar to, but less robust than, a leaf. It can assimilate CO2 whether attached or detached from other structures, consistent with leaf-like epidermal morphology of the glumes, with stomata distributed over most of the surface. The PS might contribute as much as 5% to yield (Fig. S14), an estimate consistent with earlier reports that photosynthesis from the entire sorghum inflorescence may account for 6-18% of yield^38,39^. We can now attribute that contribution to yield to a particular component of the inflorescence.

In contrast, the SS in sorghum appears to be largely heterotrophic even as early as anthesis. Carbon assimilation is lower than in the PS, but still easily detected and stomata are much less common on the surface of the glumes. Expression of transcripts encoding proteins involved in photosynthesis was low in the SS, whereas genes involved in starch synthesis were relatively high (Figs. 5, S9). The decarboxylating enzymes reported as upregulated in tobacco stems^40^ were not upregulated in the SS, providing no evidence for decarboxylation of transported four carbon compounds.

We hypothesized initially that awns in sorghum should be similar to those in wheat, barley, and rice (e.g., ^7,41–44^ and references therein), which are large, green, and provide photosynthate to the developing grain. However, no evidence of CO_2_ assimilation was found in sorghum awns whether isolated or intact (Fig. 2A), none of the metabolites were labeled (Figs. 3, 4), and no stomata are present (Fig. 2D). While the awns exhibit distinct gene expression patterns (Fig. 5), they are generally more similar to the SS. Awns in Andropogoneae are derived independently from those in wheat, barley or rice^45^, and, as shown here, differ in structure and physiology.

The results for sorghum apply to other members of the tribe Andropogoneae, as shown by data on the unrelated species *Andropogon schirensis* and *Themeda triandra*. In both species, the SS is bisexual as in sorghum, has a firm outer glume, and bears an awn. The PS in these species is staminate (male). PSs in these species are photosynthetic and export carbon to the heterotrophic SS at anthesis. The awns do not assimilate carbon and lack any evidence of carbon assimilation, although, as in sorghum, they produce awn-specific gene transcripts.

Because the structure of the spikelet pair is conserved among most of ca. 1200 species of Andropogoneae (maize being an exception), we can infer that the function of the PS as a nurse tissue has been fixed and maintained by natural selection over ca. 15 myr of evolution. The proximity of PSs to seeds may enable delivery of photoassimilates during stress such as drought-induced reduction in transpiration, as described for awns in wheat^44^.

The small photosynthetic PS boosts fitness or yield, but placing a leaf-like structure in the inflorescence closer than the flag leaf appears to be even more favorable. The bract in *Themeda triandra* assimilates considerably more carbon than the PS and transfers that carbon to the SS. Metabolically and anatomically it is leaf-like. The bract in *Andropogon schirensis* is also photosynthetically active, but appears to transfer little carbon to the developing inflorescence. This may reflect the distance of translocation within the short time frames of these experiments. While the distance from the flag leaf to the panicle in sorghum is generally at least 30 cm and often twice that, the corresponding distance in *A. schirensis* is 14-21 cm, and that in *Themeda triandra* is 2-4.5 mm.

Although the PS is undoubtedly photosynthetic, whether it maintains an active C4 cycle is unclear. A number of photosynthetic intermediates were significantly labeled in the PS and the relative expression of photosynthetic enzymes is higher in the PS than the SS or awns, but lower than in leaves or bracts, respectively (Figs. 7, S9), though these data do not distinguish the mode of carbon assimilation^20^. Vein to vein distances in the PS of sorghum are larger than in a leaf (Fig. S6), but closer than in most C_3_ species or in the SS, suggesting the possibility of a limited C_4_ shuttle, as demonstrated in other inflorescence-related structures^13,46^. We also found labeled 2-phosphoglycolate in PSs and some unlabeled 2-phosphoglycolate in other organs indicating possible photorespiration, consistent with either C3 or C4 RuBisCO-based assimilation, though 2-phosphoglycolate phosphatase, an enzyme specific to the photorespiratory pathway, was not DE. Precise determination of photosynthetic pathway operation will require immunolocalization of RuBisCO and PEPC or extended gas analyses^13^ with methods and chambers adapted for small tissue analysis.

In summary, the PS in sorghum and Andropogoneae can make a small but significant contribution to yield by fixing carbon and translocating it to the SS, which holds the developing seed. Awns, in contrast, are carbon sinks, with limited metabolic activity. These results reflect millions of years of evolution, over which the spikelet pair has been selected and conserved. The PS may thus have contributed to fitness in natural populations and could be a useful target for sorghum improvement in agricultural settings.

## METHODS

### Plant material

*Sorghum bicolor* accessions BTx623 (PI 564163), SO85 (PI 534096), Jola Nandyal (PI 534021), SAP-170 (PI 597971), SAP-51 (PI 655995), SAP-15 (PI 656014), SAP-257 (PI 656099), Combine Hegari (PI 659691) were obtained via the USDA Germplasm Resource Information Network (GRIN). All except SO85 are members of the Sorghum Association Panel^47^. BTx623 is the line from which the reference genome sequence was obtained^48,49^. Additional experiments were conducted on *Themeda triandra* (PI 208197), and *Andropogon schirensis (Pasquet s.n.)*.

The three species represent different major clades of Andropogoneae^15^. In addition, they differ in inflorescence structure. In sorghum, the pedicellate spikelets are sterile, whereas in the other two species, the pedicellate spikelets are staminate. In *T. triandra*, the sessile spikelet is associated with two staminate, pedicellate spikelets and is subtended by two additional pairs of staminate spikelets for a total of six staminate spikelets per sessile bisexual spikelet (Fig. S1). For the purposes of this experiment, all six were considered similar and were treated together. In addition, the set of spikelets in *T. triandra* is closely subtended and partially enclosed by a large bract. In *A. schirensis*, there is no bract but the uppermost leaf is reduced to a sheath and minimal blade. The distance between the node of the leaf and the inflorescence node is 14–21 cm. *S. bicolor* has no bracts or inflorescence-associated leaves. In all three species, only the sessile, seed-bearing spikelet bears an awn.

### Pulse-chase experiments,^14^C

#### Labeling

To determine which structures assimilated and fixed CO2, we traced the localization of ^14^C in *Sorghum bicolor* SO85, *Themeda triandra*, and *Andropogon schirensis*. Plants were collected at anthesis in the greenhouse between 10 and 11 AM. Culms were cut with a razorblade and placed directly into tap water before being transferred to 250 ml Erlenmeyer flasks containing filter paper and 1 mL of water, with two flasks per species. One flask (A) contained one (*Themeda, Sorghum*) or two (*Andropogon*) intact inflorescences, and was used for the 24-hour pulse-chase experiments. The second flask (B) for each species held one or two intact inflorescences and was used for the 1-hour pulse experiment. Flask B also contained an additional inflorescence dissected into sessile spikelets, pedicellate spikelets, awns, and bracts or leaves, to determine ^14^C assimilation without connection to the rest of the inflorescence. Each experiment used ca. 40-80 mg of tissue, which for *T. triandra* equated to approximately: 15 awns, 22 bracts, 16-20 sessile spikelets, and 40 staminate spikelets. For *A. schirensis* a similar amount of biomass required approximately 25 awns, 22 sessile spikelets, 30 staminate spikelets, or 1 to 2 leaves. For *S. bicolor* we used 2.5 cm of one leaf, ~6-8 sessile spikelets, ~30 staminate spikelets, and all available awns from one inflorescence.

A plastic tube containing 12.5 microcuries of ^14^C sodium bicarbonate was placed in each flask and maintained in an upright position by attachment to a plastic rod. Flasks were capped with airtight septa closures and 1 mL 6 N H2SO4 was added directly to each plastic tube using a syringe, releasing a pulse of ^14^CO_2_ into the flask. All samples were incubated for one hour in a growth chamber at ~350μE.

#### Processing

After one hour, all flasks were purged with air (30-60 sec) to remove the ^14^CO_2_. The radioactive gas stream was captured in a reservoir containing 2L of 2N KOH. For Flask A for each species, the airtight septum and closure was replaced with a sponge top prior to incubation in the growth chamber under continuous illumination for the 24-hour chase period. The contents flask B were analyzed immediately. Detached awns, sessile and pedicellate spikelets, and leaves were separated, weighed, and transferred to tubes containing cold methanol:chloroform (7:3, v:v) and steel beads. Simultaneously, the intact inflorescence was dissected, and individual components treated identically to the detached samples. Tissues were homogenized at 30 cycles/second in a bead homogenizer for two 5-minute intervals and were stored at −20°C for 48 hours. After 24 hours, samples from flask A were dissected, weighed and processed in the same way and stored at −20C for 24 hours.

All samples were extracted sequentially. The first extraction was based on 1.5 mL 7:3 (v:v) methanol:chloroform that was used to homogenize and store tissues after labeling. 200 uL from the methanol:chloroform extract was then combined with 5 mL of scintillation fluid (HIONIC FLUOR, Perkin Elmer). The remaining extract was removed from the residual biomass and water (2 mL) was added. The biomass was bead homogenized as before, centrifuged, and 200 uL was combined with 5mL scintillation fluid. Residual biomass was then treated with tissue solubilizer (ScintiGest), incubated overnight at 60°C and prepared for scintillation identically to prior extracts. Scintillation counting in disintegrations per minute (DPM) was performed on a Beckman Coulter LS-6000TA Scintillation Counter and included recording ^14^C photon emissions for five minutes. After background subtraction from a blank that contained identical amounts of solvent and scintillation cocktail, total radioactivity per amount of tissue was calculated by accounting for differences in volume and mass.

#### Analysis of ^14^C data

The total ^14^C assimilation (DPM/mg) for each organ was calculated by summing the counts from the three serial extractions and accounting for the total volumes of each extraction. Then the percent of label within awn, pedicellate and sessile spikelets was determined by calculating the fractional ^14^C assimilation in each organ. Experiments were repeated three (*Sorghum, Themeda*) or four (*Andropogon*) times over a ten-month span using individual plants harvested from a greenhouse. Percent label for each organ was averaged across experiments, and means and standard deviations plotted in MS Excel.

### ^13^C labeling in planta

#### Labeling

^13^C isotopic labeling studies were carried out on inflorescences of *Themeda triandra* and *Sorghum bicolor* accession SO85. *Themeda* and *Sorghum* plants were grown in the greenhouse until anthesis. The intact inflorescence was placed in a deflated plastic bag (inflated volume ca. 2-3 liters). A 10 mL serological pipet was fed into the bag with its tip near the apex of the inflorescence. The other end of the pipet was connected by hose to a tank of synthetic air comprised of ^13^CO_2_/N_2_/O_2_ at a ratio of: 0.033:78:21.967. For each timed treatment, the bag was rapidly inflated (~15 L/min), and then the flow of gas was decreased to approximately 2 L/min. Structures were labeled for 30, 60 or 300 seconds. Three replicate experiments were done for each time point for each species. For sorghum, each replicate was a single inflorescence from a single plant, making a total of nine inflorescences (3 reps x 3 time points). For *Themeda*, each replicate included all flowering structures on a single tiller from a single very large plant, making a total of nine tillers (3 reps x 3 time points). All experiments were conducted in the morning between 10 and 11 AM to minimize circadian effects. The synthetic air pumped into the bag flowed from the release of the pipet at the tip along the inflorescence structures and exited the bag at the point where the bag opening was grasped around the culm. At the end of the labeling period the inflorescence was cut immediately and dropped in a large pool of liquid nitrogen in a Styrofoam box (22 x 33 x 15 cm) to quench metabolism. During labeling and quenching plant tissues were exposed to greenhouse light levels between 250-400 uE.

#### Processing

Frozen tissue was dissected in liquid nitrogen to separate sessile and pedicellate spikelets, awns, and bracts. Each sample was then ground with liquid nitrogen in mortar and pestle, extracted with methanol:chloroform (7:3 v:v) and then through addition of water to segregate polar and non-polar phases as described previously^50,51^. PIPES (12 nmol) was added to each sample as an internal standard.

#### LC-MS/MS

A QTRAP 6500 tandem mass spectrometer linked to two Shimadzu LC-20AD pumps working in coordination with a SIL-20AC/HT autosampler was used to assess and quantify the isotopic labeling in intermediates of primary metabolism. Extract from ca. five percent of the total harvested sample was injected. Standards for metabolites were run separately to establish retention time and, in some cases, confirm identification of isomers. Separation on LC involved an ion pair method^51^ with flowrate of 300 microliters/min and a binary gradient buffer combination with Buffer A: 11 mM acetic acid with 10 mM tributylamine as the ion pair and Buffer B: 100% methanol. The method differed from prior work as the ramp profile was shortened with 3 min equilibration at 5% B, 10 min ramp to 35% B, 2 min ramp to 95% B, hold for 3 min, return to 5% B within 2 min, and equilibrate there for 11 min resulting in a total run time of 31 min The source inlet temperature (550°C), curtain gas (35 psi) and auxiliary gases (both set to 60 psi) were chosen based on optimal peak response. Declustering potential, collision energy and collision exit potential for individual mass transition pairs of multiple reaction monitoring were based on prior work ^50,51^.

#### Analysis of mass spectrometry data

Peak areas for individual mass traces that represent a precursor-product ion combination were integrated using MultiQuant 3.0.2 and Analyst 1.6 (AB Sciex). The relative percent combination of isotopologues was calculated, as well as ^13^C average labeling that is defined as 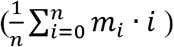 with n equal to the number of carbons and m_i_ defining the relative isotopologue abundance for each of the i isotopologues measured. Data were additionally compared through the decrease in unlabeled fraction of the metabolite pool, referred to as M0 in the text. Average labeling and unlabeled fractions were plotted with standard deviations and in addition were analyzed with PCA to assess repeatability. Statistical significance of results was assessed with ANOVA. Both PCA and ANOVA used standard programs in R, as presented in the text and figures.

### Sorghum spikelet removal experiments

#### Removal of pedicellate spikelets

The impact on grain weight of removing pedicellate spikelets was tested in all eight accessions (genotypes) of *Sorghum bicolor*. Three to five pots were prepared for each genotype, with each pot containing two plants, one of which had a subset of pedicellate spikelets removed (the treated plant, plant A) and the other of which remained intact (the untreated plant, plant B). When both plants reached anthesis, pedicellate spikelets were removed with forceps from alternating branches of the plant A, which thus had a mix of treated and untreated branches. By removing pedicellate spikelets from alternating branches, variation along the inflorescence was averaged out. The untreated branches of plant A thus serve as an internal control, and plant B is an external control. Treated branches were marked. Plants were grown to maturity, the inflorescence removed and dried at 40°C for 4-5 days, and spikelets harvested. Seeds were removed from the glumes and floral bracts. At least fifty seeds were recorded per treatment per plant although usually many more were available.

#### Removal of pedicellate spikelets, awns, and pedicellate spikelets+awns

Five of the genotypes bear awns on the sessile spikelet. These were grown with four plants per pot, six pots per genotype, with treatments as pedicellate spikelets removed (plant A), awns removed (plant B), awns and pedicellate spikelets removed (Plant C), and untreated (plant D). As with the first experiment, structures were removed from about half the branches of the treated plants in an alternating pattern up the inflorescence. Plants were grown, harvested, and mean seed weight calculated as in the previous experiment.

#### Data analysis

Seeds for each treatment/genotype combination were spread on a flatbed Epson Perfection V550 scanner (300dpi resolution, greyscale), imaged and weighed. The resulting JPG files were converted to binary images using Fiji^52^ and the Analyze Particles tool used to count the seeds in each image. Total seed weight was divided by the number of seeds to estimate individual seed weight. The statistical effect of pedicellate spikelet, awn, and pedicellate spikelet and/or awn removal on seed weight was assessed using a mixed effect linear model (*lmer* function in R packages *lme4*^53^ and *lmerTest*^54^, where genotype, pot, and experiment were specified as random effects. All data were visualized using *ggplot* ^55^.

### Scanning electron microscopy (SEM)

Plant material was fixed in formalin:acetic acid:50% alcohol (FAA) for a minimum of 24 hours and then transferred to 50% ethanol. Material was then taken through a standard ethanol dehydration series (70,80,90,95,100,100,100% EtOH) with at least 24 hours per stage. Samples were dried in a SamDri-780 critical point drier at the Washington University Center for Cellular Imaging, coated with gold palladium, and imaged on a Zeiss Merlin Field Emission SEM at 5.0 kV. Photographs were adjusted for brightness, contrast, and input levels using Adobe Photoshop.

### RNA-Seq

To complement ^13^C mass spec measurements of metabolites, expression levels for genes controlling photosynthesis, sugar and starch metabolism were estimated with RNA-seq. As with the ^13^C experiments, material of *Sorghum bicolor* and *Themeda triandra* was harvested at anthesis and immediately frozen in liquid nitrogen and dissected while frozen. Material was harvested between 10 and 11 AM to minimize circadian effects. For sorghum, mature leaves and inflorescences were harvested from three plants and used as individual biological replicates. Inflorescences were hand dissected into pedicellate spikelet, sessile spikelet, and awn samples. For *Themeda triandra*, inflorescences from three tillers from one individual were dissected into bract, pedicellate spikelet, sessile spikelet, and awn, each tiller representing a biological replicate. Each of the 24 samples (2 species x 4 organs x 3 biological replicates) was individually ground to a fine powder using liquid nitrogen. To extract total RNA, ca. 100 mg of tissue was added to 1 mL TRIzol reagent (15596026, Thermo Fisher Scientific) and the samples were vortexed. Next, 0.2 mL of chloroform was added, the samples were shaken for 15 s and incubated at room temperature for 3 min followed by centrifugation at 13,000 g for 15 min at 4°C and the upper aqueous phase was removed and added to 0.5 mL of isopropanol. The sample was incubated on ice for 10 min and centrifuged as above to obtain a nucleic acid pellet. The pellet was washed in 70 % (v/v) ethanol, air-dried briefly, and dissolved in 100 μL DEPC-treated water. Total RNA was cleaned and concentrated using an RNeasy kit (74104, QIAGEN) including on-column DNase treatment (79254, QIAGEN). Total RNA was quantified with a Qubit RNA HS kit (Q32852, Thermo Fisher Scientific). At least one sample for each organ was run on a RNA Pico Bioanalyzer microfluidics chip (Agilent) to calculate RNA integrity numbers (RIN). All RIN were in the range 7.8 – 9.5.

The 24 RNA sequencing (RNA-Seq) libraries were prepared using SENSE mRNA library prep kit V2 for Illumina (001.24, Lexogen) with 500 ng input for all samples except for the three *Themeda* awn samples (160, 420, and 350 ng, respectively). The protocol was adjusted to produce a mean insert size of ~413 bp. Each library was indexed uniquely and amplified by 12 cycles of PCR for all samples except for the three *Themeda* awn samples, which were amplified using 14 cycles. Final libraries were quantified using a Qubit DNA HS kit (Q32854, Thermo Fisher Scientific). Select libraries were also run on a High Sensitivity DNA Bioanalyzer microfluidic chip (Agilent) to confirm library size in the desired range (~400 bp). Libraries were pooled by organism (3 biological replicates x 4 organs, each), resulting in 2 pools of 12 libraries at ~0.8 nM per library (10 nM final pool concentration). These pools were sequenced using Illumina HiSeq4000 paired end (2 x 150 bp) technology at Michigan State University Genomics facility (https://rtsf.natsci.msu.edu/genomics/sequencing-services/).

#### Data analysis

See Table S11 for numbers of transcripts at each step.

a. <u>Read-mapping in sorghum.</u> Reads for *S. bicolor* were trimmed using Trimmomatic^56^ and mapped to the sorghum reference genome (Phytozome, Sbicolor_454_v3.1.1) using HISAT^57^. To create expression level analysis, *htseq-count* was used to count the number of reads per gene for each sample^58^.
b. <u>Read-mapping and identification of orthologs in Themeda.</u> Reads for *Themeda triandra* were downloaded and submitted to Trinity^59^ for automated trimming, quality filtering, assembly, and expression quantification. CDS and peptides were extracted using transdecoder^60^. Orthologs of the metabolic genes in sorghum were identified by using OrthoFinder^61^ on the longest isoform of peptide sequences from the sorghum and maize genomes, and the predicted *Themeda* peptides. If an orthogroup contained a sorghum metabolism gene as identified in the sorghum transcriptome analysis, the *Themeda* peptides in that group were blasted against the sorghum peptides. If the *Themeda* gene’s best match was a sorghum metabolism gene it was considered that gene’s ortholog. The list of *Themeda* metabolism genes generated from this approach was then used for downstream analyses. Read counts per gene were generated with *htseq-count* ^8^.
c. <u>Normalization for library size and log_2_.</u> All normalization was performed in R^62^. Raw transcript counts for each sample were normalized for library size across all samples for *S. bicolor* and *T. triandra* using the *calcNormFactors* function in the *edgeR* package^63^ and filtered to remove transcripts with fewer than 1 normalized read count in at least 3 samples, effectively discarding unexpressed transcripts. Next, library size-normalized read counts were log_2_-transformed. This step led to 17232 unique transcripts in sorghum and 20826 in *Themeda*.
d. <u>Construction of the metabolic gene set.</u> We developed a custom list of genes involved in carbon metabolism to enable comparison between the RNA-seq data and the carbon isotope data. Manually annotated pathway descriptions are based on the Kyoto Encyclopedia of Genes and Genome (KEGG) pathway and biochemical textbook descriptions supported by comparative organ expression profile data. Enzyme Commission (EC) numbers of enzymes involved in carbon metabolism were retrieved from the KEGG pathway database (https://www.genome.jp/kegg/kegg3a.html) for glycolysis/gluconeogenesis, the tricarboxylic acid (TCA) cycle, pentose phosphate pathway, pentose glucuronate interconversions, fructose/mannose interconversions, galactose metabolism, starch sucrose metabolism, sugar nucleotide metabolism, pyruvate metabolism, glyoxylate/dicarboxylate metabolism, oxidative phosphorylation, photosynthesis, Calvin cycle, etc. In instances where an enzyme could not be unambiguously assigned to a pathway, TargetP^21^, a protein localization prediction tool, was used to determine possible organellar targeting and to guide descriptions. Genes not obviously targeted to chloroplasts or mitochondria were assumed to be cytoplasmic. A subset of C_4_ genes was identified based on phylogenetic literature and comparative genomic analysis^64,65^. Expression levels of genes encoding these enzymes for each species and tissue were then extracted from the full transcriptomes using custom python scripts (https://github.com/ekellogg-lab/pedicellate-spikelet-carbonlab/pedicellate-spikelet-carbon), giving us a total of 3505 loci related to carbon metabolism. The intersection of this list with the log_2_ normalized filtered transcripts left us with 1441 transcripts in sorghum and 769 in *Themeda*. We refer to this list as our set of metabolic genes.
e. <u>Identification of DE genes.</u> Differential expression analysis was performed in R^62^. Tukey tests were performed on the expression values of the metabolic gene set using the function *TukeyHSD* in conjunction with ANOVA (Analysis of Variance) to generate all possible pairwise organ comparisons for each transcript in our metabolically-relevant gene list. The p-values generated by this approach were additionally corrected for multiple testing using *p.adjust* with method set to “BH” (Benjamini-Hochberg). 922 DE genes were identified in sorghum with at least one significant pairwise organ comparison (BH-corrected p-value < 0.05), and 322 DE genes were identified in *Themeda*.
f. <u>Data display.</u> All normalized transcripts and the normalized set of metallic genes were analyzed by Principal Component Analysis (PCA) using the function *prcomp* with log_2_-transformed expression values, and visualized with functions in the *ggplot2* package^55^. DE genes were displayed as expression heatmaps were plotted using *pheatmap*^66^ with the ward.D2 clustering option and row scaling.
g. <u>Genes to compare to ^13^C data.</u> To connect the RNA-seq and ^13^C data directly, we generated a list of 36 enzymes that could produce the metabolites we assayed with ^13^C; some of these are encoded by more than one gene. We then used this list of enzymes to generate a small focused subset of the DE metabolic genes, giving us 52 and 27 DE genes in sorghum and *Themeda*, respectively.

#### Data availability

Raw reads from the RNA-seq experiment have been deposited at GEO (accession number XXX; to be filled in upon acceptance). Data matrices for downstream analyses have been deposted at Dryad (datadryad.org; accession number XXX; to be filled in upon acceptance).

## Acknowledgements

This work was supported in part by NSF grant DEB-1457748 to EAK, and through USDA-ARS support to DKA. The Proteomics and Mass Spectrometry Core facility is acknowledged for technical assistance and mass spectral data was obtained on an instrument funded by NSF-MRI grant DBI-1427621.

## Author contributions

Conceptualization, Methodology, Investigation, Writing – Original Draft, Visualization: TA, VC, DKA, EAK. Formal Analysis and Writing – DG, VC, DKA, EAK. Review & Editing: All authors. Supervision: EAK and DKA. Project Administration: EAK. Funding Acquisition: EAK.

---

Figure 1. Spikelet pair structure in *Sorghum bicolor*. A. Illustration of two spikelet pairs, marked by dotted lines, plus a terminal spikelet, which is morphologically identical to a pedicellate spikelet. Each pair is composed of a sessile spikelet, which includes a bisexual flower and bears the seed, and a pedicellate spikelet, which may be either sterile (most commonly) or staminate. The sessile spikelet bears a twisted awn from the lemma (floral bract). B. Spikelet pair of sorghum accession SAP-15 (PI 656014). SS, sessile spikelet; PS, pedicellate spikelet. Scale bar = 1 mm.

Figure 2. *Sorghum bicolor*. A. ^14^C results. Percent dpm/mg for each organ after 1 hour exposure to ^14^C with organs removed from the axis (detached), inflorescence intact (attached), or after 24-hour chase. Plot includes mean percentages and standard deviations (n=3). Values for detached and attached organs after 1 hour are not significantly different (p>0.99). ^14^C uptake by awns is negligible and is not significantly different among treatments (p>0.99). Values for PS and SS significantly decrease (0.001<p<0.05) and increase (0.01<p<0.05), respectively, after the 24-hour chase, suggesting movement of ^14^C from pedicellate to sessile spikelet. B. Abaxial epidermis of pedicellate spikelet showing rows of stomata (arrows) and bicellular microhairs (m). C. Abaxial epidermis of sessile spikelet showing no stomata, but bicellular microhairs (m) and silica bodies (sb). D. Awn showing no stomata or other epidermal structures, except for prickles on the margin. Scale = 50 μm; note that B and C are more highly magnified than D.

Figure 3. Principal components analysis of ^13^C labeled metabolites in sorghum. Each point is the weighted average of all labeled isotopologues for a given metabolite, organ, and time point (average labeling). Values for awn and sessile spikelet are not significantly different at any time point (0.28<p<0.99), whereas values for the pedicellate spikelet are significantly different from the other organs (p<<0.0001 in most cases), with the greatest variation in labeling at 5 minutes (300 sec; Table S4). Organs are distinguished by color, time points by shape. A, awn; SS, sessile spikelet; PS, pedicellate spikelet; 30, values at 30 sec of labeling; 60, values at 60 sec of labeling; 300, values at 300 sec of labeling.

Figure 4. ^13^C labeling for individual metabolites at three time points. Fraction of metabolite unlabeled. ASP, aspartate; ADPG, ADP-glucose; F6P, fructose-6-phosphate; G6P, glucose-6-phosphate; MAL, malate; PGA, phosphoglycerate; P5P, pentose 5 phosphates; PYR, pyruvate; TP, triose phosphates; UDPG, UDP-glucose. Points are mean percentages, bars are standard deviations (n=3). Colors distinguish the three organs. Most label accumulation occurs in the pedicellate spikelet and can be seen at 300 seconds. Significance values in Table S5.

Figure 5. *Sorghum bicolor*. Relative expression of genes encoding biosynthetic enzymes immediately responsible for producing the metabolites labeled with ^13^C, a subset extracted from full set of 922 DE metabolic genes in Fig. S9. Colors reflect scaled z-scores of log_2_-normalized expression values. Labels of genes indicate enzyme name, biochemical process, and subcellular localization. SS, sessile spikelet; PS, pedicellate spikelet; cy, cytosolic localized; ch, chloroplast localized; mi, mitochondrial localized; OA/AA, Organic Acid/Amino Acid metabolism; 1) UTP-G1P UdT, UTP-glucose-1-phosphate-uridyltransferase, Sobic.002G291200.1; 2) Starch synthase, Sobic.001G239500.2; 3) Starch synthase, Sobic.010G047700.1; 4) Starch synthase, Sobic.002G116000.1; 5) Ribulose-1,5-bisphosphate carboxylase/oxygenase, small subunit, Sobic.005G042000.1; 6) F16BP aldo, Fructose-1,6-bis-phosphate aldolase, Sobic.005G056400.1; 7) F16BP aldo, Fructose-1,6-bis-phosphate aldolase, Sobic.008G053200.1; 8) GDH, glutamate dehydrogenase, Sobic.003G188400.1; 9) GS, Glutamine synthetase, Sobic.006G249400.1; 10) PGM, Phosphoglucomutase, Sobic.003G222500.1; 11) SuSY, sucrose phosphate synthase, Sobic.004G068400.1; 12) F16BPase, fructose-1,6-bisphosphatase, Sobic.003G367500.1; 13) PPE, Phosphopentose epimerase, Sobic.001G491000.1; 14) TPI, triose phosphate isomerase, Sobic.002G277100.1; 15) Starch synthase, Sobic.006G221000.1; 16) G1P-AdT, glucose-1-phosphate adenyltransferase, Sobic.007G101500.1; 17) PGI, Phosphoglucoisomerase, Sobic.002G230600.1; 18) S17BPase, sedoheptulose 1,7-bisphosphatase, Sobic.003G359100.1; 19) PGK, Phosphoglycerate kinase, Sobic.009G183700.1; 20) PPI, phosphopentose isomerase, Sobic.001G069000.1; 21) Starch synthase, Sobic.004G238600.1; 22) NADP–ME, NADP-malic enzyme, Sobic.003G036200.1; 23) NADP–MDH, NADP-malate dehydrogenase, Sobic.007G166300.1; 24) F16BPase, fructose-1.6- bisphosphatase, Sobic.010G188300.1; 25) PEPCK, phosphoenol pyruvate carboxykinase, Sobic.004G338000.1; 26) SuSY, sucrose phosphate synthase, Sobic.003G403300.1; 27) PEPC, phosophoenol pyruvate carboxylase, Sobic.010G160700.1; 28) GAPDH, glyceraldehyde-3-phosphate dehydrogenase, Sobic.005G159000.1; 29) PPE, Phosphopentose epimerase, Sobic.002G257300.1; 30) PEPC, phosphoenol pyruvate carboxylase, Sobic.004G106900.1; 31) UTP-G1P UdT, Sobic.006G213100.1; 32) G1P-AdT, glucose-1-phosphate adenyltransferase, Sobic.002G160400.1; 33) TPI, triose phosphate isomerase, Sobic.003G072300.2; 34) F16BP aldo, Fructose-1,6-bisphosphate aldolase, Sobic.004G146000.1; 35) ALAAT, alanine amino transferase, Sobic.001G260701.1; 36) PEPCK, phosphoenol pyruvate carboxykinase, Sobic.006G198400.2; 37) F16BP aldo, Fructose-1,6-bisphosphate aldolase, Sobic.003G393900.1; 38) Starch synthase, Sobic.007G068200.1; 39) PPI, Phosphopentose isomerase, Sobic.003G182400.1; 40) PPI, phosphopentose isomerase, Sobic.008G135701.1; 41) TK, transketolase, Sobic.010G024000.2; 42) NAD–GS, Sobic.003G258800.1; 43) F16BP aldo, Fructose-1,6-bisphosphate aldolase, Sobic.003G096000.2; 44) PEPC, phosphoenol pyruvate carboxylase, Sobic.003G301800.1; 45) PEPC, phosphoenol pyruvate carboxylase, Sobic.002G167000.1; 46) TK, Transketolase_Sobic.009G062800.1; 47) F16BP aldo, Fructose-1.6-bisphosphate aldolase, Sobic.009G242700.1; 48) PGM, phosphoglucomutase, Sobic.001G116500.1; 49) Starch synthase, Sobic.010G022600.1; 50) Starch synthase, Sobic.010G093400.1; 51) GAPDH, glyceraldehyde-3-phosphate dehydrogenase, Sobic.004G056400.1; 52) GAPDH, glyceraldehyde-3-phosphate dehydrogenase, Sobic.004G205100.1.

Figure 6. A, C. ^14^C results including bract. Percent dpm for each organ after 1 hour exposure to ^14^C with organs removed from the axis (detached), inflorescence intact (attached), or after 24-hour chase. Plot includes mean percentages and standard deviations, (n=4, *Andropogon*; n=3, *Themeda)*. After 1 hour, most counts are in the bract when organs are detached from the stem but in the pedicellate spikelet when they are attached. Change after 24-hour chase suggests movement of ^14^C from pedicellate to sessile spikelet. Awns remain largely unlabeled. A. *Andropogon schirensis*. C. *Themeda triandra*. B, D. Abaxial epidermis of bract showing rows of stomata (arrows). B. *Andropogon schirensis*. D. *Themeda triandra*, also with prickles (p). Scale = 50 μm.

Figure 7. Summary of biochemical pathways assessed by ^13^C and RNA-seq data. Numbered circles correspond to enzymes whose expression is shown in Figure 5. Colored squares reflect percent average label in ^13^C assays of metabolites. Upper part of figure shows autotrophic metabolism and is enriched in the leaf and PS; lower part of figure shows heterotrophic metabolism, enriched in the SS.

## Supplemental Figures

Figure S1. Spikelet pair and inflorescence structure in *Andropogon schirensis* (A, B, C), and *Themeda triandra* (D, E, F, G). A. Illustration of two spikelet pairs of *A. schirensis*, marked by dotted lines. Each pair is composed of a sessile spikelet, which includes a bisexual flower and bears the seed, and a pedicellate spikelet, which is staminate. The sessile spikelet bears a twisted awn from the lemma (floral bract). B. Spikelet pair of *A. schirensis*. C. Inflorescence of *A. schirensis*, showing two branches, each bearing 9-10 spikelet pairs. D. Spikelet pair of *T. triandra*, showing the dark indurate sessile spikelet, with two greenish pedicellate spikelets behind. One of the pedicellate spikelets is terminal on the short branch, so the three spikelets represent a pair plus a terminal spikelet that is morphologically identical to the pedicellate spikelet. E. Illustration of inflorescence structure in *T. triandra*, with three spikelet pairs, marked by dotted lines, and a terminal spikelet that is morphologically identical to the pedicellate spikelet. Spikelets in the proximal two pairs are all staminate; the distal pair includes a seed-bearing sessile spikelet and a staminate pedicellate spikelet. The sessile spikelet bears a twisted awn from the lemma (floral bract). F. Proximal spikelet pairs of *T. triandra*. All four proximal spikelets are staminate. G. Inflorescence branch of *T. triandra*, showing the spikelet complex as in D and E, subtended by a leaf-like bract. ss, sessile spikelet; ps, pedicellate spikelet; sts, staminate sessile spikelet. Scale bars B, D, F = 1 mm; C = 1 cm; G = 5 mm.

Figure S2. *Andropogon schirensis*. A. ^14^C results. Percent dpm for each organ after 1 hour exposure to ^14^C with organs removed from the axis (detached), inflorescence intact (attached), or after 24-hour chase. Plot includes mean percentages and standard deviations, (n=4). Values are similar after 1 hour, whether organs are attached or lying on filter paper. Change after 24-hour chase suggests movement of ^14^C from PS to SS. Awn is largely unlabeled. B. Abaxial epidermis of pedicellate spikelet showing rows of stomata (arrows), bicellular microhairs (m), and prickles (p). C. Abaxial epidermis of sessile spikelet showing no stomata, but silica bodies (sb). D. Awn showing no stomata, but sparse macrohairs. Scale = 50 μm; note that B and C are more highly magnified than D.

Figure S3. *Themeda triandra*. A. ^14^C results. Percent dpm for each organ after 1 hour exposure to ^14^C with organs removed from the axis (detached), inflorescence intact (attached), or after 24-hour chase. Plot includes mean percentages and standard deviations, (n=3). Values are similar after 1 hour, whether organs are attached or lying on filter paper. Change after 24-hour chase suggests movement of ^14^C from pedicellate to sessile spikelet. Awn is largely unlabeled. B. Abaxial epidermis of pedicellate spikelet showing rows of stomata (arrows), bicellular microhairs (m), and prickles (p). C. Abaxial epidermis of sessile spikelet showing no stomata, but large pits (pit) and macrohairs (mac). D. Awn showing no stomata, but long macrohairs. Scale = 50 μm; note that B and C are more highly magnified than D.

Figure S4. PCA of all isotopologues for ^13^C experiment, showing similarity of results between species. Awn and sessile spikelet are not significantly different for most metabolites, whereas pedicellate spikelet is significantly different from the other two organs. Values at 30 and 60 seconds are not significantly different for most metabolites but 300 second time point is distinct. A. *Sorghum bicolor*. B. *Themeda triandra*.

Figure S5. Isotopologues of 2-phosphoglycolate over time. 2-PG is labeled only in the pedicellate spikelet by the 300 second time point. Points are mean percentages, bars are standard deviations (n=3).

Figure S6. Cross section of glumes of sorghum spikelets, showing vein spacing. A. Pedicellate spikelet. Veins (indicated by arrows) are separated by three to five mesophyll cells, generally more than in the leaf. B. Sessile spikelet. Veins are small and distantly separated. Abaxial side is largely made up of thick-walled cells with no vascularization. Scale = 20 μm.

Figure S7. *Themeda triandra*, PCA of ^13^C labeled metabolites, average labeling. Organs are distinguished by color, time points by shape. Awn, awn; SS, sessile spikelet; PS, pedicellate spikelet; Bract, bract. 30, values at 30 sec of labeling; 60, values at 60 sec of labeling; 300, values at 300 sec of labeling. A. Spikelets and awn only. Values for awn and sessile spikelet are not significantly different at any time point, whereas values for the pedicellate spikelet are significantly different, with the greatest variation in labeling at 5 minutes (300 sec). B. Spikelets, awn, and bract. Values for awn, sessile spikelet and pedicellate spikelet the same as those in A. Awn and sessile spikelet are not significantly different at any time point, whereas values for the pedicellate spikelet are significantly different from awn and sessile spikelet, and bract different from the other three organs, with the greatest variation in labeling at 5 minutes (300 sec).

Figure S8. *Sorghum bicolor*, PCA of gene expression data for the 1441 select metabolic genes, showing distinct sets of transcripts for each organ. Values for each replicate experiment are similar. SS, sessile spikelet; PS, pedicellate spikelet.

Figure S9. *Sorghum bicolor*, heat map of the 922 DE metabolic genes, by organ. These are a subset of the 1441 genes depicted in the PCA in Figure S8. Colors reflect scaled z-scores of log_2_-normalized expression values. SS, sessile spikelet; PS, pedicellate spikelet.

Figure S10. *Themeda triandra*, PCA of gene expression data for the 769 select metabolic genes, showing distinct sets of transcripts for each organ. Values for each replicate experiments are similar, albeit with outliers for awn and bract. SS, sessile spikelet; PS, pedicellate spikelet.

Figure S11. *Themeda triandra*. Heat map of 322 DE metabolic genes, by organ. These are a subset of the 769 genes depicted in the PCA in Figure S10. Colors reflect scaled z-scores of log_2_-normalized expression values. SS, sessile spikelet; PS, pedicellate spikelet. SS, sessile spikelet; PS, pedicellate spikelet.

Figure S12. *<u>Themeda triandra</u>*. Relative expression of 24 genes encoding biosynthetic enzymes immediately responsible for producing the metabolites labeled with ^13^C, a subset extracted from full set of 322 DE metabolic genes in Fig. S11. Colors reflect scaled z-scores of log_2_-normalized expression values. Labels of genes indicate enzyme name, biochemical process, and subcellular localization. SS, sessile spikelet; PS, pedicellate spikelet; cy, cytosolic localized; ch, chloroplast localized; mi, mitochondrial localized; OA/AA, organic acid/amino acid metabolism. *transcripts also DE in *Sorghum bicolor;* see Figure 5. 1) ASPAT, aspartate aminotransferase, Sobic.009G149400.1; 2) *NADP–MDH, NADP-malate dehydrogenase, Sobic.007G166300.1; 3) *G1P-AdT, glucose-1-phosphate adenyltransferase Sobic.002G160400.1; 4) *TPI, triose phosphate isomerase, Sobic.003G072300.2; 5) SuSY,_sucrose synthase_Sobic.010G072300.1; 6) ASNS, asparagine synthetase_Sobic.010G110000.1; 7) GAPDH, glyceraldehyde 3-phosphate dehydrogenase, Sobic.006G105900.1; 8) *F16BP aldo, fructose-1,6-bis-phosphate aldolase, Sobic.005G056400.1; 9) F16BPase, fructose-1,6-bisphosphatase, Sobic.001G425400.1; 10) PRK, phosphoribulokinase, Sobic.006G200800.1; 11) *Starch synthase, Sobic.004G238600.1; 12) ASPAT, aspartate aminotransferase, Sobic.004G331700.1; 13) G1P AdT, G1P–adenyltransferase, Sobic.002G088600.2; 14) GAPDH, glyceraldehyde 3-phosphate dehydrogenase, Sobic.009G016700.1; 15) ASPAT, aspartate aminotransferase, Sobic.004G104700.1; 16) UTP-G1P UdT, UTP–G1P uridyltransferase, Sobic.004G013500.1; 17) *SuSY, sucrose phosphate synthase, Sobic.004G068400.1; 18) UTP-G1P UdT, UTP–G1P uridyltransferase, Sobic.010G251200.1; 19) PEPCK, phosphoenol pyruvate carboxykinasae, Sobic.001G432800.1; 20) ASNS, asparagine synthetase, Sobic.001G406800.1; 21) PPI, phosphopentose isomerase, Sobic.005G033000.1; 22) GS, glutamine synthetase, Sobic.001G451500.1; 23) *PGI, phosphoglucoisomerase, Sobic.002G230600.1; 24) *S17BPase, sedoheptulose 1,7-bisphosphatase, Sobic.003G359100.1.

Figure S13. Spikelet pairs from the eight accessions used for spikelet removal experiments. A. SAP-170 (PI 597971). B. BTx623 (PI 564163). C. Combine Hegari (PI 659691). D SAP-257 (656099). E. SAP-51 (655995). F. Jola Nandyal (534021). G. SAP-15 (656014), H. SO85 (PI 534096). Arrow indicates pedicellate spikelet. All spikelets to the same scale. Scale bar = 1 mm.

Figure S14. Average seed weight (mg) from inflorescences with pedicellate spikelets untouched (on) and removed (off). A. Combined results of both removal experiments, showing 5.2% reduction in average weight with spikelet removal (p=0.0197). B. Average seed weight for each accession in experiment 1. Representative spikelet pair for each accession shown in Figure S13. * = p <0.05.

