## Supplemental.Figures.and.Tables for "Sterile spikelets assimilate carbon in sorghum and related grasses"

Table S1. *Sorghum bicolor*, $^{14}$C data and analysis of variance. Detached=spikelets and awns removed from inflorescence. Attached=spikelets and awns on an intact inflorescence. Both detached and attached spikelets were given a one-hour pulse of $^{14}$C. Chase= spikelets and awns on an intact inflorescence, exposed to $^{14}$C for one hour, and then allowed to photosynthesize in air for another 24 hours. PS=pedicellate spikelet; SS=sessile spikelet; diff=difference between mean values; lwr=minus one standard deviation; upr=plus one standard deviation. Significance codes: ≤0.0001, ***; ≤ 0.001,**; ≤ 0.01,*; ≤ 0.05, . NS=not significant.

A. **Decays per minute per mg (dpm/mg) and percent dpm/mg**.

| | | **Rep1** | | **Rep2** | | **Rep3** | | **mean** |
|---|---|---|---|---|---|---|---|---|
| **treatment** | **organ** | **Dpm/mg** | **%** | **Dpm/mg** | **%** | **Dpm/mg** | **%** | **%** |
| **detached** | **awn** | 3 | 0.16 | 12 | 0.47 | 6 | 0.40 | 0.3 |
| | **SS** | 76 | 4.16 | 224 | 8.75 | 101 | 6.78 | 6.6 |
| | **PS** | 1748 | 95.68 | 2328 | 90.78 | 1383 | 92.82 | 93.1 |
| **attached** | **Awn** | 23 | 0.37 | 69 | 1.24 | 16 | 0.00 | 0.5 |
| | **SS** | 365 | 5.87 | 1091 | 19.51 | 355 | 4.00 | 9.8 |
| | **PS** | 5828 | 93.76 | 4432 | 79.25 | 7749 | 95.00 | 89.3 |
| **chase** | **Awn** | 62 | 2.91 | 158 | 4.05 | 63 | 2.00 | 3.0 |
| | **SS** | 600 | 28.20 | 1480 | 37.83 | 496 | 19.00 | 28.3 |
| | **PS** | 1466 | 68.89 | 2274 | 58.12 | 2110 | 79.00 | 68.7 |

**B. Main effects**

| | **Pr(>F)** | |
|---|---|---|
| Organ | 5.17e-16 | *** |
| Treatment | 0.999 | NS |
| Organ * treatment | 7.06e-05 | *** |

**C. Comparisons of organs**

| | **diff** | **lwr** | **upr** | **p adj** | |
|---|---|---|---|---|---|
| **PS-awn** | 82.41 | 74.81 | 90.01 | 0.00000 | *** |
| **SS-awn** | 13.61 | 6.01 | 21.21 | 0.00066 | *** |
| **SS-PS** | 68.80 | -76.40 | -61.2 | 0.00000 | *** |

**D. Organ x treatment interactions. Significant values in boldface.**

| | **diff** | **lwr** | **upr** | **p adj** |
|---|---|---|---|---|
| Within organs, between treatments | | | | |
| ss:detached-ss:attached | -3.23 | -21.30 | 14.84 | 0.99915 |
| **ss:detached-ss:chase** | **-21.78** | **-39.85** | **-3.71** | **0.01169** |
| **ss:chase-ss:attached** | **18.55** | **0.48** | **36.62** | **0.04** |
| ps:detached-ps:attached | 3.76 | -14.31 | 21.82 | 0.99756 |
| **ps:detached-ps:chase** | **24.42** | **6.36** | **42.49** | **0.00404** |
| **ps:chase-ps:attached** | **-20.67** | **-38.73** | **-2.60** | **0.01820** |

| | | | | |
|---|---|---|---|---|
| awn:detached-awn:attached | -0.19 | -18.26 | 17.87 | 1.00000 |
| awn:detached-awn:chase | -2.64 | -20.71 | 15.42 | 0.99980 |
| awn:chase-awn:attached | 2.46 | -15.62 | 20.52 | 0.99989 |
| Within treatments, between organs | | | | |
| **ps:attached-awn:attached** | **88.80** | **70.73** | **106.87** | **0.00000** |
| **ss:attached-ps:attached** | **-79.54** | **-97.61** | **-61.48** | **0.00000** |
| ss:attached-awn:attached | 9.26 | -8.81 | 27.32 | 0.68462 |
| **ps:detached-awn:detached** | **92.75** | **74.68** | **110.82** | **0.00000** |
| **ss:detached-ps:detached** | **-86.53** | **-104.60** | **-68.46** | **0.00000** |
| ss:detached-awn:detached | 6.22 | -11.85 | 24.29 | 0.94479 |
| **ps:chase-awn:chase** | **65.68** | **47.62** | **83.75** | **0.00000** |
| **ss:chase-ps:chase** | **-40.33** | **-58.39** | **-22.26** | **0.00001** |
| **ss:chase-awn:chase** | **25.36** | **7.29** | **43.42** | **0.00278** |
| Other comparisons | | | | |
| **ps:chase-awn:attached** | **68.13** | **50.07** | **86.20** | **0.00000** |
| **ss:chase-awn:attached** | **27.81** | **9.74** | **45.87** | **0.00104** |
| **ps:detached-awn:attached** | **92.56** | **74.49** | **110.62** | **0.00000** |
| ss:detached-awn:attached | 6.03 | -12.05 | 24.09 | 0.95342 |
| **awn:chase-ps:attached** | **-86.35** | **-104.42** | **-68.28** | **0.00000** |
| **ss:chase-ps:attached** | **-60.99** | **-79.06** | **-42.93** | **0.00000** |
| **awn:detached-ps:attached** | **-88.99** | **-107.06** | **-70.93** | **0.00000** |
| **ss:detached-ps:attached** | **-82.77** | **-100.84** | **-64.71** | **0.00000** |
| awn:chase-ss:attached | -6.81 | -24.87 | 11.26 | 0.91233 |
| **ps:chase-ss:attached** | **58.88** | **40.81** | **76.94** | **0.00000** |
| awn:detached-ss:attached | -9.45 | -27.51 | 8.62 | 0.66242 |
| **ps:detached-ss:attached** | **83.30** | **65.23** | **101.37** | **0.00000** |
| **ps: detached-awn:chase** | **90.11** | **72.04** | **108.17** | **0.00000** |
| ss:detached-awn:chase | 3.58 | -14.49 | 21.64 | 0.99826 |
| **awn:detached-ps:chase** | **-68.33** | **-86.39** | **-50.26** | **0.00000** |
| **ss:detached-ps:chase** | **-62.11** | **-80.17** | **-44.04** | **0.00000** |
| **awn:detached-ss:chase** | **-28.00** | **-46.07** | **-9.93** | **0.00097** |
| **ps:detached-ss:chase** | **64.75** | **46.68** | **82.82** | **0.00000** |

Table S2. *Andropogon schirensis*, $^{14}C$ data and analysis of variance. Detached=spikelets and awns removed from inflorescence. Attached=spikelets and awns on an intact inflorescence. Both detached and attached spikelets were given a one-hour pulse of $^{14}C$. Chase=spikelets and awns on an intact inflorescence, exposed to $^{14}C$ for one hour, and then allowed to photosynthesize in air for another 24 hours. PS=pedicellate spikelet; SS=sessile spikelet; diff=difference between mean values; lwr=minus one standard deviation; upr=plus one standard deviation. Significance codes: ≤0.0001, ***; ≤ 0.001,**; ≤ 0.01,*; ≤ 0.05, . NS=not significant.

A. **Decays per minute per mg (dpm/mg) and percent dpm/mg**.

| | | **Rep1** | | **Rep2** | | **Rep3** | | **mean** |
|---|---|---|---|---|---|---|---|---|
| **treatment** | **organ** | **Dpm/mg** | **%** | **Dpm/mg** | **%** | **Dpm/mg** | **%** | **%** |
| **detached** | **Awn** | 37 | 0.24 | 34 | 0.80 | 16 | 1.05 | 0.7 |
| | **SS** | 3487 | 22.52 | 809 | 19.01 | 303 | 19.83 | 20.4 |
| | **PS** | 11958 | 77.24 | 3413 | 80.19 | 1209 | 79.12 | 78.8 |
| **attached** | **Awn** | 40 | 0.10 | 10 | 0.03 | 23 | 0.12 | 0.1 |
| | **SS** | 7953 | 20.61 | 5621 | 17.87 | 4303 | 23.10 | 20.5 |
| | **PS** | 30587 | 79.28 | 25825 | 82.10 | 14300 | 76.77 | 79.4 |
| **chase** | **Awn** | 1731 | 7.01 | 152 | 0.97 | 761 | 5.89 | 4.6 |
| | **SS** | 13859 | 56.12 | 5640 | 36.07 | 5391 | 41.73 | 44.6 |
| | **PS** | 9105 | 36.87 | 9846 | 62.96 | 6768 | 52.38 | 50.7 |

**B. Main effects**

| | **Pr(>F)** | |
|---|---|---|
| Organ | 1.26e-14 | *** |
| Treatment | 1 | NS |
| Organ * treatment | 1.67e-06 | *** |

**C. Comparisons of organs**

| | **diff** | **lwr** | **upr** | **p adj** | |
|---|---|---|---|---|---|
| **PS-awn** | 67.86 | 60.81 | 74.91 | 0 | *** |
| **SS-awn** | 26.74 | 19.69 | 33.79 | 0 | *** |
| **SS-PS** | -41.12 | -48.17 | 34.07 | 0 | *** |

**D. Organ x treatment interactions. Significant values in boldface.**

| | **diff** | **lwr** | **upr** | **p adj** |
|---|---|---|---|---|
| Within organs, between treatments | | | | |
| ss:detached-ss:attached | -0.08 | -16.85 | 16.69 | 1.00000 |
| **ss:detached-ss:chase** | **-24.19** | **-40.96** | **-7.42** | **0.00209** |
| **ss:chase-ss:attached** | **24.11** | **7.34** | **40.88** | **0.00217** |

| | | | | |
|---|---|---|---|---|
| ps:detached-ps:attached | -0.53 | -17.30 | 16.23 | 1.00000 |
| **ps:detached-ps:chase** | **28.11** | **11.34** | **44.88** | **0.00039** |
| **ps:chase-ps:attached** | **-28.64** | **-45.41** | **-11.88** | **0.00032** |
| awn:detached-awn:attached | 0.61 | -16.16 | 17.37 | 1.00000 |
| awn:detached-awn:chase | -3.93 | -20.70 | 12.84 | 0.99452 |
| awn:chase-awn:attached | 4.54 | -12.23 | 21.30 | 0.98628 |
| Within treatments, between organs | | | | |
| **ps:attached-awn:attached** | **79.30** | **62.53** | **96.06** | **0.00000** |
| **ss:attached-ps:attached** | **-58.85** | **-75.62** | **-42.09** | **0.00000** |
| **ss:attached-awn:attached** | **20.44** | **3.68** | **37.21** | **0.01058** |
| **ps:detached-awn:detached** | **78.16** | **61.39** | **94.92** | **0.00000** |
| **ss:detached-ps:detached** | **-58.40** | **-75.17** | **-41.63** | **0.00000** |
| **ss:detached-awn:detached** | **19.76** | **2.99** | **36.52** | **0.01422** |
| **ps:chase-awn:chase** | **46.12** | **29.35** | **62.88** | **0.00000** |
| ss:chase-ps:chase | -6.10 | -22.87 | 10.67 | 0.92640 |
| **ss:chase-awn:chase** | **40.02** | **23.25** | **56.78** | **0.00000** |
| Other comparisons | | | | |
| **awn:chase-ps:attached** | **-74.76** | **-91.53** | **-57.99** | **0.00000** |
| awn:chase-ss:attached | -15.91 | -32.67 | 0.86 | 0.07072 |
| **awn:detached-ps:attached** | **-78.69** | **-95.46** | **-61.92** | **0.00000** |
| **awn:detached-ps:chase** | **-50.05** | **-66.81** | **-33.28** | **0.00000** |
| **awn:detached-ss:attached** | **-19.84** | **-36.60** | **-3.07** | **0.01374** |
| **awn:detached-ss:chase** | **-43.95** | **-60.71** | **-27.18** | **0.00000** |
| **ps:detached-awn:chase** | **74.23** | **57.46** | **90.99** | **0.00000** |
| **ps:chase-awn:attached** | **50.65** | **33.89** | **67.42** | **0.00000** |
| **ps:chase-ss:attached** | **30.21** | **13.44** | **46.98** | **0.00017** |
| **ps:detached-awn:attached** | **78.76** | **62.00** | **95.53** | **0.00000** |
| **ps:detached-ss:attached** | **58.32** | **41.55** | **75.09** | **0.00000** |
| **ps:detached-ss:chase** | **34.21** | **17.44** | **50.98** | **0.00003** |
| **ss:chase-awn:attached** | **44.55** | **27.79** | **61.32** | **0.00000** |
| **ss:chase-ps:attached** | **-34.74** | **-51.51** | **-17.98** | **0.00003** |
| **ss:detached-awn:attached** | **20.36** | **3.60** | **37.13** | **0.01095** |
| ss:detached-awn:chase | 15.83 | -0.94 | 32.59 | 0.07301 |
| **ss:detached-ps:attached** | **-58.93** | **-75.70** | **-42.17** | **0.00000** |
| **ss:detached-ps:chase** | **-30.29** | **-47.06** | **-13.52** | **0.00016** |

Table S3. *Themeda triandra*, $^{14}$C data and analysis of variance. Detached=spikelets and awns removed from inflorescence. Attached=spikelets and awns on an intact inflorescence. Both detached and attached spikelets were given a one-hour pulse of $^{14}$C. Chase=spikelets and awns on an intact inflorescence, exposed to $^{14}$C for one hour, and then allowed to photosynthesize in air for another 24 hours. PS=pedicellate spikelet; SS=sessile spikelet; diff=difference between mean values; lwr=minus one standard deviation; upr=plus one standard deviation. Significance codes: ≤0.0001, ***; ≤ 0.001,**; ≤ 0.01,*; ≤ 0.05, . NS=not significant.

A. **Decays per minute per mg (dpm/mg) and percent dpm/mg**.

| | | **Rep1** | | **Rep2** | | **Rep3** | | **mean** |
|---|---|---|---|---|---|---|---|---|
| **treatment** | **organ** | **Dpm/mg** | **%** | **Dpm/mg** | **%** | **Dpm/mg** | **%** | **%** |
| **detached** | **Awn** | 22 | 0.75 | 12 | 1.44 | 14 | 0.45 | 0.9 |
| | **SS** | 32 | 1.09 | 29 | 3.50 | 43 | 1.37 | 2.0 |
| | **PS** | 2870 | 98.15 | 788 | 95.05 | 3079 | 98.18 | 97.1 |
| **attached** | **Awn** | 69 | 0.42 | 35 | 0.35 | 225 | 1.89 | 0.9 |
| | **SS** | 1761 | 10.74 | 1006 | 10.16 | 2139 | 17.98 | 12.9 |
| | **PS** | 14560 | 88.84 | 8860 | 89.49 | 9532 | 80.13 | 86.1 |
| **chase** | **Awn** | 3589 | 15.19 | 3476 | 18.82 | 1437 | 10.12 | 14.7 |
| | **SS** | 8749 | 37.04 | 9823 | 53.17 | 9023 | 63.52 | 51.2 |
| | **PS** | 11284 | 47.77 | 5175 | 28.12 | 3746 | 26.37 | 34.0 |

B. **Main effects**

| | **Pr(>F)** | |
|---|---|---|
| **Organ** | 6.89e-14 | *** |
| **Treatment** | 1 | NS |
| **Organ * treatment** | 1.99e-10 | *** |

C. **Comparisons of organs**

| | **diff** | **lwr** | **upr** | **p adj** | |
|---|---|---|---|---|---|
| **PS-awn** | 66.92 | 59.00 | 74.85 | 0.00000 | *** |
| **SS-awn** | 16.57 | 8.65 | 24.50 | 0.00013 | ** |
| **SS-PS** | -50.35 | -58.28 | -42.43 | 0.00000 | *** |

D. **Organ x treatment interactions. Significant values in boldface.**

| | **diff** | **lwr** | **upr** | **p adj** |
|---|---|---|---|---|
| Within organs, between treatments | | | | |
| ss:detached-ss:attached | -10.96 | -29.81 | 7.88 | 0.53861 |
| **ss:detached-ss:chase** | **-49.25** | **-68.10** | **-30.41** | **0.00000** |
| **ss:chase-ss:attached** | **38.29** | **19.45** | **57.13** | **0.00004** |
| ps:detached-ps:attached | 10.93 | -7.91 | 29.78 | 0.54195 |
| **ps:detached-ps:chase** | **63.04** | **44.20** | **81.89** | **0.00000** |
| **ps:chase-ps:attached** | **-52.11** | **-70.95** | **-33.27** | **0.00000** |
| awn:detached-awn:attached | 0.01 | -18.84 | 18.85 | 1.00000 |

| | | | | |
|---|---|---|---|---|
| awn:detached-awn:chase | -13.83 | -32.67 | 5.02 | 0.26375 |
| awn:chase-awn:attached | 13.83 | -5.01 | 32.68 | 0.26324 |
| Within treatments, between organs | | | | |
| **ps:attached-awn:attached** | **85.26** | **66.42** | **104.11** | **0.00000** |
| **ss:attached-ps:attached** | **-73.19** | **-92.03** | **-54.34** | **0.00000** |
| ss:attached-awn:attached | 12.08 | -6.77 | 30.92 | 0.41966 |
| **ps:detached-awn:detached** | **96.19** | **77.35** | **115.03** | **0.00000** |
| **ss:detached-ps:detached** | **-95.08** | **-113.93** | **-76.24** | **0.00000** |
| ss:detached-awn:detached | 1.11 | -17.74 | 19.95 | 1.00000 |
| **ps:chase-awn:chase** | **19.32** | **0.48** | **38.16** | **0.04201** |
| ss:chase-ps:chase | 17.21 | -1.63 | 36.06 | 0.08924 |
| **ss:chase-awn:chase** | **36.53** | **17.69** | **55.38** | **0.00007** |
| Other comparisons | | | | |
| **awn:chase-ps:attached** | **-71.43** | **-90.27** | **-52.59** | **0.00000** |
| awn:chase-ss:attached | 1.76 | -17.09 | 20.60 | 0.99999 |
| **awn:detached-ps:attached** | **-85.26** | **-104.10** | **-66.41** | **0.00000** |
| **awn:detached-ps:chase** | **-33.15** | **-51.99** | **-14.30** | **0.00022** |
| **awn:detached-ss:attached** | -12.07 | -30.91 | 6.77 | 0.42034 |
| **awn:detached-ss:chase** | **-50.36** | **-69.20** | **-31.52** | **0.00000** |
| **ps:detached-awn:chase** | **82.36** | **63.52** | **101.21** | **0.00000** |
| **ps:chase-awn:attached** | **33.15** | **14.31** | **52.00** | **0.00022** |
| **ps:chase-ss:attached** | **21.08** | **2.23** | **39.92** | **0.02182** |
| **ps:detached-awn:attached** | **96.20** | **77.35** | **115.04** | **0.00000** |
| **ps:detached-ss:attached** | **84.12** | **65.28** | **102.96** | **0.00000** |
| **ps:detached-ss:chase** | **45.83** | **26.99** | **64.67** | **0.00000** |
| **ss:chase-awn:attached** | **50.37** | **31.52** | **69.21** | **0.00000** |
| **ss:chase-ps:attached** | **-34.90** | **-53.74** | **-16.05** | **0.00012** |
| ss:detached-awn:attached | 1.11 | -17.73 | 19.96 | 1.00000 |
| ss:detached-awn:chase | -12.72 | -31.56 | 6.12 | 0.35716 |
| **ss:detached-ps:attached** | **-84.15** | **-102.99** | **-65.31** | **0.00000** |
| **ss:detached-ps:chase** | **-32.04** | **-50.88** | **-13.20** | **0.00033** |

Table S4. *Sorghum bicolor*, $^{13}$C data, analysis of variance. Average labeling of each metabolite. ADPG, ADP-glucose; ASP, aspartate; F6P, fructose-6-phosphate; G6P, glucose-6=phosphate; MAL, malate; P5P, pentose-5 phosphates; PGA, phosphoglycerate; PYR, pyruvate; TP, triose phosphate; UDPG, UDP-glucose. Significance codes: ≤0.0001, ***; ≤ 0.001,**; ≤ 0.01,*; ≤ 0.05, .; NS, non-significant.

A. Main effects.

| | **ADPG** | | **ASP** | | **F6P** | | **G6P** | | **MAL** | |
|---|---|---|---|---|---|---|---|---|---|---|
| **organ** | 2.81E-07 | *** | 0.001648 | * | 2.11E-12 | *** | 3.71E-14 | *** | 4.58E-08 | *** |
| **time** | 1.12E-06 | *** | 0.172662 | NS | 3.94E-11 | *** | 4.27E-14 | *** | 0.000473 | ** |
| **organ*time** | 3.27E-07 | *** | 0.396077 | NS | 1.96E-11 | *** | 1.51E-14 | *** | 0.001805 | * |

| | **P5P** | | **PGA** | | **PYR** | | **TP** | | **UDPG** | |
|---|---|---|---|---|---|---|---|---|---|---|
| **organ** | 5.98E-08 | *** | 1.67E-09 | *** | 0.003704 | * | 6.23E-11 | *** | 1.71E-10 | *** |
| **time** | 0.001917 | * | 1.11E-07 | *** | 3.53E-05 | *** | 3.85E-10 | *** | 1.28E-10 | *** |
| **organ*time** | 0.005079 | * | 3.23E-07 | *** | 0.000818 | *** | 1.05E-08 | *** | 4.93E-11 | *** |

B. Organ and time. Tukey's honestly significant difference.

| | **ADPG** | | **ASP** | | **F6P** | | **G6P** | | **MAL** | |
|---|---|---|---|---|---|---|---|---|---|---|
| **SS-awn** | 0.934858 | NS | 0.977818 | NS | 0.999815 | NS | 0.982508 | NS | 0.881856 | NS |
| **PS-awn** | 1.82E-06 | *** | 0.003267 | * | 1.34E-11 | *** | 2.50E-13 | *** | 1.57E-07 | *** |
| **PS-SS** | 9.64E-07 | *** | 0.005078 | * | 1.37E-11 | *** | 2.84E-13 | *** | 3.54E-07 | *** |
| **60-30** | 0.828695 | NS | 0.733657 | NS | 0.388814 | NS | 0.347807 | NS | 0.303661 | NS |
| **300-30** | 2.89E-06 | *** | 0.152875 | NS | 1.16E-10 | *** | 1.89E-13 | *** | 0.000396 | ** |
| **300-60** | 8.91E-06 | *** | 0.470255 | NS | 5.99E-10 | *** | 5.35E-13 | *** | 0.010953 | . |

| | **P5P** | | **PGA** | | **PYR** | | **TP** | | **UDPG** | |
|---|---|---|---|---|---|---|---|---|---|---|
| **SS-awn** | 0.440988 | NS | 0.950910 | NS | 0.207205 | NS | 0.365458 | NS | 0.280613 | NS |
| **PS-awn** | 1.17E-07 | *** | 1.18E-08 | *** | 0.105272 | NS | 1.74E-10 | *** | 3.28E-09 | *** |
| **PS-SS** | 1.01E-06 | *** | 7.55E-09 | *** | 0.002660 | * | 9.91E-10 | *** | 4.08E-10 | *** |
| **60-30** | 0.700554 | NS | 0.283460 | NS | 0.954232 | NS | 0.047993 | NS | 0.936898 | NS |
| **300-30** | 0.002204 | * | 1.71E-07 | *** | 0.000170 | ** | 5.45E-10 | *** | 6.18E-10 | *** |
| **300-60** | 0.012874 | . | 2.76E-06 | *** | 9.15E-05 | *** | 1.95E-08 | *** | 9.66E-10 | *** |

C. Organ x time interactions. Significant values in boldface. Only rows containing at least one significant value are included.

| | **ADPG** | **ASP** | **F6P** | **G6P** | **MAL** |
|---|---|---|---|---|---|
| PS:60-A:30 | 0.978254 | 0.492821 | 0.053432 | 0.198622 | **0.006711** |
| SS:300-A:30 | 0.999989 | 0.999878 | 0.771386 | 0.482259 | 0.898013 |
| PS:300-A:30 | **3.36E-08** | **0.030255** | **3.35E-13** | **2.55E-14** | **8.48E-07** |
| PS:60-SS:30 | 0.609531 | 0.487696 | **0.013744** | 0.081440 | **0.008855** |
| PS:300-SS:30 | **1.00E-08** | **0.029725** | **2.21E-13** | **2.55E-14** | **1.05E-06** |
| PS:300-PS:30 | **2.32E-08** | 0.182989 | **8.70E-13** | **2.55E-14** | **6.68E-05** |
| PS:60-A:60 | 0.895684 | 0.543686 | 0.069777 | 0.227494 | **0.021173** |
| PS:300-A:60 | **2.09E-08** | **0.035932** | **3.65E-13** | **2.55E-14** | **2.12E-06** |
| PS:60-SS:60 | 0.651851 | 0.527887 | **0.021419** | 0.095260 | **0.010367** |

| | | | | | |
|---|---|---|---|---|---|
| PS:300-SS:60 | **1.10E-08** | **0.034084** | **2.52E-13** | **2.55E-14** | **1.19E-06** |
| A:300-PS:60 | 0.865153 | 0.549651 | 0.101230 | 0.606306 | **0.012927** |
| PS:300-PS:60 | **1.46E-07** | 0.771567 | **3.65E-12** | **2.71E-14** | **0.003724** |
| PS:300-A:300 | **1.88E-08** | **0.036652** | **4.13E-13** | **2.58E-14** | **1.42E-06** |
| PS:300-SS:300 | **5.50E-08** | 0.077426 | **9.60E-13** | **2.64E-14** | **8.44E-06** |

| | **P5P** | **PGA** | **PYR** | **TP** | **UDPG** |
|---|---|---|---|---|---|
| PS:60-A:30 | **0.021449** | **0.027167** | 0.944297 | **0.002130** | 0.999984 |
| SS:300-A:30 | 0.825546 | 0.825340 | 0.999986 | **0.021430** | 1.000000 |
| PS:300-A:30 | **2.06E-06** | **8.93E-10** | **0.001343** | **7.52E-12** | **5.34E-12** |
| PS:60-SS:30 | **0.028069** | **0.017812** | 0.969371 | **0.002251** | **0.351023** |
| SS:300-SS:30 | 0.883156 | 0.714977 | 0.603403 | **0.022624** | 0.551484 |
| PS:300-SS:30 | **2.58E-06** | **7.05E-10** | **1.56E-05** | **7.71E-12** | **1.15E-12** |
| PS:300-PS:30 | **0.000588** | **5.63E-09** | **3.52E-05** | **5.63E-11** | **2.61E-12** |
| PS:60-A:60 | **0.028503** | 0.097858 | 0.999998 | **0.011346** | 0.936281 |
| PS:300-A:60 | **2.61E-06** | **1.93E-09** | **6.91E-05** | **1.63E-11** | **2.62E-12** |
| PS:60-SS:60 | 0.060451 | **0.011441** | 0.997284 | **0.007977** | 0.468820 |
| PS:300-SS:60 | **5.01E-06** | **5.53E-10** | **2.93E-05** | **1.38E-11** | **1.34E-12** |
| A:300-PS:60 | **0.016981** | 0.119157 | 0.953918 | **0.035720** | 0.999646 |
| PS:300-PS:60 | **0.003559** | **1.27E-07** | **0.000120** | **1.64E-09** | **7.42E-12** |
| PS:300-A:300 | **1.71E-06** | **2.19E-09** | **0.001232** | **2.89E-11** | **4.55E-12** |
| PS:300-SS:300 | **3.09E-05** | **5.63E-09** | **0.000642** | **4.55E-10** | **5.38E-12** |

Table S5. *Sorghum bicolor*, $^{13}C$ data, analysis of variance. Isotopomers of each metabolite analyzed separately. ADPG, ADP-glucose; ASP, aspartate; F6P, fructose-6-phosphate; G6P, glucose-6=phosphate; MAL, malate; P5P, pentose-5 phosphates; PGA, phosphoglycerate; PYR, pyruvate; TP, triose phosphate; UDPG, UDP-glucose. M0 = percentage of metabolite with no labeled carbons; M1, percentage of metabolite with one labeled carbon; M2, percentage with two labeled carbons; M3, percentage with three labeled carbons. Significance codes: ≤0.0001, ***; ≤ 0.001,**; ≤ 0.01,*; ≤ 0.05, .; NS, non-significant.

A. Main effects

| | | **M0** | | **M1** | | **M2** | | **M3** | |
|---|---|---|---|---|---|---|---|---|---|
| **ADPG** | **organ** | 4.09E-07 | *** | 0.002279 | * | 0.000243 | ** | 8.89E-07 | *** |
| | **time** | 3.00E-05 | *** | 0.134129 | NS | 0.184621 | NS | 3.82E-07 | *** |
| | **organ*time** | 1.22E-05 | *** | 0.393704 | NS | 0.000526 | ** | 1.40E-06 | *** |
| **ASP** | **organ** | 0.000535 | ** | 9.38E-05 | *** | 0.017530 | . | 0.022028 | . |
| | **time** | 0.317841 | NS | 0.427623 | NS | 0.059559 | NS | 0.019047 | . |
| | **organ*time** | 0.643006 | NS | 0.706122 | NS | 0.139083 | NS | 0.043625 | . |
| **F6P** | **organ** | 5.67E-12 | *** | 6.35E-10 | *** | 6.51E-10 | *** | 4.24E-13 | *** |
| | **time** | 7.27E-09 | *** | 0.073813 | NS | 8.31E-07 | *** | 1.97E-12 | *** |
| | **organ*time** | 1.92E-08 | *** | 0.031743 | . | 4.08E-06 | *** | 1.36E-12 | *** |
| **G6P** | **organ** | 2.39E-12 | *** | 3.31E-07 | *** | 4.52E-12 | *** | 9.30E-15 | *** |
| | **time** | 1.65E-11 | *** | 0.000512 | ** | 5.33E-12 | *** | 4.21E-15 | *** |
| | **organ*time** | 2.69E-11 | *** | 0.043564 | . | 6.62E-12 | *** | 1.66E-15 | *** |
| **MAL** | **organ** | 6.87E-08 | *** | 6.12E-08 | *** | 2.54E-06 | *** | 4.60E-09 | *** |
| | **time** | 0.017252 | . | 0.139876 | NS | 1.92E-05 | *** | 5.57E-09 | *** |
| | **organ*time** | 0.090347 | NS | 0.460770 | NS | 6.84E-05 | *** | 1.12E-08 | *** |
| **P5P** | **organ** | 1.58E-08 | *** | 4.67E-07 | *** | 0.000153 | ** | 8.11E-06 | *** |
| | **time** | 0.013016 | . | 0.222652 | NS | 0.105821 | NS | 0.000208 | ** |
| | **organ*time** | 0.133491 | NS | 0.000237 | ** | 0.170909 | NS | 0.000142 | ** |
| **PGA** | **organ** | 5.84E-10 | *** | 1.52E-09 | *** | 1.20E-09 | *** | 7.48E-07 | *** |
| | **time** | 1.78E-06 | *** | 0.005759 | * | 7.35E-09 | *** | 1.44E-07 | *** |
| | **organ*time** | 8.59E-06 | *** | 0.042593 | . | 1.54E-08 | *** | 2.37E-07 | *** |
| **PYR** | **organ** | 0.002262 | * | 0.015527 | . | 0.340756 | NS | 0.000654 | ** |
| | **time** | 1.91E-05 | *** | 0.000278 | ** | 0.073944 | NS | 3.94E-06 | *** |
| | **organ*time** | 0.003301 | * | 0.187843 | NS | 0.090480 | NS | 8.85E-06 | *** |
| **TP** | **organ** | 6.57E-10 | *** | 5.76E-08 | *** | 4.91E-09 | *** | 4.87E-11 | *** |
| | **time** | 1.52E-07 | *** | 0.005435 | * | 5.20E-10 | *** | 3.28E-12 | *** |
| | **organ*time** | 1.60E-05 | *** | 0.204168 | NS | 5.24E-08 | *** | 6.66E-12 | *** |
| **UDPG** | **organ** | 2.00E-08 | *** | 0.000175 | ** | 4.24E-07 | *** | 2.86E-13 | *** |
| | **time** | 6.21E-08 | *** | 0.078223 | NS | 1.78E-06 | *** | 9.98E-14 | *** |
| | **organ*time** | 6.75E-08 | *** | 0.053127 | NS | 1.07E-06 | *** | 3.43E-14 | *** |

B. Organ and time. Tukey's honestly significant difference.

| | | M0 | | M1 | | M2 | | M3 | |
|---|---|---|---|---|---|---|---|---|---|
| **ADPG** | **SS-awn** | 0.859831 | NS | 0.996652 | NS | 0.173908 | NS | 0.999104 | NS |
| | **PS-awn** | 3.14E-06 | *** | 0.004956 | * | 0.010762 | . | 3.77E-06 | *** |
| | **PS-ss** | 1.19E-06 | *** | 0.005874 | * | 0.000182 | ** | 4.07E-06 | *** |
| | **60-30** | 0.430737 | NS | 0.130083 | NS | 0.997156 | NS | 0.620457 | NS |
| | **300-30** | 3.58E-05 | *** | 0.309235 | NS | 0.258078 | NS | 8.08E-07 | *** |
| | **300-60** | 5.22E-04 | *** | 0.855809 | NS | 0.230746 | NS | 4.62E-06 | *** |
| **ASP** | **SS-awn** | 0.973026 | NS | 0.964799 | NS | 0.987983 | NS | 0.970717 | NS |
| | **PS-awn** | 0.001163 | * | 0.000232 | ** | 0.028002 | . | 0.031680 | . |
| | **PS-ss** | 0.001895 | * | 0.000402 | ** | 0.037911 | . | 0.050618 | NS |
| | **60-30** | 0.601271 | NS | 0.420051 | NS | 0.888987 | NS | 0.961063 | NS |
| | **300-30** | 0.293947 | NS | 0.617148 | NS | 0.064222 | NS | 0.026980 | . |
| | **300-60** | 0.837216 | NS | 0.939148 | NS | 0.150813 | NS | 0.046593 | . |
| **F6P** | **SS-awn** | 0.943959 | NS | 0.475197 | NS | 0.791563 | NS | 0.698573 | NS |
| | **PS-awn** | 3.05E-11 | *** | 1.72E-09 | *** | 6.09E-09 | *** | 1.78E-12 | *** |
| | **PS-ss** | 4.38E-11 | *** | 9.16E-09 | *** | 2.42E-09 | *** | 4.04E-12 | *** |
| | **60-30** | 0.201378 | NS | 0.158281 | NS | 0.145234 | NS | 0.219163 | NS |
| | **300-30** | 1.21E-08 | *** | 0.084251322 | NS | 8.71E-07 | *** | 5.38E-12 | *** |
| | **300-60** | 2.05E-07 | *** | 0.935060412 | NS | 3.93E-05 | *** | 3.54E-11 | *** |
| **G6P** | **SS-awn** | 0.894049 | NS | 0.565067 | NS | 0.404245 | NS | 0.700357 | NS |
| | **PS-awn** | 1.21E-11 | *** | 6.63E-07 | *** | 6.39E-11 | *** | 7.99E-14 | *** |
| | **PS-ss** | 1.97E-11 | *** | 4.45E-06 | *** | 1.48E-11 | *** | 1.22E-13 | *** |
| | **60-30** | 0.135805 | NS | 0.154344 | NS | 0.022476 | . | 0.443966 | NS |
| | **300-30** | 3.72E-11 | *** | 0.000365 | ** | 8.68E-12 | *** | 4.91E-14 | *** |
| | **300-60** | 4.00E-10 | *** | 0.024800 | . | 2.43E-10 | *** | 8.13E-14 | *** |
| **MAL** | **SS-awn** | 0.909937 | NS | 0.941420 | NS | 0.838946 | NS | 0.781292 | NS |
| | **PS-awn** | 2.42E-07 | *** | 2.32E-07 | *** | 6.29E-06 | *** | 1.54E-08 | *** |
| | **PS-ss** | 4.95E-07 | *** | 4.09E-07 | *** | 1.92E-05 | *** | 4.35E-08 | *** |
| | **60-30** | 0.210406 | NS | 0.129023 | NS | 0.556553 | NS | 0.782318 | NS |
| | **300-30** | 0.013160 | . | 0.358196 | NS | 2.77E-05 | *** | 1.85E-08 | *** |
| | **300-60** | 0.344482 | NS | 0.798833 | NS | 0.000248 | ** | 5.25E-08 | *** |
| **P5P** | **SS-awn** | 0.039034 | . | 0.001986 | * | 0.025811 | . | 0.918859 | NS |
| | **PS-awn** | 1.62E-08 | *** | 2.79E-07 | *** | 9.96E-05 | *** | 2.14E-05 | *** |
| | **PS-ss** | 1.37E-06 | *** | 0.000825662 | *** | 0.047684 | . | 4.77E-05 | *** |
| | **60-30** | 0.502032 | NS | 0.486248 | NS | 0.571680 | NS | 0.904291 | NS |
| | **300-30** | 0.010810 | . | 0.816176 | NS | 0.088622 | NS | 0.000405 | ** |
| | **300-60** | 0.108080 | NS | 0.204634 | NS | 0.450618 | NS | 0.001030 | * |
| **PGA** | **SS-awn** | 0.898603 | NS | 0.794030 | NS | 0.972922 | NS | 0.998133 | NS |
| | **PS-awn** | 4.64E-09 | *** | 1.42E-08 | *** | 8.03E-09 | *** | 3.51E-06 | *** |
| | **PS-ss** | 2.51E-09 | *** | 5.46E-09 | *** | 5.82E-09 | *** | 3.15E-06 | *** |
| | **60-30** | 0.099404 | NS | 0.019608 | . | 0.295943 | NS | 0.956656 | NS |
| | **300-30** | 1.60E-06 | *** | 0.008241 | * | 1.40E-08 | *** | 5.41E-07 | *** |
| | **300-60** | 0.000122 | ** | 0.913535 | NS | 1.59E-07 | *** | 8.95E-07 | *** |

| | | | | | | | | | |
|---|---|---|---|---|---|---|---|---|---|
| **PYR** | **SS-awn** | 0.224349 | NS | 0.643493 | NS | 0.417001 | NS | 0.452497 | NS |
| | **PS-awn** | 0.063394 | NS | 0.088859 | NS | 0.999407 | NS | 0.009128 | * |
| | **PS-ss** | 0.001645 | * | 0.014250 | . | 0.399610 | NS | 0.000625 | ** |
| | **60-30** | 0.813809 | NS | 0.192965 | NS | 0.484787 | NS | 0.452799 | NS |
| | **300-30** | 3.87E-05 | *** | 0.000211 | ** | 0.420890 | NS | 6.68E-05 | *** |
| | **300-60** | 0.000140 | ** | 0.010950 | . | 0.060303 | NS | 5.70E-06 | *** |
| **TP** | **SS-awn** | 0.223132 | NS | 0.084916 | NS | 0.997963 | NS | 0.668999 | NS |
| | **PS-awn** | 1.33E-09 | *** | 6.40E-08 | *** | 2.55E-08 | *** | 1.81E-10 | *** |
| | **PS-ss** | 1.53E-08 | *** | 3.38E-06 | *** | 2.80E-08 | *** | 5.25E-10 | *** |
| | **60-30** | 0.020536 | . | 0.009985 | * | 0.111734 | NS | 0.769233 | NS |
| | **300-30** | 1.18E-07 | *** | 0.013680 | . | 8.77E-10 | *** | 1.46E-11 | *** |
| | **300-60** | 2.85E-05 | *** | 0.988165 | NS | 1.74E-08 | *** | 3.14E-11 | *** |
| **UDPG** | **SS-awn** | 0.024848 | . | 0.001441 | * | 0.282467 | NS | 0.940750 | NS |
| | **PS-awn** | 2.41E-06 | *** | 0.703704 | NS | 1.11E-05 | *** | 1.50E-12 | *** |
| | **PS-ss** | 1.85E-08 | *** | 0.000250 | ** | 5.88E-07 | *** | 2.08E-12 | *** |
| | **60-30** | 0.910616 | NS | 0.945604 | NS | 0.882585 | NS | 0.787569 | NS |
| | **300-30** | 2.20E-07 | *** | 0.090695 | NS | 4.88E-06 | *** | 4.84E-13 | *** |
| | **300-60** | 4.47E-07 | *** | 0.160648 | NS | 1.24E-05 | *** | 8.48E-13 | *** |

C. Organ x time interactions. Significant values in boldface. Only rows containing at least one significant value are included.

| | | **M0** | **M1** | **M2** | **M3** |
|---|---|---|---|---|---|
| **ADPG** | **PS:60-A:30** | 0.441161 | **0.039234** | 1.000000 | 0.861362 |
| | **PS:300-A:30** | **6.18E-07** | 0.164329 | 0.267783 | **3.59E-08** |
| | **PS:300-SS:30** | **1.16E-07** | 0.253869 | **0.000318** | **1.94E-08** |
| | **PS:300-PS:30** | **6.92E-07** | 0.933281 | **0.002845** | **3.21E-08** |
| | **PS:300-A:60** | **6.10E-07** | 0.918981 | **0.002423** | **3.33E-08** |
| | **PS:300-SS:60** | **1.29E-07** | 0.239942 | **0.000428** | **2.39E-08** |
| | **PS:300-PS:60** | **2.72E-05** | 0.996545 | 0.195878 | **3.02E-07** |
| | **PS:300-A:300** | **2.11E-07** | 0.234591 | **0.000441** | **7.61E-08** |
| | **PS:300-SS:300** | **1.40E-06** | 0.866466 | **0.003530** | **2.34E-07** |
| **ASP** | **PS:60-A:30** | 0.171237086 | **0.022209792** | 0.983372821 | 0.999745568 |
| | **PS:300-A:30** | 0.052063454 | 0.102076827 | **0.028349988** | **0.008829978** |
| | **PS:60-SS:30** | 0.175254627 | **0.024943775** | 0.971709123 | 0.999786564 |
| | **PS:300-SS:30** | 0.05346476 | 0.113327433 | **0.023570048** | **0.009060794** |
| | **PS:300-PS:30** | 0.489738351 | 0.95980419 | **0.0429197** | **0.011968658** |
| | **PS:60-A:60** | 0.210713351 | **0.032482614** | 0.981939679 | 0.999924173 |
| | **PS:300-A:60** | 0.066188206 | 0.143249098 | **0.027595171** | **0.010406728** |
| | **PS:60-SS:60** | 0.203870651 | **0.031807785** | 0.978173912 | 0.999793885 |
| | **PS:300-SS:60** | 0.063682748 | 0.140632355 | **0.025881075** | **0.009106588** |
| | **A:300-PS:60** | 0.19876422 | **0.026332848** | 0.989751843 | 0.999977562 |
| | **PS:300-PS:60** | 0.998936037 | 0.996038 | 0.178611043 | **0.026185279** |
| | **PS:300-A:300** | 0.061828745 | 0.118958581 | **0.032837167** | **0.011943248** |
| | **PS:300-SS:300** | 0.128283593 | 0.233426644 | 0.069066259 | **0.030154933** |
| **F6P** | **PS:30-A:30** | 0.141283747 | **0.000844674** | 0.64907191 | 0.943644624 |
| | **PS:60-A:30** | **0.000782674** | **4.45E-06** | **0.004965677** | **0.025739843** |
| | **PS:300-A:30** | **1.47E-11** | **0.000106454** | **4.89E-09** | **4.73E-14** |
| | **PS:30-SS:30** | **0.030902112** | **0.000226142** | 0.111658694 | 0.86893877 |

| | | | | | |
|---|---|---|---|---|---|
| | **PS:60-SS:30** | **0.000158485** | **1.47E-06** | **0.000404246** | **0.016243059** |
| | **PS:300-SS:30** | **6.89E-12** | **3.07E-05** | **1.12E-09** | **4.40E-14** |
| | **A:60-PS:30** | 0.182142925 | **0.000849721** | 0.822644376 | 0.984323879 |
| | **SS:60-PS:30** | 0.063234785 | **0.000706769** | 0.200629621 | 0.919433785 |
| | **A:300-PS:30** | 0.177968809 | **0.000484719** | 0.682011809 | 0.999592606 |
| | **PS:300-PS:30** | **3.34E-10** | 0.975804883 | **5.88E-08** | **7.99E-14** |
| | **PS:60-A:60** | **0.001057421** | **4.47E-06** | **0.009522657** | **0.041760222** |
| | **PS:300-A:60** | **1.70E-11** | **0.000107058** | **7.29E-09** | **5.17E-14** |
| | **PS:60-SS:60** | **0.000325472** | **3.83E-06** | **0.000798832** | **0.021586217** |
| | **PS:300-SS:60** | **9.63E-12** | **8.99E-05** | **1.67E-09** | **4.60E-14** |
| | **A:300-PS:60** | **0.001028135** | **2.79E-06** | **0.005570392** | 0.09059362 |
| | **SS:300-PS:60** | 0.054735529 | **0.00051261** | 0.075031099 | 0.940062209 |
| | **PS:300-PS:60** | **7.04E-09** | 0.719604882 | **4.64E-06** | **3.66E-13** |
| | **PS:300-A:300** | **1.67E-11** | **6.29E-05** | **5.25E-09** | **6.13E-14** |
| | **PS:300-SS:300** | **1.29E-10** | **0.018191505** | **2.87E-08** | **1.73E-13** |
| **G6P** | **PS:60-A:30** | **0.008529384** | **0.001060759** | 0.061130035 | 0.364421014 |
| | **PS:300-A:30** | **2.68E-13** | **1.09E-05** | **2.92E-13** | **2.53E-14** |
| | **PS:60-SS:30** | **0.005243047** | **0.003237914** | **0.001177764** | 0.24585675 |
| | **SS:300-SS:30** | 0.21558182 | 0.677338246 | **0.045074082** | 0.096209491 |
| | **PS:300-SS:30** | **2.32E-13** | **2.93E-05** | **9.46E-14** | **2.53E-14** |
| | **PS:60-PS:30** | 0.089728977 | 0.109064291 | **0.00776564** | 0.544249995 |
| | **PS:300-PS:30** | **5.62E-13** | **0.00089758** | **1.57E-13** | 2.53E-01 |
| | **PS:60-A:60** | **0.014401074** | **0.002427771** | 0.083501512 | 0.405300766 |
| | **PS:300-A:60** | **3.14E-13** | **2.27E-05** | **3.23E-13** | **2.53E-14** |
| | **PS:60-SS:60** | **0.005797829** | **0.002520069** | **0.003396761** | 0.264203432 |
| | **PS:300-SS:60** | **2.39E-13** | **2.34E-05** | **1.25E-13** | **2.53E-14** |
| | **A:300-PS:60** | 0.064544645 | **0.010641945** | 0.125242977 | 0.87363309 |
| | **PS:300-PS:60** | **5.80E-12** | 0.35207114 | **2.89E-12** | **2.53E-14** |
| | **PS:300-A:300** | **5.03E-13** | **8.67E-05** | **3.71E-13** | **2.53E-14** |
| | **PS:300-SS:300** | **1.32E-12** | **0.001005251** | **8.19E-13** | **2.53E-14** |
| **MAL** | **PS:30-A:30** | 0.098519772 | **0.016679807** | 0.997562061 | 0.998824946 |
| | **PS:60-A:30** | **0.001025173** | **9.45E-05** | 0.426044288 | 0.665083908 |
| | **PS:300-A:30** | **2.18E-05** | **0.00086632** | **6.41E-07** | **1.80E-10** |
| | **PS:30-SS:30** | 0.143172763 | **0.030596876** | 0.997909518 | 0.988528661 |
| | **PS:60-SS:30** | **0.001569288** | **0.000169062** | 0.434911208 | 0.505975525 |
| | **PS:300-SS:30** | **3.19E-05** | **0.00159915** | **6.57E-07** | **1.35E-10** |
| | **SS:60-PS:30** | 0.163704084 | **0.035465078** | 0.999202996 | 0.992789837 |
| | **A:300-PS:30** | 0.163201676 | **0.026992573** | 0.999987008 | 0.999998803 |
| | **PS:300-PS:30** | **0.011457905** | 0.870375502 | **2.10E-06** | **3.71E-10** |
| | **PS:60-A:60** | **0.004528083** | **0.000591167** | 0.553709791 | 0.621519699 |
| | **PS:300-A:60** | **8.41E-05** | **0.005866219** | **9.12E-07** | **1.67E-10** |
| | **PS:60-SS:60** | **0.001839646** | **0.000195464** | 0.4867054 | 0.544324655 |
| | **PS:300-SS:60** | **3.69E-05** | **0.001861959** | **7.60E-07** | **1.45E-10** |
| | **A:300-PS:60** | **0.001832872** | **0.000149646** | 0.643969847 | 0.869509621 |
| | **SS:300-PS:60** | **0.012986607** | **0.000783369** | 0.995991663 | 0.999999997 |
| | **PS:300-PS:60** | 0.574461354 | 0.964044076 | **2.98E-05** | **1.40E-09** |
| | **PS:300-A:300** | **3.67E-05** | **0.001406783** | **1.16E-06** | **2.77E-10** |
| | **PS:300-SS:300** | **0.000226799** | **0.007820261** | **7.24E-06** | **1.62E-09** |
| **P5P** | **PS:30-A:30** | **0.020025139** | **0.001772771** | 0.967549848 | 0.997852157 |
| | **PS:60-A:30** | **0.000465535** | **1.12E-05** | 0.10084684 | 0.826693205 |
| | **SS:300-A:30** | 0.224303071 | **0.029920567** | 0.439015726 | 0.997645606 |
| | **PS:300-A:30** | **1.08E-05** | 0.668812796 | **0.045977898** | **4.27E-06** |
| | **PS:60-SS:30** | **0.001711727** | **0.000301525** | 0.132684682 | 0.765907504 |

| | | | | | |
|---|---|---|---|---|---|
| | **PS:300-SS:30** | **3.48E-05** | 1 | 0.061770352 | **3.48E-06** |
| | **A:60-PS:30** | **0.013371887** | **0.000935053** | 0.466397015 | 0.995766414 |
| | **SS:60-PS:30** | 0.133755858 | **0.035854869** | 0.999999991 | 0.993919334 |
| | **A:300-PS:30** | **0.014985111** | **0.002857634** | 0.740254165 | 0.993919334 |
| | **PS:300-PS:30** | **0.027501933** | 0.072182644 | 0.315875072 | **1.51E-05** |
| | **PS:60-A:60** | **0.000313558** | **6.47E-06** | **0.012256329** | 0.789386376 |
| | **SS:300-A:60** | 0.161395791 | **0.015887921** | 0.077368845 | 0.995411388 |
| | **PS:300-A:60** | **7.64E-06** | 0.482527707 | **0.005226741** | **3.76E-06** |
| | **PS:60-SS:60** | **0.00348679** | **0.000174986** | 0.455054362 | 0.765907504 |
| | **PS:300-SS:60** | **6.66E-05** | 0.999986771 | 0.254545518 | **3.48E-06** |
| | **A:300-PS:60** | **0.000350269** | **1.70E-05** | **0.031475183** | 0.765907504 |
| | **SS:300-PS:60** | 0.104028818 | **0.019136752** | 0.988298712 | 0.995105143 |
| | **PS:300-PS:60** | 0.576866157 | **0.000356356** | 0.999961978 | **6.89E-05** |
| | **SS:300-A:300** | 0.177424706 | **0.047342776** | 0.17802988 | 0.993447496 |
| | **PS:300-A:300** | **8.42E-06** | 0.797836582 | **0.013624647** | **3.48E-06** |
| | **PS:300-SS:300** | **0.001936412** | 0.597409945 | 0.906965169 | **1.54E-05** |
| **PGA** | **PS:30-A:30** | 0.295655093 | **0.014589812** | 0.998337838 | 0.999999999 |
| | **PS:60-A:30** | **0.000942009** | **1.19E-05** | 0.057944735 | 0.998977885 |
| | **PS:300-A:30** | **3.92E-09** | **7.48E-06** | **1.18E-10** | **1.06E-08** |
| | **PS:30-SS:30** | 0.194187991 | **0.006297992** | 0.999338487 | 0.999996287 |
| | **PS:60-SS:30** | **0.000549157** | **5.81E-06** | 0.068098959 | 0.99330974 |
| | **PS:300-SS:30** | **2.83E-09** | **3.69E-06** | **1.29E-10** | **8.10E-09** |
| | **SS:60-PS:30** | 0.141686588 | **0.006519668** | 0.937298673 | 0.999991857 |
| | **PS:60-PS:30** | 0.14117828 | **0.041686985** | 0.201654877 | 0.999702391 |
| | **A:300-PS:30** | 0.641596548 | **0.021542701** | 0.999993988 | 0.999345569 |
| | **PS:300-PS:30** | **1.15E-07** | **0.024646849** | **2.51E-10** | **1.21E-08** |
| | **PS:60-A:60** | **0.00800325** | **0.000244726** | 0.162641605 | 0.998756017 |
| | **PS:300-A:60** | **1.49E-08** | **0.000145987** | **2.18E-10** | **1.03E-08** |
| | **PS:60-SS:60** | **0.000379191** | **5.98E-06** | **0.019494343** | 0.991469514 |
| | **PS:300-SS:60** | **2.27E-09** | **3.80E-06** | **6.56E-11** | **7.77E-09** |
| | **A:300-PS:60** | **0.003435941** | **1.68E-05** | 0.332996754 | 1 |
| | **SS:300-PS:60** | **0.026205187** | **0.000185567** | 0.6951503 | 0.99997825 |
| | **PS:300-PS:60** | **1.41E-05** | 0.999998833 | **6.13E-09** | **2.50E-08** |
| | **PS:300-A:300** | **8.70E-09** | **1.05E-05** | **3.58E-10** | **2.72E-08** |
| | **PS:300-SS:300** | **3.27E-08** | **0.000111144** | **7.32E-10** | **4.25E-08** |
| **PYR** | **PS:300-A:30** | **0.000326052** | **0.001093038** | 0.998987923 | **4.30E-05** |
| | **PS:300-SS:30** | **1.18E-05** | **0.000861602** | 0.169448153 | **1.07E-06** |
| | **PS:300-PS:30** | **4.37E-05** | **0.004717893** | 0.189801449 | **9.11E-07** |
| | **PS:300-A:60** | **0.000156963** | **0.029784469** | 0.152995963 | **8.74E-07** |
| | **PS:300-SS:60** | **3.21E-05** | **0.003061138** | 0.197112817 | **8.74E-07** |
| | **PS:300-PS:60** | **0.000242706** | **0.031288027** | 0.238908114 | **1.33E-06** |
| | **PS:300-A:300** | **0.002219466** | 0.094249374 | 0.608683709 | **1.05E-05** |
| | **PS:300-SS:300** | **0.000967881** | **0.048609494** | 0.495926517 | **1.02E-05** |
| **TP** | **PS:30-A:30** | 0.14574671 | **0.01040567** | 1 | 0.999999662 |
| | **PS:60-A:30** | **0.000188643** | **1.40E-05** | 0.110161954 | 0.947620669 |
| | **SS:300-A:30** | **0.027166872** | 0.057108299 | 0.111649832 | 0.1389848 |
| | **PS:300-A:30** | **1.27E-09** | **8.31E-05** | **1.51E-10** | **2.68E-13** |
| | **PS:60-SS:30** | **0.000383714** | **8.47E-05** | **0.018699286** | 0.910276867 |
| | **SS:300-SS:30** | 0.0551806 | 0.30760206 | 0.018982903 | 0.109731087 |
| | **PS:300-SS:30** | **1.93E-09** | **0.000558792** | **5.66E-11** | **2.47E-13** |
| | **A:300-PS:30** | 0.686810553 | **0.039060228** | 0.812753574 | 0.990330363 |
| | **PS:300-PS:30** | **5.51E-08** | 0.346575496 | **1.60E-10** | **3.08E-13** |
| | **PS:60-A:60** | **0.001844512** | **0.00028964** | 0.131646931 | 0.976009521 |

| | | | | | |
|---|---|---|---|---|---|
| | **PS:300-A:60** | **4.96E-09** | **0.002019349** | **1.68E-10** | **2.95E-13** |
| | **PS:60-SS:60** | **0.001692545** | **0.000428223** | 0.05710207 | 0.934342046 |
| | **PS:300-SS:60** | **4.70E-09** | **0.003025835** | 1.03E-10 | 2.59E-13 |
| | **A:300-PS:60** | **0.001879299** | **4.65E-05** | 0.855225791 | 1 |
| | **SS:300-PS:60** | 0.323686726 | **0.012597411** | 1 | 0.703948158 |
| | **PS:300-PS:60** | **1.09E-05** | 0.346575496 | **5.28E-09** | **5.67E-13** |
| | **PS:300-A:300** | **5.01E-09** | **0.000296967** | **9.08E-10** | **5.41E-13** |
| | **PS:300-SS:300** | **2.00E-07** | 0.084246769 | **5.23E-09** | **1.86E-12** |
| **UDPG** | **SS:30-A:30** | 0.078649435 | **0.009570051** | 0.494032424 | 0.99970549 |
| | **SS:60-A:30** | 0.149302194 | **0.023839265** | 0.656053157 | 0.999879879 |
| | **PS:300-A:30** | **1.78E-08** | 0.997026502 | **2.50E-07** | **2.66E-14** |
| | **PS:60-SS:30** | 0.051866115 | **0.015271254** | 0.217372054 | 0.673095471 |
| | **PS:300-SS:30** | **3.43E-10** | **0.002025421** | **1.24E-08** | **2.60E-14** |
| | **PS:300-PS:30** | **2.50E-09** | 0.16210025 | **5.51E-08** | **2.66E-14** |
| | **PS:300-A:60** | **2.98E-09** | 0.279051976 | **5.34E-08** | **2.64E-14** |
| | **PS:60-SS:60** | 0.101200364 | **0.037592705** | 0.329338105 | 0.709191585 |
| | **PS:300-SS:60** | **5.08E-10** | **0.005094248** | **1.78E-08** | **2.62E-14** |
| | **PS:300-PS:60** | **2.39E-08** | 0.98351863 | **6.15E-07** | **2.89E-14** |
| | **PS:300-A:300** | **6.87E-09** | 0.586753636 | **9.30E-08** | **2.73E-14** |
| | **PS:300-SS:300** | **5.16E-09** | 0.168680856 | **1.21E-07** | **3.18E-14** |

Table S6. *Themeda triandra*, $^{13}C$ data, analysis of variance. Average labeling of each metabolite. ADPG, ADP-glucose; ASP, aspartate; F6P, fructose-6-phosphate; G6P, glucose-6=phosphate; MAL, malate; P5P, pentose-5 phosphates; PGA, phosphoglycerate; PYR, pyruvate; UDPG, UDP-glucose. Triose phosphate was investigated but was unlabeled. Significance codes: ≤0.0001, ***; ≤ 0.001,**; ≤ 0.01,*; ≤ 0.05, .; NS, non-significant.

A. Main effects.

| | ADPG | | ASP | | F6P | | G6P | | MAL | |
|---|---|---|---|---|---|---|---|---|---|---|
| **organ** | 0.114381 | NS | 1.26E-07 | *** | 5.73E-07 | *** | 6.58E-06 | *** | 5.67E-08 | *** |
| **time** | 0.553770 | NS | 0.000745 | ** | 4.42E-05 | *** | 0.085881 | NS | 0.001853 | * |
| **organ*time** | 0.184321 | NS | 0.000184 | ** | 6.24E-06 | *** | 0.086412 | NS | 0.000124 | ** |

| | P5P | | PGA | | PYR | | UDPG | |
|---|---|---|---|---|---|---|---|---|
| **organ** | 0.233849 | NS | 0.589927 | NS | 1.23E-06 | *** | 0.000291 | ** |
| **time** | 0.323862 | NS | 0.12515 | NS | 1.25E-06 | *** | 0.000234 | ** |
| **organ*time** | 0.069049 | NS | 0.77500 | NS | 1.67E-07 | *** | 3.32E-05 | *** |

B. Organ and time. Tukey's honestly significant difference.

| | ADPG | | ASP | | F6P | | G6P | | MAL | |
|---|---|---|---|---|---|---|---|---|---|---|
| **SS-awn** | 0.861379 | NS | 0.985358 | NS | 0.506551 | NS | 0.943214 | NS | 0.880378 | NS |
| **STS-awn** | 0.112786 | NS | 7.07E-07 | *** | 8.64E-06 | *** | 1.88E-05 | *** | 4.37E-07 | *** |
| **STS-ss** | 0.270265 | NS | 5.30E-07 | *** | 1.03E-06 | *** | 3.64E-05 | *** | 1.91E-07 | *** |
| **60-30** | 0.972899 | NS | 0.964799 | NS | 0.588625 | NS | 0.658311 | NS | 0.967942 | NS |
| **300-30** | 0.692297 | NS | 0.001518 | * | 6.17E-05 | *** | 0.073931 | NS | 0.003481 | * |
| **300-60** | 0.556585 | NS | 0.002655 | * | 0.000513 | ** | 0.328322 | NS | 0.005921 | * |

| | P5P | | PGA | | PYR | | UDPG | |
|---|---|---|---|---|---|---|---|---|
| **SS-awn** | 0.213692 | NS | 0.645987 | NS | 0.634804 | NS | 0.676894 | NS |
| **STS-awn** | 0.511467 | NS | 0.999998 | NS | 1.43E-05 | *** | 0.002465 | * |
| **STS-ss** | 0.807165 | NS | 0.644840 | NS | 2.43E-06 | *** | 0.000385 | ** |
| **60-30** | 0.573255 | NS | 0.272801 | NS | 0.908294 | NS | 0.898329 | NS |
| **300-30** | 0.305277 | NS | 0.126410 | NS | 8.22E-06 | *** | 0.000446 | ** |
| **300-60** | 0.872823 | NS | 0.888221 | NS | 3.67E-06 | *** | 0.001170 | ** |

C. Organ x time interactions. Significant values in boldface. Only rows containing at least one significant value are included.

| | ADPG | ASP | F6P | G6P | MAL |
|---|---|---|---|---|---|
| **STS:300-A:30** | 0.359848 | **8.42E-07** | **4.13E-07** | **0.000313** | **4.85E-07** |
| **STS:300-SS:30** | 0.477909 | **7.07E-07** | **1.74E-07** | **0.000604** | **9.46E-07** |
| **STS:300-STS:30** | 0.265064 | **5.40E-05** | **4.75E-07** | **0.031638** | **6.75E-05** |
| **STS:300-A:60** | 0.195544 | **1.35E-06** | **4.06E-07** | **0.000895** | **1.98E-06** |
| **STS:300-SS:60** | 0.215991 | **7.21E-07** | **1.54E-07** | **0.000616** | **3.80E-07** |
| **STS:300-STS:60** | 0.560465 | **7.02E-05** | **9.93E-06** | 0.201840 | **8.63E-05** |
| **STS:300-A:300** | 0.101175 | **8.70E-07** | **5.65E-07** | **0.000400** | **7.07E-07** |
| **STS:300-SS:300** | 0.291077 | **1.23E-06** | **1.76E-07** | **0.000936** | **4.90E-07** |

| | **P5P** | **PGA** | **PYR** | **UDPG** |
|---|---|---|---|---|
| **STS:300-A:30** | 0.673962 | 0.718591 | **1.95E-07** | **3.49E-05** |
| **STS:300-SS:30** | 0.494692 | 0.782377 | **9.92E-09** | **9.61E-06** |
| **STS:300-STS:30** | 0.290837 | 0.992998 | **2.78E-08** | **3.72E-06** |
| **STS:300-A:60** | 0.278966 | 1.000000 | **1.74E-08** | **9.11E-06** |
| **STS:300-SS:60** | 1.000000 | 0.986306 | **1.71E-08** | **9.84E-06** |
| **STS:300-STS:60** | 0.461005 | 0.997683 | **6.08E-08** | **5.48E-05** |
| **STS:300-A:300** | 0.401500 | 1.000000 | **2.88E-08** | **3.34E-05** |
| **STS:300-SS:300** | 0.832169 | 0.999556 | **6.16E-08** | **7.72E-06** |

Table S7. Spikelet and awn removal experiments, average seed weight (mg). Gray shading = pairs of values in which removal of the structure led to reduced seed weight. NA=not applicable; line is awnless.

A. Small-scale (pilot) experiment to test for presence/absence of an effect. Spikelet removal and awn removal both had significant effects, with effect size generally greater for PS removal. Considering all lines, PS removal only, p=0.0018 (paired T-test). *missing data replaced with mean for analysis. Effect of awn removal was not significant.

| Code | name | PS on awn on | PS off awns on | awn off PS on | awn off PS off |
|---|---|---|---|---|---|
| 534021 | Jola Nandyal | 28.2 | 26.4 | 27.6 | 26.8 |
| 534021 | Jola Nandyal | 25.2 | 24.3 | 22.7 | 20.5 |
| 534096 | SO-85 | 22.9 | 20 | -* | 23.2 |
| 534096 | SO-85 | 22.4 | 18.8 | 19.4 | 21.4 |
| 656014 | SAP-15 | 18 | 14.5 | 16 | 13.6 |
| 656014 | SAP-15 | 19.7 | 16.2 | 19.4 | 16.6 |
| 656099 | SAP-257 | 31 | 26 | 30.5 | 25 |
| 656099 | SAP-257 | 34.8 | 32.8 | 28.6 | 34.6 |
| 597971 | SAP-170 | 32 | 31 | NA | NA |
| 597971 | SAP-170 | 30 | 30 | NA | NA |
| 659691 | Combine hegari | 32.4 | 33.1 | NA | NA |
| 659691 | Combine hegari | 34.6 | 33.7 | NA | NA |
| | BTx623 | 29.9 | 29.6 | NA | NA |
| | BTx623 | 29 | 27.4 | NA | NA |
| | BTx623 | 22.6 | 21 | NA | NA |
| | BTx623 | 33.7 | 34.2 | NA | NA |
| | BTx623 | 36.5 | 34.7 | NA | NA |
| | BTx623 | 35.8 | 33.8 | NA | NA |
| **Mean, PS only, all lines** | | **28.82** | **27.08** | | |
| **Mean, awned lines only** | | **25.28** | **22.38** | **23.46** | **22.71** |

B. Experiment 1 to test the effect of PS removal and control for possible wounding response. p, p value for individual line, from anova. Overall effect of PS removal, p=0.108 (not significant). †Excluded from analyses because of missing data; if this line is included, overall p= 0.0255.

| | | | Unaltered plant | | Altered plant | | p |
|---|---|---|---|---|---|---|---|
| code | name | rep | control 1 | control 2 | control 3 | PS removed | |
| 534021 | Jola nandyal | 1 | 22.5817 | 21.2941 | 21.3779 | 19.0807 | |
| | | 2 | 13.3958 | 12.9116 | 12.5092 | 11.038 | |
| | | 3 | 27.8127 | 27.7527 | 27.6567 | 26.9528 | |
| | | **mean** | **21.2634** | **20.6528** | **20.5146** | **19.0238** | **0.0019** |

| | | | | | | | |
|---|---|---|---|---|---|---|---|
| 534096 | so85 | 1 | 20.0787 | 20.1045 | 18.3237 | 17.6512 | |
| | | 2 | 22.2868 | 23.25 | 17.4175 | 17.168 | |
| | | 3 | 18.6506 | 19.1616 | 17.4595 | 17.3756 | |
| | | 4 | 15.6197 | 16.3712 | 17.7041 | 17.2475 | |
| | | 5 | 20.2976 | 20.2669 | 16.5024 | 16.931 | |
| | | **mean** | **19.38668** | **19.83084** | **17.48144** | **17.2747** | **0.094** |
| 597971 | sap170 | 1 | 20.2761 | 20.4822 | 21.7447 | 20.5565 | |
| | | 2 | 19.915 | 19.7366 | 20.648 | 18.9695 | |
| | | 3 | 25.3559 | 24.773 | 26.7461 | 22.9521 | |
| | | 4 | 12.5674 | 18.2376 | 26.739 | 23.8691 | |
| | | 5 | 17.017 | 17.1057 | 26.154 | 25.5848 | |
| | | **mean** | **19.02628** | **20.06702** | **24.40636** | **22.3864** | **0.565** |
| 655995 | sap51 | 1 | 17.708 | 17.0794 | 17.8559 | 17.1795 | |
| | | 2 | 17.4825 | 17.4262 | 17.2538 | 16.5153 | |
| | | 3 | 18.2625 | 17.6768 | 17.7731 | 16.8158 | |
| | | **mean** | **17.8177** | **17.3941** | **17.6276** | **16.8369** | **0.0056** |
| 656014 | sap15 | 1 | 20.0872 | 18.9949 | 19.3521 | 20.1773 | |
| | | 2 | 19.7078 | 19.9798 | 15.0152 | 15.476 | |
| | | 3 | 19.6606 | 19.8416 | 18.5034 | 18.2385 | |
| | | 4 | 21.9626 | 21.0877 | 19.8094 | 19.3216 | |
| | | **mean** | **20.35455** | **19.976** | **18.170025** | **18.30335** | **0.19** |
| †656099 | sap257 | 1 | 32.9846 | 32.19 | 26.5788 | 18.0409 | |
| | | 2 | | | 32.2051 | | |
| | | 3 | | | 35.7525 | 30.6934 | 0.0527 |
| 659691 | Combin e hegari | 1 | 33.1778 | 33.2906 | 32.1471 | 30.475 | |
| | | 2 | 25.7581 | 25.7962 | 29.6308 | 29.6489 | |
| | | 3 | 34.2697 | 34.0315 | 29.1065 | 28.7598 | |
| | | **mean** | **31.0685** | **31.0394** | **30.2948** | **29.6279** | **0.4757** |
| btx623 | btx623 | 1 | 34.662 | 34.1934 | 35.336 | 31.6906 | |
| | | 2 | 33.2947 | 33.3116 | 33.7919 | 29.3817 | |
| | | 3 | 35.8384 | 35.4025 | 27.996 | 29.8358 | |
| | | **mean** | **34.5984** | **34.3025** | **32.375** | **30.3027** | **0.0381** |

C. Experiment 2 to test the effect of PS removal, awn removal, and PS+awn removal; only PS removal calculated. p, p value for PS removal only for individual line, from anova. Overall effect of PS removal only, p=0.09284 (not significant), awn removal, p=0.69173 (not significant).

| | | | plant A remove PS | | plant B remove awn | | plant C remove PS and awn | | plant D control | p |
|---|---|---|---|---|---|---|---|---|---|---|
| **Code** | **name** | **rep** | **on** | **off** | **on** | **off** | **on** | **off** | | |
| 534021 | Jola nan-dyal | 1 | 26.6818 | 26.0345 | 26.4960 | 26.8533 | 31.0407 | 30.1444 | 30.1854 | |
| | | 2 | 32.0812 | 28.7032 | 27.5349 | 27.6709 | 30.3660 | 27.9155 | 30.5350 | |
| | | 3 | 31.5640 | 31.2738 | 31.5762 | 31.4537 | 29.8343 | 28.9761 | 32.7598 | |
| | | 4 | 21.2500 | 18.7017 | 22.4053 | 22.0371 | 20.0420 | 18.0007 | 19.0618 | |
| | | 5 | 28.4169 | 27.8130 | 22.4072 | 21.6440 | 27.6145 | 27.5496 | 26.3633 | |
| | | 6 | 28.1821 | 27.0225 | 30.1023 | 29.5062 | 32.4243 | 32.1355 | 28.5988 | |
| | **mean** | | **28.0293** | **26.5914** | **26.7537** | **26.5275** | **28.5537** | **27.4536** | **27.9173** | **0.442** |
| 656099 | Sap-257 | 1 | 36.1653 | 36.6503 | 37.8258 | 37.0147 | 33.8095 | 33.4670 | 36.4390 | |
| | | 2 | 36.1873 | 36.1682 | 32.4015 | 31.2967 | 33.3481 | 32.0733 | 35.0965 | |
| | | 3 | 27.7984 | 25.5675 | 37.9149 | 37.8612 | 35.5478 | 33.1449 | 39.2459 | |
| | | 4 | 11.8756 | 9.1034 | 39.4475 | 40.5622 | 37.1174 | 35.8115 | 41.0308 | |
| | | 5 | 40.6955 | 40.5712 | 38.8632 | 37.2804 | 36.6557 | 34.9159 | 37.3090 | |
| | | 6 | 37.6536 | 38.6545 | 24.6485 | 24.1550 | 16.2006 | 16.3087 | 37.8178 | |
| | **mean** | | **31.7293** | **31.1192** | **35.1836** | **34.6950** | **32.1132** | **30.9535** | **37.8232** | **0.204** |
| 534096 | SO-85 | 1 | 15.8976 | 16.3371 | 15.6566 | 15.5149 | 14.2329 | 14.1980 | 14.2377 | |
| | | 2 | 12.8596 | 13.1284 | 10.7975 | 10.3481 | 13.3984 | 12.7493 | 16.0233 | |
| | | 3 | 16.3289 | 16.3794 | 14.2000 | 13.8630 | 14.3130 | 13.6380 | 19.3753 | |
| | | 4 | 11.8398 | 11.5310 | 14.6764 | 14.5612 | 10.3203 | 9.9143 | 11.9579 | |
| | | 5 | 12.8950 | 12.1761 | 19.6246 | 19.1962 | 14.4853 | 14.1580 | 12.8162 | |
| | | 6 | 14.9077 | 14.2925 | 15.4063 | 15.2545 | 13.0078 | 11.4700 | 15.9104 | |
| | **mean** | | **14.1214** | **13.9741** | **15.0602** | **14.7897** | **13.2929** | **12.6879** | **15.0535** | **0.098** |
| 656014 | SAP-15 | 1 | 17.3681 | 17.4735 | 18.4154 | 18.9236 | 17.1777 | 17.1861 | 17.4491 | |
| | | 2 | 17.5936 | 17.6506 | 16.6821 | 16.7063 | 19.4724 | 19.3528 | 20.1685 | |
| | | 3 | 17.9455 | 17.9249 | 18.6698 | 18.7610 | 18.7657 | 18.8634 | 17.5876 | |
| | | 4 | 19.4335 | 19.4512 | 18.9240 | 18.6876 | 18.4103 | 18.2291 | 19.6645 | |
| | | 5 | 18.9327 | 18.8410 | 19.2110 | 19.5825 | 19.0619 | 19.4200 | 19.7508 | |
| | | 6 | 17.8545 | 17.6757 | 17.4733 | 17.4819 | 16.3683 | 14.7568 | 17.6135 | |
| | **mean** | | **18.1880** | **18.1695** | **18.2293** | **18.3572** | **18.2094** | **17.9680** | **18.7057** | **0.365** |
| 655995 | SAP-51 | 1 | 19.1572 | 18.7448 | 18.8768 | 19.0326 | 20.3947 | 19.9671 | 19.2766 | |
| | | 2 | 18.4414 | 18.9396 | 18.0012 | 18.2891 | 18.4466 | 18.2522 | 19.0525 | |
| | | 3 | 16.4906 | 18.1418 | 16.9052 | 16.8452 | 20.0926 | 18.9000 | 17.9415 | |
| | **mean** | | **18.0297** | **18.6087** | **17.9277** | **18.0556** | **19.6446** | **19.0398** | **18.7569** | **0.422** |

Table S8. $^{14}C$ data for *Andropogon schirensis*, analysis of variance. Detached=spikelets, awns and bract removed from inflorescence. Attached=spikelets, awns, and bract on an intact inflorescence. Both detached and attached structures were given a one-hour pulse of $^{14}C$. Chase= spikelets, awns, and bracts on an intact inflorescence, exposed to $^{14}C$ for one hour, and then allowed to photosynthesize in air for another 24 hours. PS=pedicellate spikelet; SS=sessile spikelet; diff=difference between mean values; lwr=minus one standard deviation; upr=plus one standard deviation. Significance codes: ≤0.0001, ***; ≤ 0.001,**; ≤ 0.01,*; ≤ 0.05, . NS=not significant.

A. **Decays per minute per mg (dpm) and percent dpm**.

| | | **Rep1** | | **Rep2** | | **Rep3** | | **mean** |
|---|---|---|---|---|---|---|---|---|
| **treatment** | **organ** | **Dpm/mg** | **%** | **Dpm/mg** | **%** | **Dpm/mg** | **%** | **%** |
| **detached** | **Awn** | 37 | 0.14 | 34 | 0.24 | 16 | 0.42 | 0.3 |
| | **SS** | 3487 | 12.76 | 809 | 5.74 | 303 | 8.03 | 8.8 |
| | **PS** | 11958 | 43.77 | 3413 | 24.23 | 1209 | 32.02 | 33.3 |
| | **bract** | 11840 | 43.34 | 9828 | 69.78 | 2248 | 59.53 | 57.6 |
| **attached** | **Awn** | 40 | 0.10 | 10 | 0.03 | 23 | 0.12 | 0.1 |
| | **SS** | 7953 | 15.81 | 5621 | 13.45 | 4303 | 15.93 | 15.1 |
| | **PS** | 30587 | 60.8 | 25825 | 61.81 | 14300 | 52.94 | 58.5 |
| | **bract** | 11725 | 23.31 | 10322 | 24.71 | 8387 | 31.05 | 26.3 |
| **chase** | **Awn** | 1731 | 5.47 | 152 | 0.67 | 761 | 3.43 | 3.2 |
| | **SS** | 13859 | 43.78 | 5640 | 24.94 | 5391 | 24.28 | 31.0 |
| | **PS** | 9105 | 28.76 | 9846 | 43.54 | 6768 | 30.47 | 34.3 |
| | **bract** | 6960 | 21.99 | 6976 | 30.85 | 9287 | 41.82 | 31.5 |

B. **ANOVA Main effects**

| | **Pr(>F)** | |
|---|---|---|
| Organ | 1.66e-11 | *** |
| Treatment | 1 | NS |
| Organ * treatment | 3.45e-06 | *** |

C. **Comparisons of organs**

| | **diff** | **lwr** | **upr** | **p adj** | |
|---|---|---|---|---|---|
| **Bract-awn** | 37.31 | 27.95 | 46.68 | 0 | *** |
| **PS-awn** | 40.87 | 31.50 | 50.23 | 0 | *** |
| **SS-awn** | 17.13 | 7.76 | 26.49 | 0.00020 | ** |
| **PS-bract** | 3.55 | -5.81 | 12.92 | 0.72431 | NS |
| **SS-bract** | -20.19 | -29.55 | 10.82 | 0.00002 | *** |
| **SS-PS** | -23.74 | -33.10 | 14.37 | 0.00000 | *** |

**D. Organ x treatment interactions. Significant values ($p<0.05$) in boldface.**

| | diff | lwr | upr | p adj |
|---|---|---|---|---|
| Within organs, between treatments | | | | |
| ss:detached-ss:attached | -6.22 | -27.42 | 14.98 | 0.99397 |
| **ss:detached-ss:chase** | **-22.15** | **-43.35** | **-0.95** | **0.03508** |
| ss:chase-ss:attached | 15.94 | -5.26 | 37.14 | 0.27907 |
| **ps:detached-ps:attached** | **-25.18** | **-46.38** | **-3.98** | **0.01085** |
| ps:detached-ps:chase | -0.92 | -22.12 | 20.28 | 1.00000 |
| **ps:chase-ps:attached** | **-24.26** | **-45.46** | **-3.06** | **0.01559** |
| awn:detached-awn:attached | 0.21 | -20.99 | 21.41 | 1.00000 |
| awn:detached-awn:chase | -2.92 | -24.12 | 18.28 | 1.00000 |
| awn:chase-awn:attached | 3.13 | -18.07 | 24.33 | 0.99999 |
| **bract:detached-bract:attached** | **31.21** | **10.01** | **52.41** | **0.00093** |
| **bract:detached-bract:chase** | **26.01** | **4.81** | **47.21** | **0.00779** |
| bract:chase-bract:attached | 5.20 | -16.00 | 26.40 | 0.99868 |
| Within treatments, between organs | | | | |
| **bract:attached-awn:attached** | **26.29** | **5.09** | **47.49** | **0.00695** |
| **ps:attached-awn:attached** | **58.46** | **37.26** | **79.66** | **0.00000** |
| **ps:attached-bract:attached** | **32.17** | **10.97** | **53.37** | **0.00063** |
| **ss:attached-ps:attached** | **-43.46** | **-64.66** | **-22.26** | **0.00001** |
| ss:attached-awn:attached | 15.00 | -6.20 | 36.20 | 0.35728 |
| ss:attached-bract:attached | -11.29 | -32.49 | 9.91 | 0.73677 |
| **ps:detached-awn:detached** | **33.07** | **11.87** | **54.27** | **0.00043** |
| **ps:detached-bract:detached** | **-24.22** | **-45.42** | **-3.02** | **0.01586** |
| **ss:detached-bract:detached** | **-48.71** | **-69.91** | **-27.51** | **0.00000** |
| **ss:detached-ps:detached** | **-24.50** | **-45.70** | **-3.30** | **0.01421** |
| ss:detached-awn:detached | 8.58 | -12.62 | 29.78 | 0.93809 |
| **ss:detached-bract:detached** | **-48.71** | **-69.91** | **-27.51** | **0.00000** |
| **bract:chase-awn:chase** | **28.36** | **7.16** | **49.56** | **0.00300** |
| **ps:chase-awn:chase** | **31.07** | **9.87** | **52.27** | **0.00099** |
| ps:chase-bract:chase | 2.71 | -18.49 | 23.91 | 1.00000 |
| ss:chase-bract:chase | -0.55 | -21.75 | 20.65 | 1.00000 |
| ss:chase-ps:chase | -3.26 | -24.46 | 17.94 | 0.99998 |
| **ss:chase-awn:chase** | **27.81** | **6.61** | **49.01** | **0.00376** |
| Other comparisons | | | | |
| **awn:chase-bract:attached** | **-23.16** | **-44.36** | **-1.96** | **0.02388** |
| **awn:chase-ps:attached** | **-55.33** | **-76.53** | **-34.13** | **0.00000** |
| awn:chase-ss:attached | -11.87 | -33.07 | 9.33 | 0.67744 |
| **awn:detached-bract:attached** | **-26.08** | **-47.28** | **-4.88** | **0.00755** |
| **awn: detached-bract:chase** | **-31.28** | **-52.48** | **-10.08** | **0.00091** |
| **awn:detached-ps:attached** | **-58.25** | **-79.45** | **-37.05** | **0.00000** |
| **awn:detached-ps:chase** | **-33.99** | **-55.19** | **-12.79** | **0.00030** |
| awn:detached-ss:attached | -14.79 | -35.99 | 6.41 | 0.37612 |
| **awn:detached-ss:chase** | **-30.73** | **-51.93** | **-9.53** | **0.00114** |
| **bract:chase-awn:attached** | **31.49** | **10.29** | **52.69** | **0.00083** |
| **bract:chase-ps:attached** | **-26.97** | **-48.17** | **-5.77** | **0.00529** |

| | | | | |
|---|---|---|---|---|
| bract:chase-ss:attached | 16.49 | -4.71 | 37.69 | 0.23873 |
| **bract:detached-awn:attached** | **57.50** | **36.30** | **78.70** | **0.00000** |
| **bract:detached-awn:chase** | **54.37** | **33.17** | **75.57** | **0.00000** |
| bract:detached-ps:attached | -0.96 | -22.16 | 20.24 | 1.00000 |
| **bract:detached-ps:chase** | **23.30** | **2.10** | **44.50** | **0.02266** |
| **bract:detached-ss:attached** | **42.50** | **21.30** | **63.70** | **0.00001** |
| **bract:detached-ss:chase** | **26.56** | **5.36** | **47.76** | **0.00624** |
| **ps:chase-awn:attached** | **34.20** | **13.00** | **55.40** | **0.00027** |
| ps:chase-bract:attached | 7.91 | -13.29 | 29.11 | 0.96358 |
| ps:chase-ss:attached | 19.20 | -2.00 | 40.40 | 0.10154 |
| **ps:detached-awn:chase** | **30.15** | **8.95** | **51.35** | **0.00144** |
| **ps:detached-awn:attached** | **33.28** | **12.08** | **54.48** | **0.00040** |
| ps:detached-bract:attached | 6.99 | -14.21 | 28.19 | 0.98492 |
| ps:detached-bract:chase | 1.79 | -19.41 | 22.99 | 1.00000 |
| ps:detached-ss:attached | 18.28 | -2.92 | 39.48 | 0.13772 |
| ps:detached-ss:chase | 2.34 | -18.86 | 23.54 | 1.00000 |
| **ss:chase-awn:attached** | **30.94** | **9.74** | **52.14** | **0.00104** |
| ss:chase-bract:attached | 4.65 | -16.55 | 25.85 | 0.99952 |
| ss:chase-bract:chase | -0.55 | -21.75 | 20.65 | 1.00000 |
| **ss:chase-ps:attached** | **-27.52** | **-48.72** | **-6.32** | **0.00423** |
| **ss:detached-awn:attached** | 8.78 | -12.42 | 29.98 | 0.92829 |
| **ss:detached-awn:chase** | 5.66 | -15.54 | 26.86 | 0.99726 |
| ss:det-bract:attached | -17.51 | -38.71 | 3.69 | 0.17605 |
| **ss:det-bract:chase** | **-22.71** | **-43.91** | **-1.51** | **0.02845** |
| **ss:detached-ps:attached** | **-49.67** | **-70.87** | **-28.47** | **0.00000** |
| **ss:detached-ps:chase** | **-25.41** | **-46.61** | **-4.21** | **0.00988** |

Table S9. $^{14}C$ data for *Themeda triandra* with bract, analysis of variance. Raw data for Awn, SS, and PS the same as in Table S3. Detached=spikelets, awns and bract removed from inflorescence. Attached=spikelets, awns, and bract on an intact inflorescence. Both detached and attached structures were given a one-hour pulse of $^{14}C$. Chase= spikelets, awns, and bracts on an intact inflorescence, exposed to $^{14}C$ for one hour, and then allowed to photosynthesize in air for another 24 hours. PS=pedicellate spikelet; SS=sessile spikelet; diff=difference between mean values; lwr=minus one standard deviation; upr=plus one standard deviation. Significance codes: ≤0.0001, ***; ≤ 0.001,**; ≤ 0.01,*; ≤ 0.05, . NS=not significant.

A. **Decays per minute per mg (dpm/mg) and percent dpm/mg**.

| | | **Rep1** | | **Rep2** | | **Rep3** | | **mean** |
|---|---|---|---|---|---|---|---|---|
| **treatment** | **organ** | **Dpm/mg** | **%** | **Dpm/mg** | **%** | **Dpm/mg** | **%** | **%** |
| **detached** | **Awn** | 22 | 0.19 | 12 | 0.37 | 14 | 0.17 | 0.0 |
| | **SS** | 32 | 0.28 | 29 | 0.89 | 43 | 0.51 | 0.7 |
| | **PS** | 2870 | 24.92 | 788 | 24.18 | 3079 | 36.73 | 28.7 |
| | **bract** | 8592 | 74.61 | 2430 | 74.56 | 5246 | 62.59 | 71.0 |
| **attached** | **Awn** | 69 | 0.15 | 35 | 0.09 | 225 | 0.85 | 0.3 |
| | **SS** | 1761 | 3.83 | 1006 | 2.67 | 2139 | 8.08 | 5.0 |
| | **PS** | 14560 | 31.63 | 8860 | 23.54 | 9532 | 36.00 | 30.7 |
| | **bract** | 29644 | 64.40 | 27735 | 73.69 | 14585 | 55.08 | 64.3 |
| **chase** | **Awn** | 3589 | 11.96 | 3476 | 12.23 | 1437 | 8.16 | 10.7 |
| | **SS** | 8749 | 29.15 | 9823 | 34.56 | 9023 | 51.21 | 38.3 |
| | **PS** | 11284 | 37.60 | 5175 | 18.21 | 3746 | 21.26 | 25.7 |
| | **bract** | 6388 | 21.29 | 9950 | 35.01 | 3414 | 19.38 | 25.0 |

**B. ANOVA Main effects**

| | **Pr(>F)** | |
|---|---|---|
| Organ | 4.12e-13 | *** |
| Treatment | 0.998 | NS |
| Organ * treatment | 7.09e-09 | *** |

**C. Comparisons of organs**

| | **diff** | **lwr** | **upr** | **p adj** |
|---|---|---|---|---|
| **Bract-awn** | 49.78 | 40.87 | 58.68 | 0.00000 |
| **PS-awn** | 49.78 | 40.87 | 58.68 | 0.00000 |
| **SS-awn** | 11.00 | 2.10 | 19.90 | 0.01157 |
| **PS-bract** | -25.11 | -34.02 | -16.21 | 0.00000 |
| **SS-bract** | -38.78 | -47.68 | -29.87 | 0.00000 |
| **SS-PS** | -13.67 | -22.57 | -4.76 | 0.00155 |

**D. Organ x treatment interactions. Significant values (p<0.05) in boldface.**

| | diff | lwr | upr | p adj |
|---|---|---|---|---|
| Within organs, between treatments | | | | |
| ss:detached-ss:attached | -4.33 | -24.49 | 15.83 | 0.99960 |
| **ss:detached-ss:chase** | **-37.67** | **-57.83** | **-17.51** | **0.00003** |
| **ss:chase-ss:attached** | **33.33** | **13.17** | **53.49** | **0.00019** |
| ps:detached-ps:attached | -2.00 | -22.16 | 18.16 | 1.00000 |
| ps:detached-ps:chase | 3.00 | -17.16 | 23.16 | 0.99999 |
| ps:chase-ps:attached | -5.00 | -25.16 | 15.16 | 0.99855 |
| awn:detached-awn:attached | -0.33 | -20.49 | 19.83 | 1.00000 |
| awn:detached-awn:chase | -10.67 | -30.83 | 9.49 | 0.74394 |
| awn:chase-awn:attached | 10.33 | -9.83 | 30.49 | 0.77735 |
| bract:detached-bract:attached | 6.67 | -13.49 | 26.83 | 0.98459 |
| **bract:detached-bract:chase** | **46.00** | **25.84** | **66.16** | **0.00000** |
| **bract:chase-bract:attached** | **-39.33** | **-59.49** | **-19.17** | **0.00002** |
| **Within treatments, between organs** | | | | |
| **bract:attached-awn:attached** | **64.00** | **43.84** | **84.16** | **0.00000** |
| **ps:attached-awn:attached** | **30.33** | **10.17** | **50.49** | **0.00070** |
| **ps:attached-bract:attached** | **-33.67** | **-53.83** | **-13.51** | **0.00017** |
| **ss:attached-ps:attached** | **-25.67** | **-45.83** | **-5.51** | **0.00524** |
| ss:attached-awn:attached | 4.67 | -15.49 | 24.83 | 0.99922 |
| **ss:attached-bract:attached** | **-59.33** | **-79.49** | **-39.17** | **0.00000** |
| **ps:detached-awn:detached** | **28.67** | **8.51** | **48.83** | **0.00144** |
| **ps:detached-bract:detached** | **-42.33** | **-62.49** | **-22.17** | **0.00000** |
| **ss:detached-bract:detached** | **-70.33** | **-90.49** | **-50.17** | **0.00000** |
| **ss:detached-ps:detached** | **-28.00** | **-48.16** | **-7.84** | **0.00193** |
| ss:detached-awn:detached | 0.67 | -19.49 | 20.83 | 1.00000 |
| **ss:detached-bract:detached** | **-70.33** | **-90.49** | **-50.17** | **0.00000** |
| bract:chase-awn:chase | 14.33 | -5.83 | 34.49 | 0.35077 |
| ps:chase-awn:chase | 15.00 | -5.16 | 35.16 | 0.29178 |
| **ps:chase-bract:chase** | **-38.67** | **-58.83** | **-18.51** | **0.00002** |
| ss:chase-bract:chase | 13.33 | -6.83 | 33.49 | 0.45089 |
| ss:chase-ps:chase | 12.67 | -7.49 | 32.83 | 0.52345 |
| **ss:chase-awn:chase** | **27.67** | **7.51** | **47.83** | **0.00222** |
| **Other comparisons** | | | | |
| **awn:chase-bract:attached** | -53.67 | -73.83 | -33.51 | 0.00000 |
| **awn:chase-ps:attached** | -20.00 | -40.16 | 0.16 | 0.05315 |
| awn:chase-ss:attached | 5.67 | -14.49 | 25.83 | 0.99574 |
| **awn:detached-bract:attached** | -64.33 | -84.49 | -44.17 | 0.00000 |
| **awn: detached-bract:chase** | -25.00 | -45.16 | -4.84 | 0.00695 |
| **awn:detached-ps:attached** | -30.67 | -50.83 | -10.51 | 0.00061 |
| **awn:detached-ps:chase** | -25.67 | -45.83 | -5.51 | 0.00524 |
| awn:detached-ss:attached | -5.00 | -25.16 | 15.16 | 0.99855 |
| **awn:detached-ss:chase** | -38.33 | -58.49 | -18.17 | 0.00002 |
| **bract:chase-awn:attached** | 24.67 | 4.51 | 44.83 | 0.00800 |
| **bract:chase-ps:attached** | -5.67 | -25.83 | 14.49 | 0.99574 |

| | | | | |
|---|---|---|---|---|
| bract:chase-ss:attached | 20.00 | -0.16 | 40.16 | 0.05315 |
| **bract:detached-awn:attached** | 70.67 | 50.51 | 90.83 | 0.00000 |
| **bract:detached-awn:chase** | 60.33 | 40.17 | 80.49 | 0.00000 |
| bract:detached-ps:attached | 40.33 | 20.17 | 60.49 | 0.00001 |
| **bract:detached-ps:chase** | 45.33 | 25.17 | 65.49 | 0.00000 |
| **bract:detached-ss:attached** | 66.00 | 45.84 | 86.16 | 0.00000 |
| **bract:detached-ss:chase** | 32.67 | 12.51 | 52.83 | 0.00026 |
| **ps:chase-awn:attached** | 25.33 | 5.17 | 45.49 | 0.00604 |
| ps:chase-bract:attached | -38.67 | -58.83 | -18.51 | 0.00002 |
| ps:chase-ss:attached | 20.67 | 0.51 | 40.83 | 0.04106 |
| **ps:detached-awn:chase** | 18.00 | -2.16 | 38.16 | 0.11117 |
| **ps:detached-awn:attached** | 28.33 | 8.17 | 48.49 | 0.00167 |
| ps:detached-bract:attached | -35.67 | -55.83 | -15.51 | 0.00007 |
| ps:detached-bract:chase | 3.67 | -16.49 | 23.83 | 0.99992 |
| ps:detached-ss:attached | 23.67 | 3.51 | 43.83 | 0.01217 |
| ps:detached-ss:chase | -9.67 | -29.83 | 10.49 | 0.83814 |
| **ss:chase-awn:attached** | 38.00 | 17.84 | 58.16 | 0.00003 |
| ss:chase-bract:attached | -26.00 | -46.16 | -5.84 | 0.00454 |
| **ss:chase-ps:attached** | 7.67 | -12.49 | 27.83 | 0.95849 |
| **ss:detached-awn:attached** | 0.33 | -19.83 | 20.49 | 1.00000 |
| **ss:detached-awn:chase** | -10.00 | -30.16 | 10.16 | 0.80885 |
| ss:det-bract:attached | -63.67 | -83.83 | -43.51 | 0.00000 |
| **ss:det-bract:chase** | -24.33 | -44.49 | -4.17 | 0.00921 |
| **ss:detached-ps:attached** | -30.00 | -50.16 | -9.84 | 0.00081 |
| **ss:detached-ps:chase** | -25.00 | -45.16 | -4.84 | 0.00695 |
| ss:det-ss:attached | -4.33 | -24.49 | 15.83 | 0.99960 |

Table S10. *Themeda triandra*, $^{13}C$ data and analysis of variance, including data for the bract. PS includes proximal spikelet pairs, which are all staminate. Average labeling of each metabolite. ADPG, ADP-glucose; ASP, aspartate; F6P, fructose-6-phosphate; G6P, glucose-6=phosphate; MAL, malate; P5P, pentose-5 phosphates; PGA, phosphoglycerate; PYR, pyruvate; UDPG, UDP-glucose. Triose phosphate was investigated but was unlabeled. Significance codes: ≤0.0001, ***; ≤ 0.001,**; ≤ 0.01,*; ≤ 0.05, .; NS, non-significant.

A. Main effects.

| | **ADPG** | | **ASP** | | **F6P** | | **G6P** | | **MAL** | |
|---|---|---|---|---|---|---|---|---|---|---|
| **organ** | 3.75E-09 | *** | 2.94E-13 | *** | 6.56E-15 | *** | 2.16E-10 | *** | 3.85E-09 | *** |
| **time** | 0.000107 | ** | 5.32E-06 | *** | 5.53E-10 | *** | 1.46E-06 | *** | 2.82E-05 | *** |
| **organ*time** | 4.57E-06 | *** | 1.61E-05 | *** | 1.29E-10 | *** | 8.17E-08 | *** | 7.68E-05 | *** |

| | **P5P** | | **PGA** | | **PYR** | | **UDPG** | |
|---|---|---|---|---|---|---|---|---|
| **organ** | 0.258031 | NS | 0.628476 | NS | 4.91E-10 | *** | 6.27E-16 | *** |
| **time** | 0.421195 | NS | 0.597142 | NS | 6.86E-11 | *** | 5.45E-12 | *** |
| **organ*time** | 0.348409 | NS | 0.488737 | NS | 1.76E-10 | *** | 1.72E-12 | *** |

B. Organ and time. Tukey's honestly significant difference.

| | **ADPG** | | **ASP** | | **F6P** | | **G6P** | |
|---|---|---|---|---|---|---|---|---|
| **bract-awn** | 2.52E-08 | *** | 2.00E-12 | *** | 1.01E-13 | *** | 1.06E-09 | *** |
| **SS-awn** | 0.996459 | NS | 0.999866 | NS | 0.967069 | NS | 0.999174 | NS |
| **PS-awn** | 0.820852 | NS | 0.009630 | * | 0.056630 | NS | 0.082757 | NS |
| **SS-bract** | 3.95E-08 | *** | 1.80E-12 | *** | 6.52E-14 | *** | 1.36E-09 | *** |
| **PS-bract** | 1.62E-07 | *** | 6.96E-10 | *** | 3.73E-12 | *** | 1.82E-07 | *** |
| **PS-SS** | 0.912757 | NS | 0.008127 | * | 0.020591 | . | 0.107233 | NS |
| **60-30** | 0.610698 | NS | 0.484540 | NS | 0.012796 | . | 0.380783 | NS |
| **300-30** | 0.000140 | ** | 7.65E-06 | *** | 4.97E-10 | *** | 2.02E-06 | *** |
| **300-60** | 0.001549 | ** | 0.000139 | ** | 3.19E-07 | *** | 5.55E-05 | *** |

| | **P5P** | | **PGA** | | **PYR** | | **UDPG** | |
|---|---|---|---|---|---|---|---|---|
| **bract-awn** | 0.216520 | NS | 0.711681 | NS | 3.75E-09 | *** | 3.02E-14 | *** |
| **SS-awn** | 0.506962 | NS | 0.872748 | NS | 0.969784 | NS | 0.945756 | |
| **PS-awn** | 0.802975 | NS | 1.000000 | NS | 0.023463 | . | 0.075266 | NS |
| **SS-bract** | 0.933913 | NS | 0.989294 | NS | 1.58E-09 | *** | 2.55E-14 | *** |
| **PS-bract** | 0.696131 | NS | 0.710758 | NS | 3.09E-06 | *** | 2.70E-13 | *** |
| **PS-SS** | 0.957366 | NS | 0.872071 | NS | 0.008299 | * | 0.022932 | . |
| **60-30** | 0.803198 | NS | 0.896628 | NS | 0.934623 | NS | 0.009407 | * |
| **300-30** | 0.388384 | NS | 0.569490 | NS | 4.70E-10 | *** | 6.53E-12 | *** |
| **300-60** | 0.763000 | NS | 0.833222 | NS | 9.15E-10 | *** | 2.16E-09 | *** |

C. Organ x time interactions. Significant values in boldface. Only rows containing at least one significant value are included.

| | **ADPG** | **ASP** | **F6P** | **G6P** | **MAL** |
|---|---|---|---|---|---|

| | | | | | |
|---|---|---|---|---|---|
| **B:30-A:30** | 0.983379 | **0.006047** | 0.680573 | 0.904910 | 0.358168 |
| **B:60-A:30** | 0.099398 | **3.63E-05** | **5.02E-06** | **0.044496** | 0.096102 |
| **B:300-A:30** | **1.12E-08** | **5.52E-11** | **5.06E-14** | **9.40E-11** | **2.06E-08** |
| **PS:300-A:30** | 0.997192 | **0.019023** | **0.023725** | 0.438995 | **0.015921** |
| **SS:30-B:30** | 0.990768 | **0.005400** | 0.536440 | 0.944141 | 0.459767 |
| **A:60-B:30** | 0.959334 | **0.008168** | 0.677829 | 0.961162 | 0.578532 |
| **B:60-B:30** | 0.645410 | 0.603150 | 0.000562 | 0.631969 | 0.999733 |
| **SS:60-B:30** | 0.964043 | **0.005471** | 0.515726 | 0.945101 | 0.323927 |
| **A:300-B:30** | 0.920234 | **0.006175** | 0.728976 | 0.921141 | 0.414324 |
| **B:300-B:30** | **1.18E-07** | **1.92E-07** | **4.00E-13** | **1.30E-09** | **4.09E-06** |
| **SS:300-B:30** | 0.976314 | **0.007703** | 0.539206 | 0.962853 | 0.359843 |
| **PS:300-B:30** | 1.000000 | 0.999996 | 0.732084 | 0.999185 | 0.904611 |
| **B:60-SS:30** | 0.118575 | **3.25E-05** | **3.00E-06** | 0.059039 | 0.136759 |
| **B:300-SS:30** | **1.32E-08** | **5.12E-11** | **4.32E-14** | **1.16E-10** | **2.91E-08** |
| **PS:300-SS:30** | 0.998794 | **0.017055** | **0.013995** | 0.518592 | **0.024004** |
| **B:60-PS:30** | 0.083676 | **0.000427** | **5.44E-06** | 0.259814 | 0.643161 |
| **B:300-PS:30** | **9.54E-09** | **2.98E-10** | **5.21E-14** | **4.18E-10** | **2.33E-07** |
| **PS:300-PS:30** | 0.994320 | 0.168009 | **0.025749** | 0.926595 | 0.202553 |
| **B:60-A:60** | 0.071294 | **4.89E-05** | **4.97E-06** | 0.069508 | 0.195115 |
| **B:300-A:60** | **8.26E-09** | **6.76E-11** | **5.05E-14** | **1.32E-10** | **4.24E-08** |
| **PS:300-A:60** | 0.989972 | **0.025389** | **0.023486** | 0.567071 | **0.036878** |
| **SS:60-B:60** | 0.075055 | **3.29E-05** | **2.79E-06** | 0.059530 | 0.083944 |
| **PS:60-B:60** | 0.132455 | **0.000494** | **2.94E-05** | 0.469273 | 0.677094 |
| **A:300-B:60** | 0.051515 | **3.71E-05** | **6.01E-06** | **0.049493** | 0.117668 |
| **B:300-B:60** | **1.02E-05** | **2.07E-05** | **1.47E-09** | **6.47E-08** | 2.22E-05 |
| **SS:300-B:60** | 0.088079 | **4.61E-05** | **3.03E-06** | 0.070816 | 0.096716 |
| **PS:300-B:60** | 0.495370 | 0.317485 | 0.051692 | 0.977604 | 0.999297 |
| **B:300-SS:60** | **8.65E-09** | **5.17E-11** | **4.22E-14** | **1.17E-10** | **1.81E-08** |
| **SS:300-SS:60** | 1 | 1 | 1 | 1 | 1 |
| **PS:300-SS:60** | 0.991576 | **0.017269** | **0.012954** | 0.521012 | **0.013647** |
| **B:300-PS:60** | **1.48E-08** | **3.29E-10** | **1.00E-13** | **8.26E-10** | **2.61E-07** |
| **PS:300-PS:60** | 0.999344 | 0.187426 | 0.122709 | 0.991758 | 0.222919 |
| **B:300-A:300** | **6.21E-09** | **5.60E-11** | **5.38E-14** | **1.02E-10** | **2.50E-08** |
| **PS:300-A:300** | 0.974044 | **0.019410** | **0.028455** | 0.468192 | 0.020118 |
| **SS:300-B:300** | **1.00E-08** | **6.50E-11** | **4.33E-14** | **1.34E-10** | **2.07E-08** |
| **PS:300-B:300** | **7.27E-08** | **7.11E-08** | **4.89E-12** | **5.85E-09** | **0.000154** |
| **PS:300-SS:300** | 0.995344 | **0.024003** | **0.014139** | 0.572693 | **0.016038** |

| | P5P | PGA | PYR | UDPG |
|---|---|---|---|---|
| **B:30-A:30** | 0.999849 | 0.984646 | 1.000000 | 0.942024 |
| **B:60-A:30** | 0.999987 | 1.000000 | 0.999805 | **6.34E-06** |
| **B:300-A:30** | 0.976651 | 1.000000 | **4.66E-12** | **2.18E-14** |
| **PS:300-A:30** | 0.938701 | 0.938373 | **0.002220** | **0.005118** |
| **SS:30-B:30** | 0.997742 | 0.992051 | 0.999862 | 0.747770 |
| **A:60-B:30** | 0.974286 | 0.999998 | 0.999996 | 0.736715 |
| **B:60-B:30** | 1.000000 | 0.945829 | 0.994173 | **0.000183** |
| **SS:60-B:30** | 0.999525 | 0.999990 | 0.999995 | 0.752659 |
| **A:300-B:30** | 0.993157 | 1.000000 | 1.000000 | 0.938077 |
| **B:300-B:30** | 0.999983 | 0.997551 | **2.94E-12** | **2.50E-14** |
| **SS:300-B:30** | 0.999998 | 1.000000 | 1.000000 | 0.700823 |
| **PS:300-B:30** | 0.999683 | 1.000000 | **0.000992** | 0.125624 |
| **B:60-SS:30** | 0.999545 | 1.000000 | 0.829854 | **2.29E-06** |
| **B:300-SS:30** | 0.925761 | 1.000000 | **1.18E-12** | **2.15E-14** |
| **PS:300-SS:30** | 0.852816 | 0.960639 | **0.000187** | **0.001707** |
| **B:60-PS:30** | 0.991539 | 0.996833 | 0.958044 | **1.06E-06** |
| **B:300-PS:30** | 0.793106 | 0.999995 | **1.91E-12** | **2.14E-14** |
| **PS:300-PS:30** | 0.681097 | 0.999813 | **0.000458** | **0.000732** |
| **B:60-A:60** | 0.990109 | 0.758073 | 0.914033 | **2.19E-06** |
| **B:300-A:60** | 0.781257 | 0.951486 | **1.53E-12** | **2.15E-14** |
| **PS:300-A:60** | 0.667495 | 1.000000 | **0.000307** | **0.001630** |
| **SS:60-B:60** | 0.997666 | 0.998481 | 0.911664 | **2.33E-06** |
| **PS:60-B:60** | 0.999225 | 0.992449 | 0.991519 | **8.99E-06** |
| **A:300-B:60** | 0.998129 | 0.878116 | 0.960527 | **6.12E-06** |
| **B:300-B:60** | 0.999813 | 0.999997 | **1.31E-11** | **8.93E-12** |
| **SS:300-B:60** | 1.000000 | 0.982435 | 0.991798 | **1.92E-06** |
| **PS:300-B:60** | 0.998298 | 0.854091 | **0.012256** | 0.229248 |
| **B:300-SS:60** | 1.000000 | 0.999999 | **1.52E-12** | **2.15E-14** |
| **SS:300-SS:60** | 0.977530 | 1.000000 | 0.999998 | 1 |
| **PS:300-SS:60** | 1.000000 | 0.999495 | **0.000301** | **0.001743** |
| **B:300-PS:60** | 0.910857 | 0.999961 | **2.74E-12** | **2.20E-14** |
| **PS:300-PS:60** | 0.830964 | 0.999963 | **0.000878** | **0.007384** |
| **B:300-A:300** | 0.878805 | 0.987577 | **1.94E-12** | **2.18E-14** |
| **PS:300-A:300** | 0.786800 | 1.000000 | **0.000472** | **0.004936** |
| **SS:300-B:300** | 0.995260 | 0.999760 | **2.76E-12** | **2.15E-14** |
| **PS:300-B:300** | 1.000000 | 0.982238 | **1.66E-08** | **2.28E-13** |
| **PS:300-SS:300** | 0.981573 | 0.999996 | **0.000888** | **0.001408** |

Table S11. Numbers of unique transcripts after successive filters, starting with raw count numbers, for *Sorghum bicolor* and *Themeda triandra* and their associated figures. DE = differential expression

| **Set #** | **Transcript set** | ***Sorghum bicolor*** | **Used to produce Figure** | ***Themeda triandra*** | **Used to produce Figure** |
|---|---|---|---|---|---|
| 1 | Input | 47121 | | 202776 | |
| 2 | Expression filter pass | 17323 | | 20826 | |
| 3 | Metabolic genes: curated list | 3505 | | 1826 | |
| 4 | Metabolic genes: overlap with set 2 | 1441 | S8 | 769 | S10 |
| 5 | DE between any 2 organs in set 4 ($q < 0.05$) | 922 | S9 | 322 | S11 |
| 6 | Metabolic enzymes producing $^{13}C$ metabolites (may correspond to multiple genes) | 36 | | 36 | |
| 7 | Set 5 (DE between any 2 organs; $q < 0.05$) overlap with set 6 | 54 | 5 | 24 | S12 |

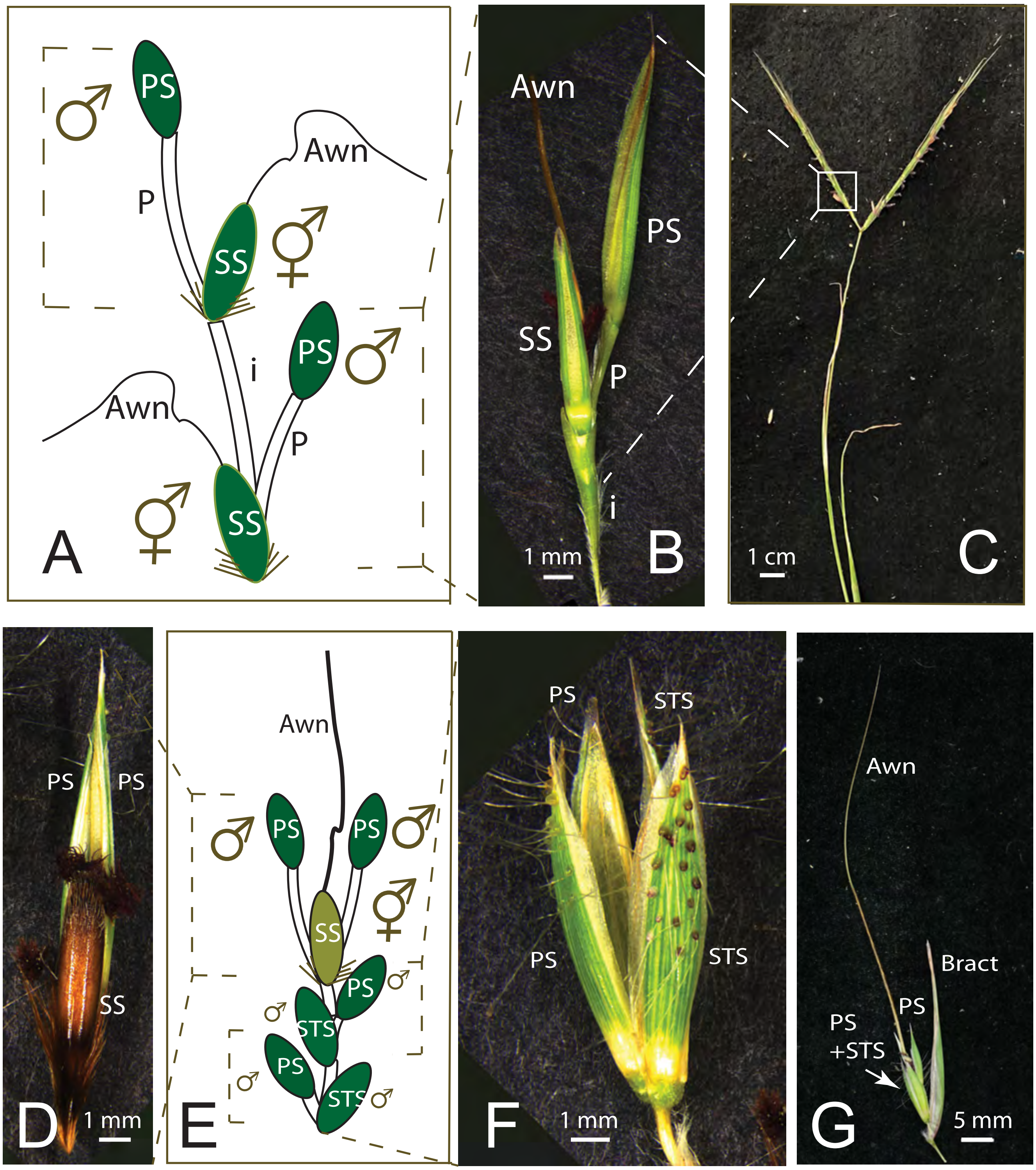

Figure S1. Spikelet pair and inflorescence structure in *Andropogon schirensis* (A, B, C), and *Themeda triandra* (D, E, F, G). A. Illustration of two spikelet pairs of A. schirensis, marked by dotted lines. Each pair is composed of a sessile spikelet, which includes a bisexual flower and bears the seed, and a pedicellate spikelet, which is staminate. The sessile spikelet bears a twisted awn from the lemma (floral bract). B. Spikelet pair of *A. schirensis.* C. Inflorescence of *A. schirensis*, showing two branches, each bearing 9-10 spikelet pairs. D. Spikelet pair of T. triandra, showing the dark indurate sessile spikelet, with two greenish pedicellate spikelets behind. One of the pedicellate spikelets is terminal on the short branch, so the three spikelets represent a pair plus a terminal spikelet that is morphologically identical to the pedicellate spikelet. E. Illustration of inflorescence structure in T. triandra, with three spikelet pairs, marked by dotted lines, and a terminal spikelet that is morphologically identical to the pedicellate spikelet. Spikelets in the proximal two pairs are all staminate; the distal pair includes a seed-bearing sessile spikelet and a staminate pedicellate spikelet. The sessile spikelet bears a twisted awn from the lemma (floral bract). F. Proximal spikelet pairs of T. triandra. All four proximal spikelets are staminate. G. Inflorescence branch of T. triandra, showing the spikelet complex as in D and E, subtended by a leaf-like bract. ss, sessile spikelet; ps, pedicellate spikelet; sts, staminate sessile spikelet. Scale bars B, D, F = 1 mm; C = 1 cm; G = 5 mm.

### Figure S2.

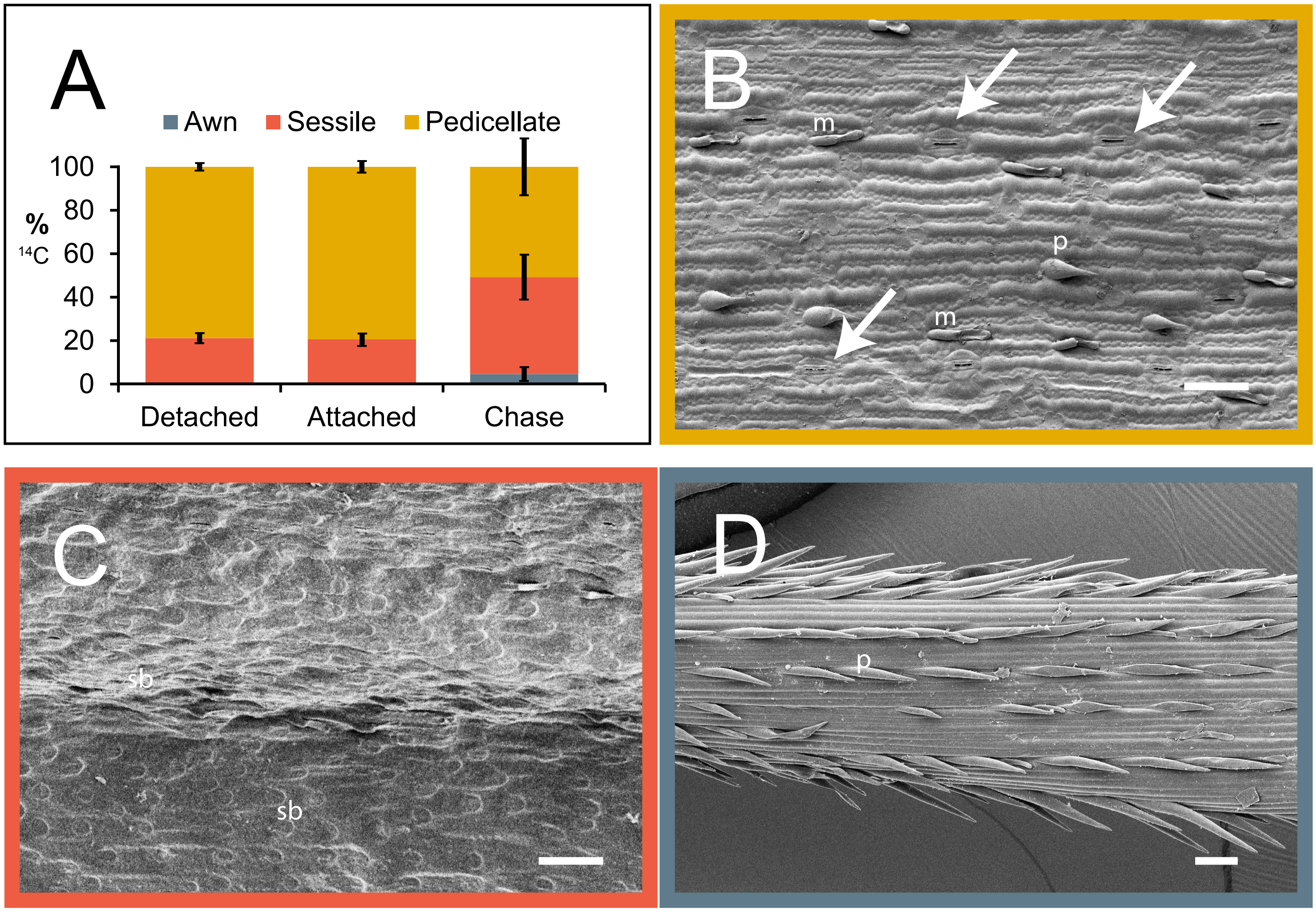

Figure S2. *Andropogon schirensis*. A. $^{14}C$ results. Percent dpm for each organ after 1 hour exposure to $^{14}C$ with organs removed from the axis (detached), inflorescence intact (attached), or after 24-hour chase. Plot includes mean percentages and standard deviations, (n=4). Values are similar after 1 hour, whether organs are attached or lying on filter paper. Change after 24-hour chase suggests movement of $^{14}C$ from PS to SS. Awn is largely unlabeled. B. Abaxial epidermis of pedicellate spikelet showing rows of stomata (arrows), bicellular microhairs (m), and prickles (p). C. Abaxial epidermis of sessile spikelet showing no stomata, but silica bodies (sb). D. Awn showing no stomata, but sparse macrohairs. Scale = 50 µm; note that B and C are more highly magnified than D.

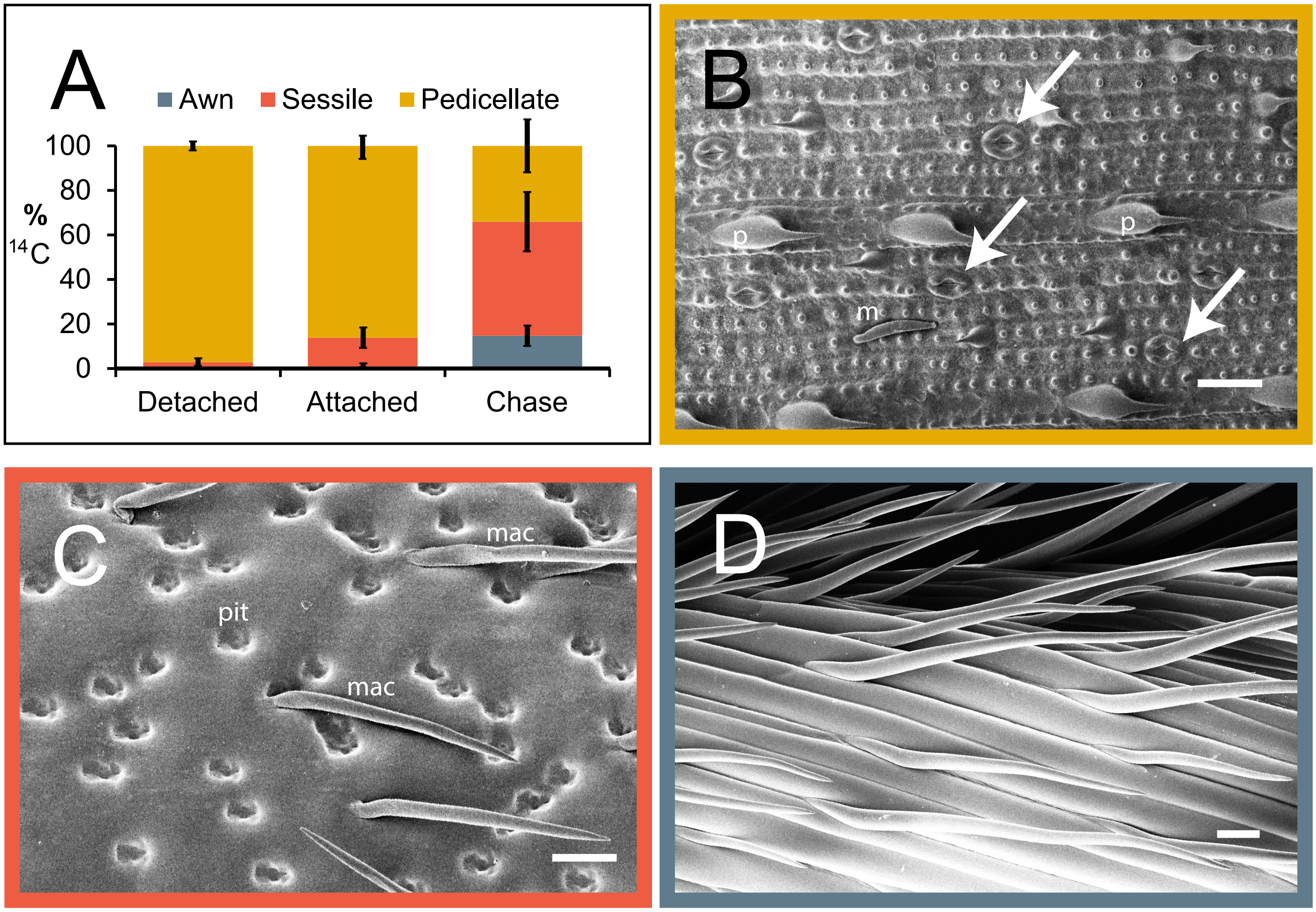

Figure S3. *Themeda triandra*. A. $^{14}C$ results. Percent dpm for each organ after 1 hour exposure to $^{14}C$ with organs removed from the axis (detached), inflorescence intact (attached), or after 24-hour chase. Plot includes mean percentages and standard deviations (n=3). Values are similar after 1 hour, whether organs are attached or lying on filter paper. Change after 24-hour chase suggests movement of $^{14}C$ from pedicellate to sessile spikelet. Awn is largely unlabeled. B. Abaxial epidermis of pedicellate spikelet showing rows of stomata (arrows), bicellular microhairs (m), and prickles (p). C. Abaxial epidermis of sessile spikelet showing no stomata, but large pits (pit) and macrohairs (mac). D. Awn showing no stomata, but long macrohairs. Scale = 50 µm; note that B and C are more highly magnified than D.

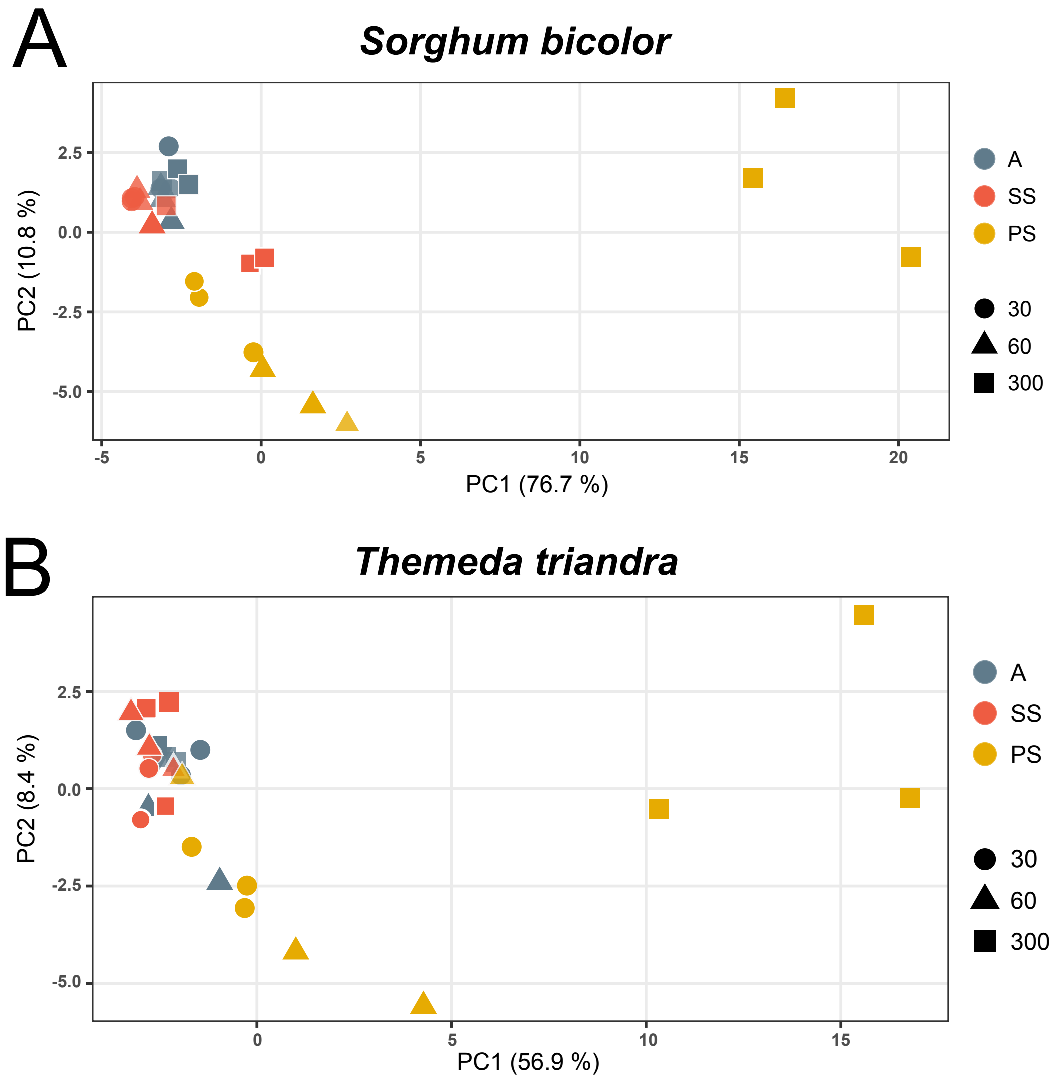

Figure S4. PCA of all isotopologues for $^{13}C$ experiment, showing similarity of results between species. Awn and sessile spikelet are not significantly different for most metabolites, whereas pedicellate spikelet is significantly different from the other two organs. Values at 30 and 60 seconds are not significantly different for most metabolites but 300 second time point is distinct. A. *Sorghum bicolor*. B. *Themeda triandra*.

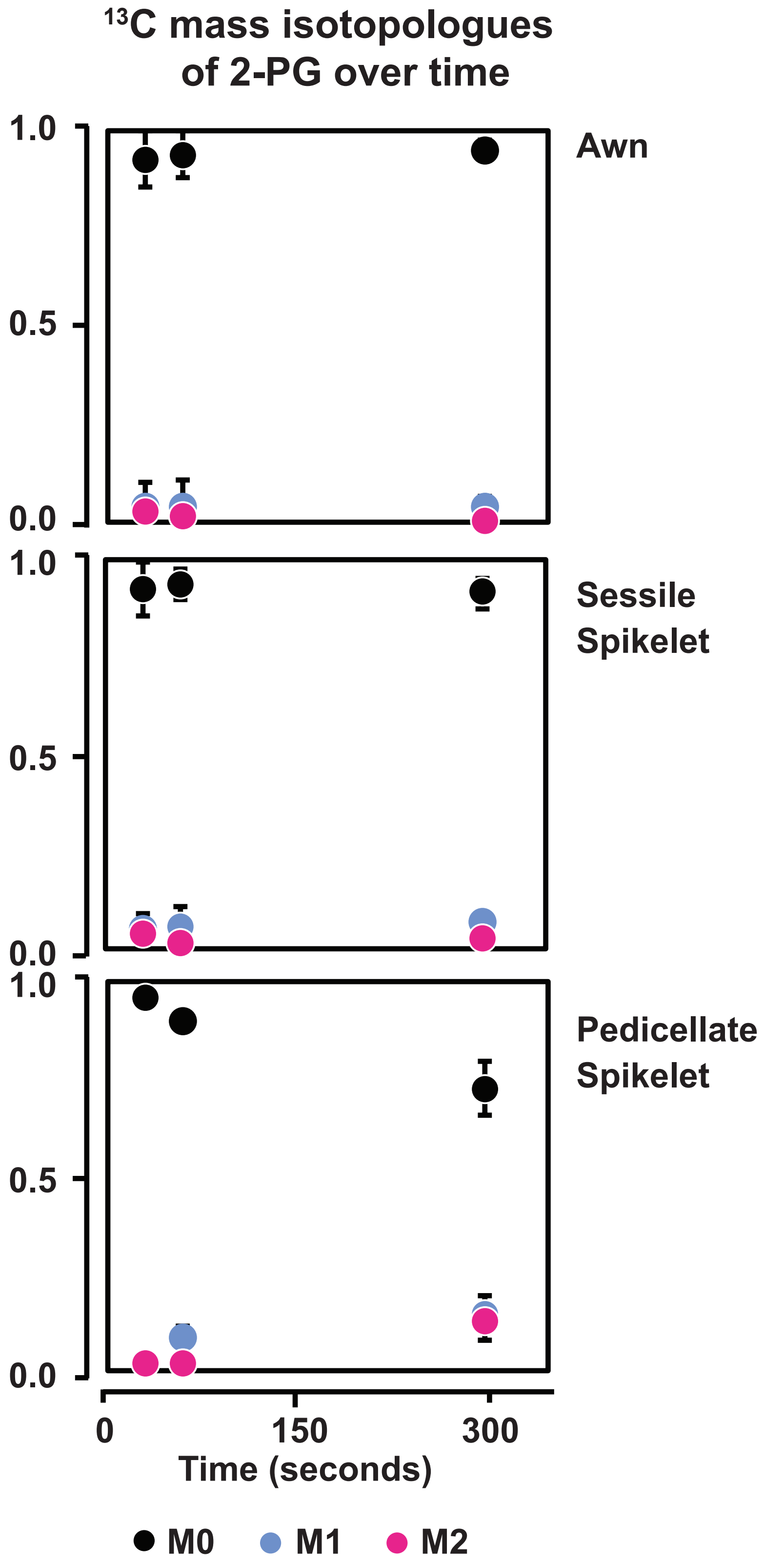

Figure S5. Isotopologues of 2-phosphoglycolate over time. Points are mean percentages, bars are standard deviations (n=3). 2-PG is labeled only in the pedicellate spikelet by the 300 second time point.

**Figure S6.**

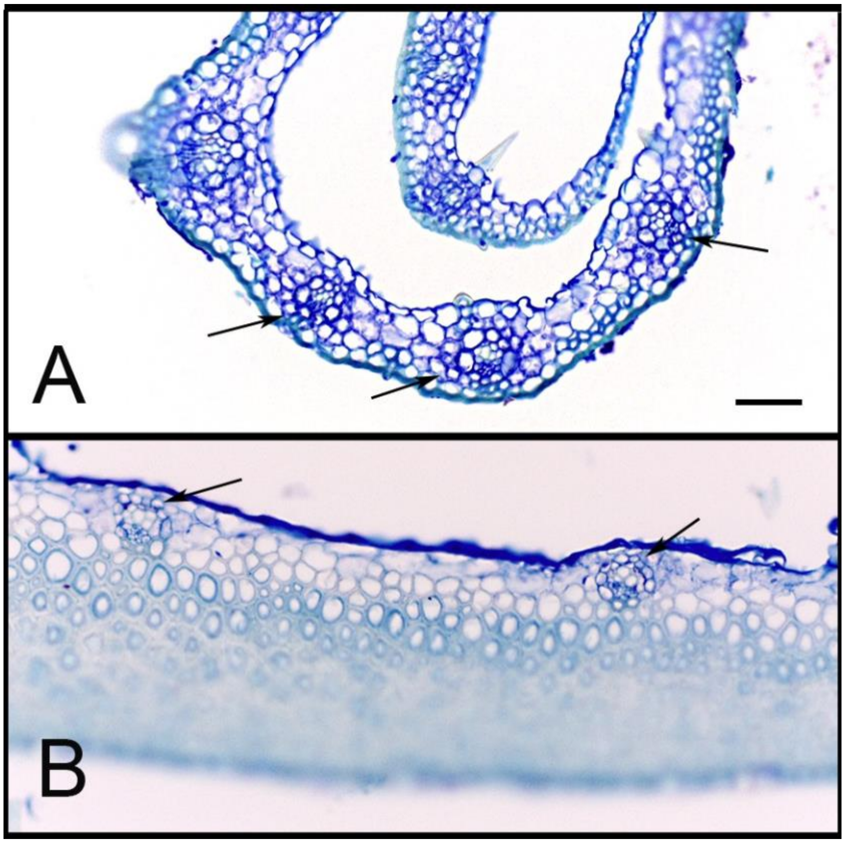

Figure S6. Cross section of glumes of sorghum spikelets, showing vein spacing. A. Pedicellate spikelet. Veins (indicated by arrows) are separated by three to five mesophyll cells, generally more than in the leaf. B. Sessile spikelet. Veins are small and distantly separated. Abaxial side is largely made up of thick-walled cells with no vascularization. Scale = 20 µm.

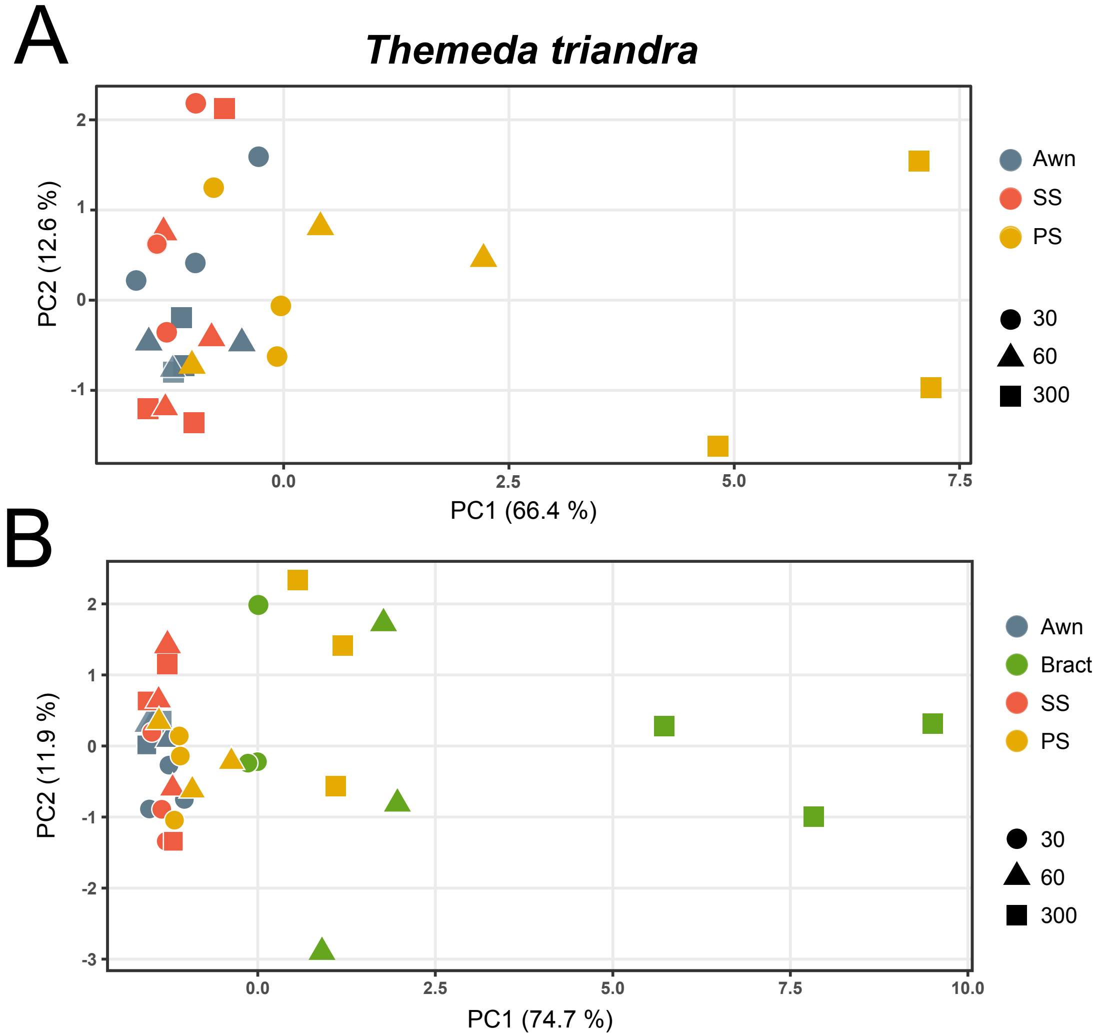

Figure S7. *Themeda triandra*, PCA of $^{13}C$ labeled metabolites, average labeling. Organs are distinguished by color, time points by shape. A. Spikelets and awn only. Values for awn and sessile spikelet are not significantly different at any time point, whereas values for the pedicellate spikelet are significantly different, with the greatest variation in labeling at 5 minutes (300 sec). B. Spikelets, awn, and bract. Values for awn, sessile spikelet and pedicellate spikelet the same as those in A. Awn and sessile spikelet are not significantly different at any time point, whereas values for the pedicellate spikelet are significantly different from awn and sessile spikelet, and bract different from the other three organs, with the greatest variation in labeling at 5 minutes (300 sec). Awn, awn; SS, sessile spikelet; PS, pedicellate spikelet; Bract, bract. 30, values at 30 sec of labeling; 60, values at 60 sec of labeling; 300, values at 300 sec of labeling.

### Figure S8.

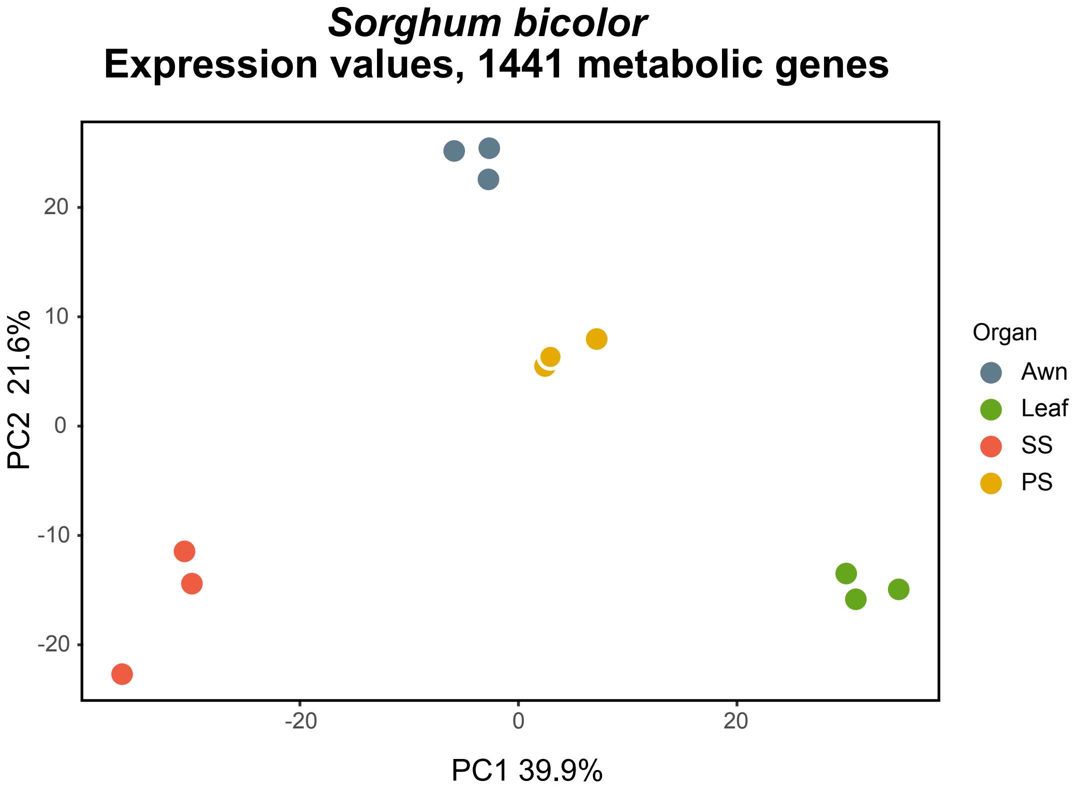

Figure S8. *Sorghum bicolor*, PCA of gene expression data for the 1441 select metabolic genes, showing distinct sets of transcripts for each organ. Values for each replicate experiment are similar. SS, sessile spikelet; PS, pedicellate spikelet.

#### Figure S9.

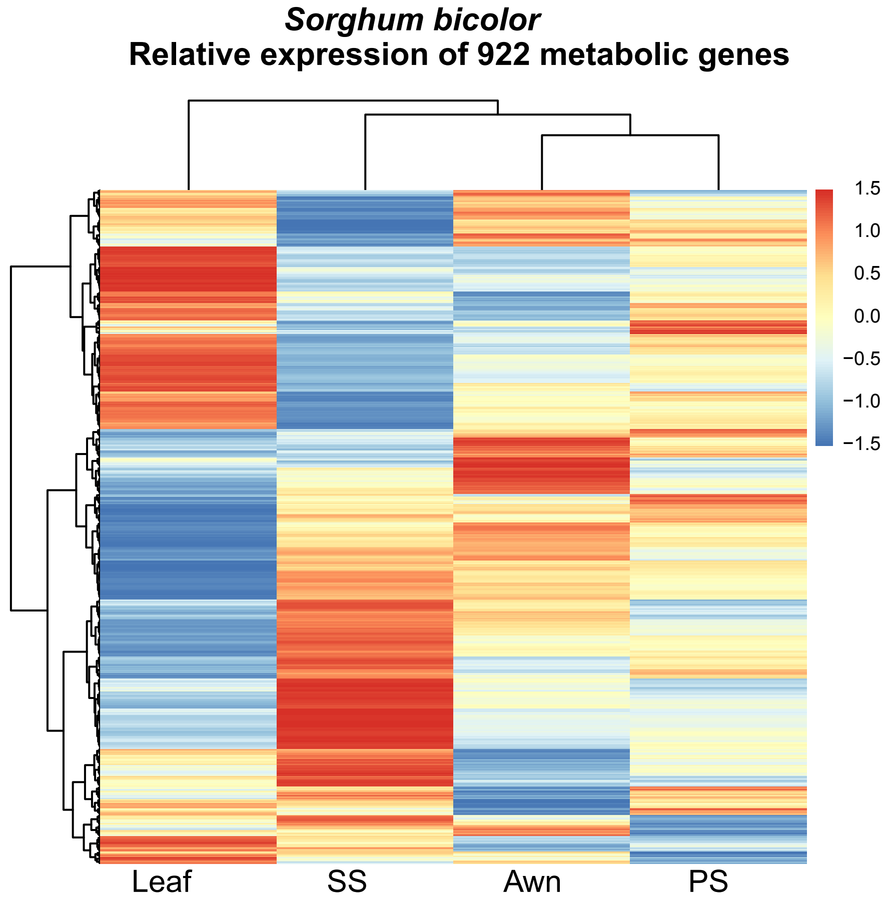

Figure S9. *Sorghum bicolor*, heat map of the 922 differentially expressed metabolic genes, by organ. These are a subset of the 1441 genes depicted in the PCA in Figure S8. Colors indicate relative expression level of genes, normalized by $log_2$. SS, sessile spikelet; PS, pedicellate spikelet.

### Figure S10.

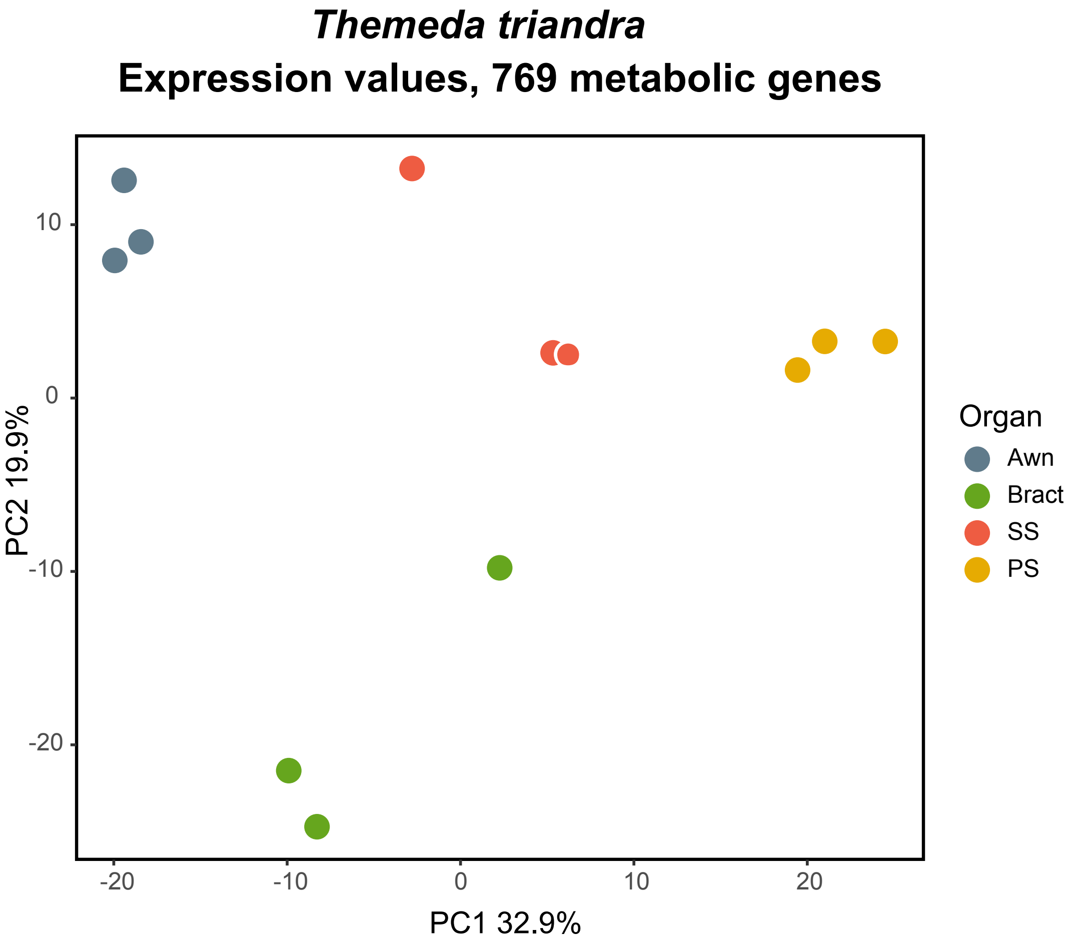

Figure S10. *Themeda triandra*, PCA of gene expression data for the 769 select metabolic genes,, showing distinct sets of transcripts for each organ. Values for each replicate experiments are similar, albeit with outliers for awn and bract. SS, sessile spikelet; PS, pedicellate spikelet.

#### Figure S11.

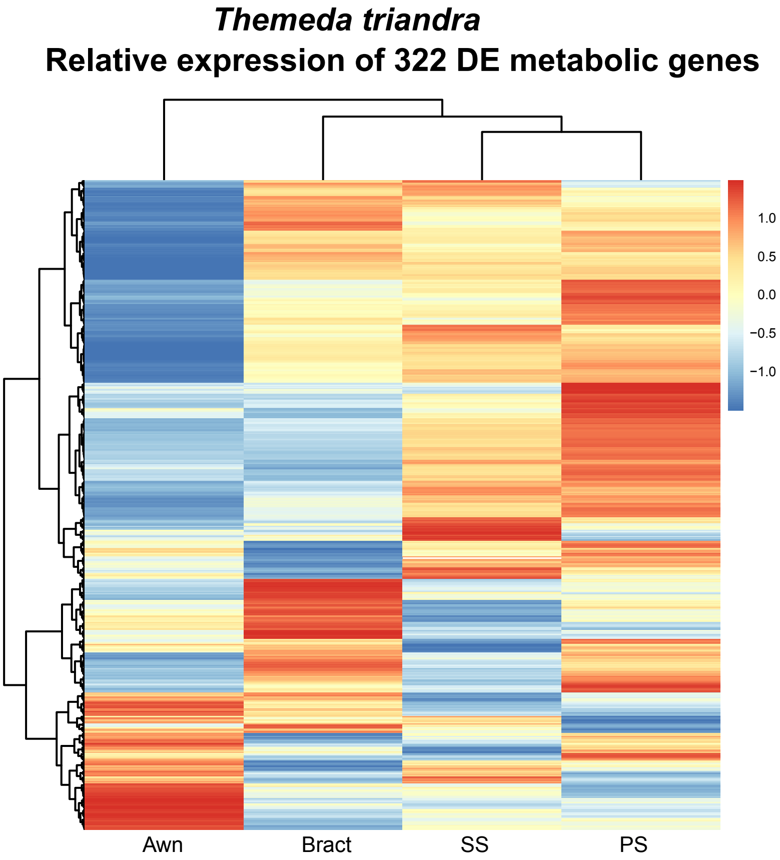

Figure S11. *Themeda triandra*. Heat map of 322 differentially expressed metabolic genes, by organ. These are a subset of the 769 genes depicted in the PCA in Figure S10. Colors reflect scaled z-scores of $log_2$-normalized expression values. SS, sessile spikelet; PS, pedicellate spikelet. Putative biosynthetic pathways indicated as in Figure 5. SS, sessile spikelet; PS, pedicellate spikelet.

Figure S12. *Themeda triandra*. Relative expression of 24 genes encoding biosynthetic enzymes immediately responsible for producing the metabolites labeled with $^{13}C$, a subset extracted from full set of 322 DE metabolic genes in Fig. S11. Colors reflect scaled z-scores of $log_2$-normalized expression values. Labels of genes indicate enzyme name, biochemical process, and subcellular localization. SS, sessile spikelet; PS, pedicellate spikelet; cy, cytosolic localized; ch, chloroplast localized; mi, mitochondrial localized; OA/AA, organic acid/amino acid metabolism.
*transcripts also DE in *Sorghum bicolor*; see Figure 5.
1) ASPAT, aspartate aminotransferase, Sobic.009G149400.1; 2) *NADP−MDH, NADP-malate dehydrogenase, Sobic.007G166300.1; 3) *G1P-AdT, glucose-1-phosphate adenyltransferase Sobic.002G160400.1; 4) *TPI, triose phosphate isomerase, Sobic.003G072300.2; 5) SuSY,_sucrose synthase_Sobic.010G072300.1; 6) ASNS, asparagine synthetase_Sobic.010G110000.1; 7) GAPDH, glyceraldehyde 3-phosphate dehydrogenase, Sobic.006G105900.1; 8) *F16BP aldo, fructose-1,6-bis-phosphate aldolase, Sobic.005G056400.1; 9) F16BPase, fructose-1,6-bisphosphatase, Sobic.001G425400.1; 10) PRK, phosphoribulokinase, Sobic.006G200800.1; 11) *Starch synthase, Sobic.004G238600.1; 12) ASPAT, aspartate aminotransferase, Sobic.004G331700.1; 13) G1P AdT, G1P−adenyltransferase, Sobic.002G088600.2; 14) GAPDH, glyceraldehyde 3-phosphate dehydrogenase, Sobic.009G016700.1; 15) ASPAT, aspartate aminotransferase, Sobic.004G104700.1; 16) UTP-G1P UdT, UTP−G1P uridyltransferase, Sobic.004G013500.1; 17) *SuSY, sucrose phosphate synthase, Sobic.004G068400.1; 18) UTP-G1P UdT, UTP−G1P uridyltransferase, Sobic.010G251200.1; 19) PEPCK, phosphoenol pyruvate carboxykinasae, Sobic.001G432800.1; 20) ASNS, asparagine synthetase, Sobic.001G406800.1; 21) PPI, phosphopentose isomerase, Sobic.005G033000.1; 22) GS, glutamine synthetase, Sobic.001G451500.1; 23) *PGI, phosphoglucoisomerase, Sobic.002G230600.1; 24) *S17BPase, sedoheptulose 1,7-bisphosphatase, Sobic.003G359100.1.

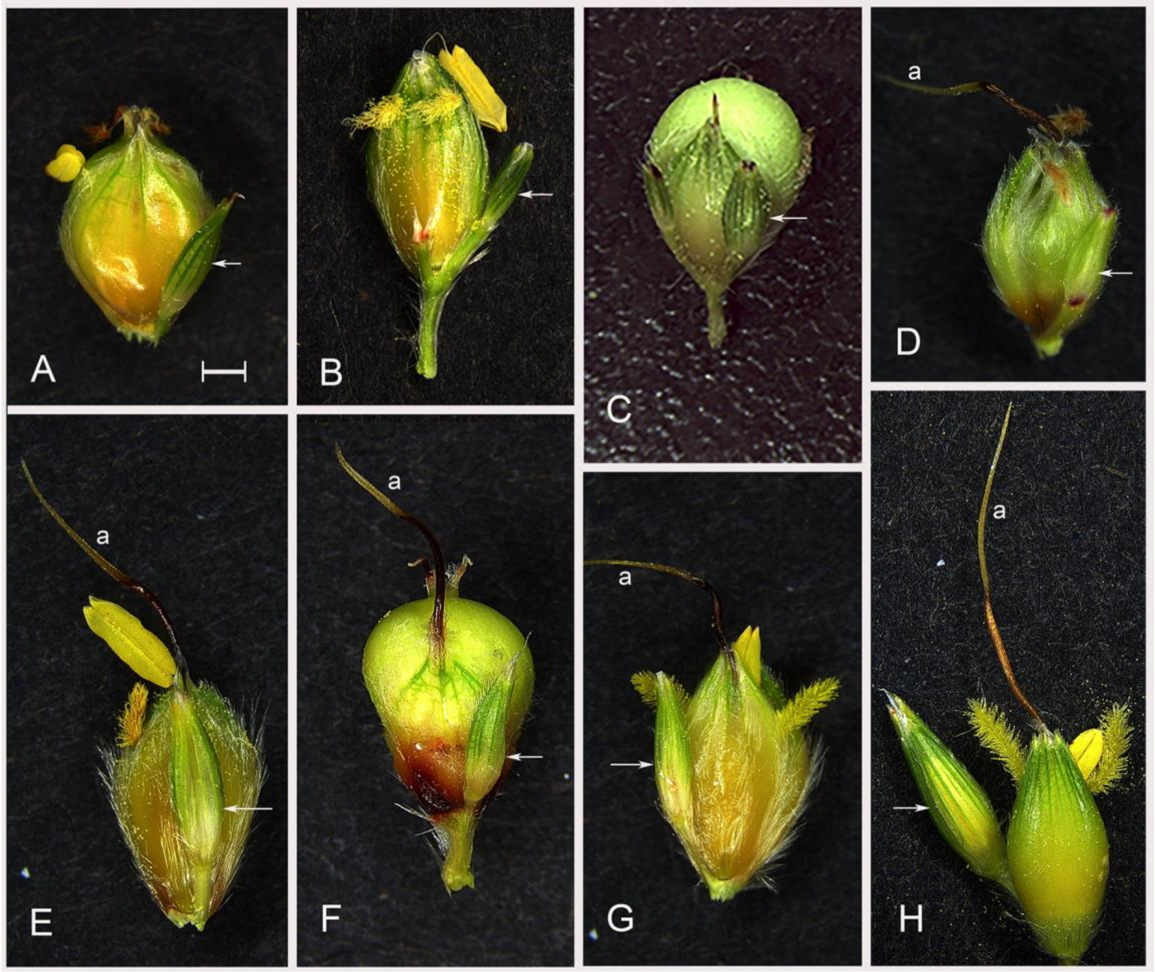

Figure S13. Spikelet pairs from the eight accessions used for spikelet removal experiments. A. SAP-170 (PI 597971). B. BTx623 (PI 564163). C. Combine Hegari (PI 659691). D SAP-257 (656099). E. SAP-51 (655995). F. Jola Nandyal (534021). G. SAP-15 (656014), H. SO85 (PI 534096). Arrow indicates pedicellate spikelet. All spikelets to the same scale. Scale bar = 1 mm.

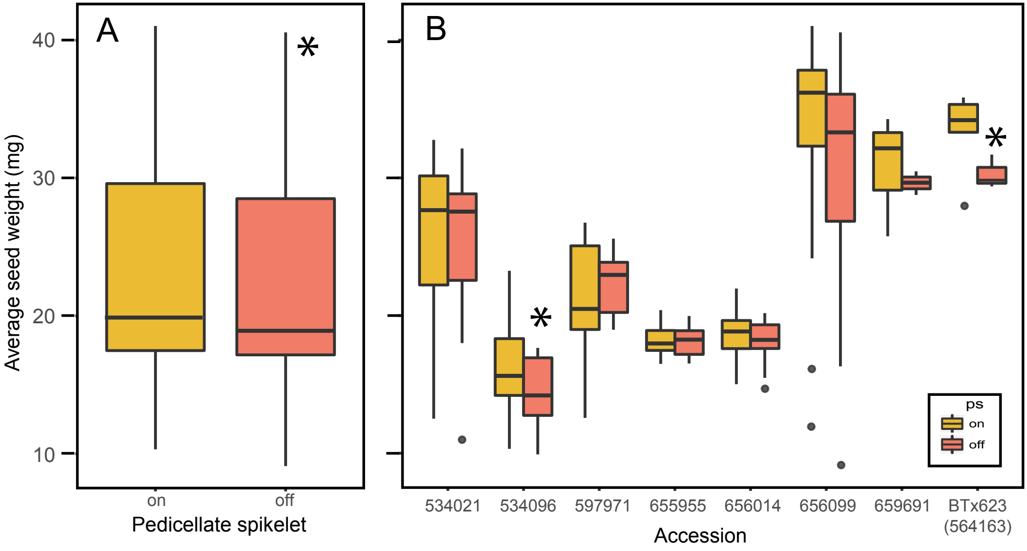

Figure S14. Average seed weight (mg) from inflorescences with pedicellate spikelets untouched (on) and removed (off). A. Combined results of both removal experiments, showing 5.2% reduction in average weight with spikelet removal (p=0.0197). B. Average seed weight for each accession in experiment 1. Representative spikelet pair for each accession shown in Figure S1. * = p <0.05.
